# Converging evidence for differential regulatory control of *APOEε4* on African versus European haplotypes

**DOI:** 10.1101/2021.08.23.457375

**Authors:** Karen Nuytemans, Marina Lipkin, Liyong Wang, Derek Van Booven, Antony J. Griswold, Farid Rajabli, Katrina Celis, Oded Oron, Natalia Hofmann, Sophie Rolati, Catherine Garcia-Serje, Shanshan Zhang, Fulai Jin, Mariana Argenziano, Struan F.A. Grant, Alessandra Chesi, Christopher D. Brown, Juan I. Young, Derek M. Dykxhoorn, Margaret A. Pericak-Vance, Jeffery M. Vance

## Abstract

**INTRODUCTION:** The difference in *APOEε4* risk for Alzheimer disease (AD) between different populations is associated with *APOEε4* local ancestry (LA). We examined LA SNPs with significant frequency differences between African and European/Japanese *APOEε4* haplotypes for areas of differential regulation.

**METHODS:** We performed two enhancer Massively Parallel Reporter Assay (MPRA) approaches, supplemented with single fragment reporter assays. We utilized Capture C analyses to support interactions with the *APOE* promoter.

**RESULTS:** The *TOMM40* intron 2 and 3 region showed increased enhancer activity in the European/Japanese versus African LA haplotypes in astrocytes and microglia. This region overlaps with *APOE* promoter interactions as assessed by Capture C analysis. Single variant analyses pinpoints rs2075650/rs157581, and rs59007384 as functionally different on these haplotypes.

**DISCUSSION:** Both differential regulatory function and Capture C data support an intronic region in *TOMM40* as contributing to the differential *APOE* expression between African and European/Japanese LA.

## 1. Introduction

Alzheimer Disease (AD) is the most common neurodegenerative disorder, affecting up to 8% of those over the age of 65^1^. Currently there are no treatments that effectively prevent disease progression. The most significant genetic risk factor for AD is *APOE*, with the *APOE*ε4 allele inferring both an increased risk for disease and a decrease in age-at-onset^2,3^. Interestingly, the risk effect of this allele varies greatly between populations^4^. East-Asian populations and non-Latino/Hispanic Whites carrying an ε4 allele have a considerable higher risk for AD than African-ancestry populations (East-Asian; ε4/ε4 OR: 11.8-33.1, Non-Latino/Hispanic whites ε4/ε4 OR: 14.9, African Ancestry ε4/ε4 OR: 2.2-5.7)^4-7^. Thus, identification of the mechanism leading to this lower or “protective” risk for AD in the African genome for carriers of *APOE*ε4 would have tremendous potential for therapeutic intervention in non-African ancestry populations.

Though studies in African-Ancestry populations confirmed an increased risk for AD with *APOEε4*, they often showed inconsistent results for the risk effect^7-10^. Rajabli et al. hypothesized that the lower risk for AD and the range of OR could be ascribed to the variable admixture in the African-ancestry populations. Using local ancestry mapping in a large group of admixed individuals (African Americans and Puerto Ricans), the authors confirmed that the ancestry-specific region surrounding the *APOE* gene (“local ancestry, LA”) is directly associated with the difference in the risk effect of AD between populations^11^. A similar finding was subsequently reported by Blue et al^12^. Thus, the global “average ancestry” of the individuals (correlated with ethnic, cultural, and environmental factors) generally used in *APOE* analyses was not by itself associated with *APOEε4* risk. Comparing allele frequencies on the respective *APOEε4* haplotypes of different ancestry (1000 genomes), Rajabli et al. identified variants with significant differences between high risk (Japanese and European) and low risk (African) *APOEε4* haplotypes in the LA surrounding *APOE*^*11*^. None of the variants with differential allele frequency on the different haplotypes were located in any of the coding region of genes within the region surrounding the *APOE* gene, suggesting that the potential protective factor(s) are likely noncoding variants affecting regulatory elements that potentially influence gene expression levels.

The identification of the functional noncoding variant(s) driving association effects is complicated by linkage disequilibrium between variants on a haplotype and the considerable distance that can exist between regulatory elements influencing each other (e.g., enhancers to promoters). The advent of Hi-C chromosomal conformation assays has facilitated the identification of genomic regions with increased cross-regulation within its boundaries, so called topologically associated domains (TADs). These TADs allow for the identification of the likely regulatory region around a gene. Several studies have identified potential modulators of *APOE* expression, including *APOE* promoter variants (rs405509/rs440446)^13-17^, enhancers^18-20^, and CpG island methylation^21-23^ in *APOE*. However, these studies are often limited to candidate regulatory regions in *APOE* (i.e., *APOE* promoter, exon 4) and have not considered variants in the larger TAD surrounding *APOE* (further referred to as ‘TAD variants’ or ‘TV’). Additionally, none of these studies have focused on potential regulatory differences between *APOEε4* ancestral haplotypes.

We hypothesized that the variant contributing to the lower *APOEε4* risk 1) has a differential allele frequency between the high-risk and low-risk *APOEε4* haplotypes and 2) will have an effect on regulation of expression of *APOE* and/or other genes in this region, and as such is potentially contributing to the lower risk (or “protective”) effect seen on the African versus Japanese or European *APOEε4* haplotypes. To evaluate the functional effect of variants with allelic differences between populations, we leveraged several unbiased approaches for identification of regulatory potential for TV surrounding *APOE* using two different massively parallel reporter assays (MPRA) designs, both supported by single variant luciferase reporter assays and Capture C analyses.

## 2. Material and Methods

### 2.1. Creation of MPRA enhancer libraries

The TV variants identified to have significant differential allele frequencies on the different risk *APOEε4* haplotypes peaked in a small region (∼subTAD) surrounding *APOE* (chr19:45375k-45440k)^11^. This lies within a larger well-defined TAD, based on the 3D genome browser data from hippocampus^24^ and Hi-C data from the Jin lab^25^ (**Supplementary Figure 1**). We selected all variants identified through 1000 Genomes in this subTAD region directly surrounding *APOE* that met Bonferroni significance in differential frequency on the African versus European or African versus Japanese *APOEε4* haplotype (Bonferroni adjusted p<0.05, N= 56) and had a greater than 0.01 allele frequency. We will further refer to these significantly different variants as ‘significant TV’ or ‘sTV’. To assess the functional potential of these variants, we designed two different MPRA assays, one using large PCR-based genome fragments and the other using a probe-based design assessing single variants.

#### 1.1.1. MPRA design using large PCR-based fragments

In the first assay, we amplified larger fragments (∼850bp) encompassing the sTV haplotypes using PCR (**Figure 1**). These larger amplicons allow for full inclusion of larger regulatory elements in the fragment, as well as inclusion of multiple variants closely located to each other on the same haplotype permitting inclusion of synergistic signals. This allows the fragments to contain additional TV located on African or European haplotypes that could participate in regulation but did not survive the significance threshold determined by Rajabli et al.^11^ We designed 39 fragments of ∼850bp encompassing a total of 56 sTV; with 11 fragments harboring either two or more sTV. Variant or variant haplotype fragments were amplified in African American heterozygous individuals carrying the variants of interest located in the middle of the fragments (**Supplementary Table 2 and 3**) and cloned into the enhancer reporter vector pGL4.24 (E8421, Promega). Subsequently, a library of 20bp barcodes was cloned into the 3’UTR of the vector library. This final library was used for the transfection experiments. The variant (haplotype) was linked to a unique barcode using the methodology published in Trizzino et al^26^. More information on the library creation, cell culture and sequencing can be found in **Supplementary Material**.

**Figure 1.**
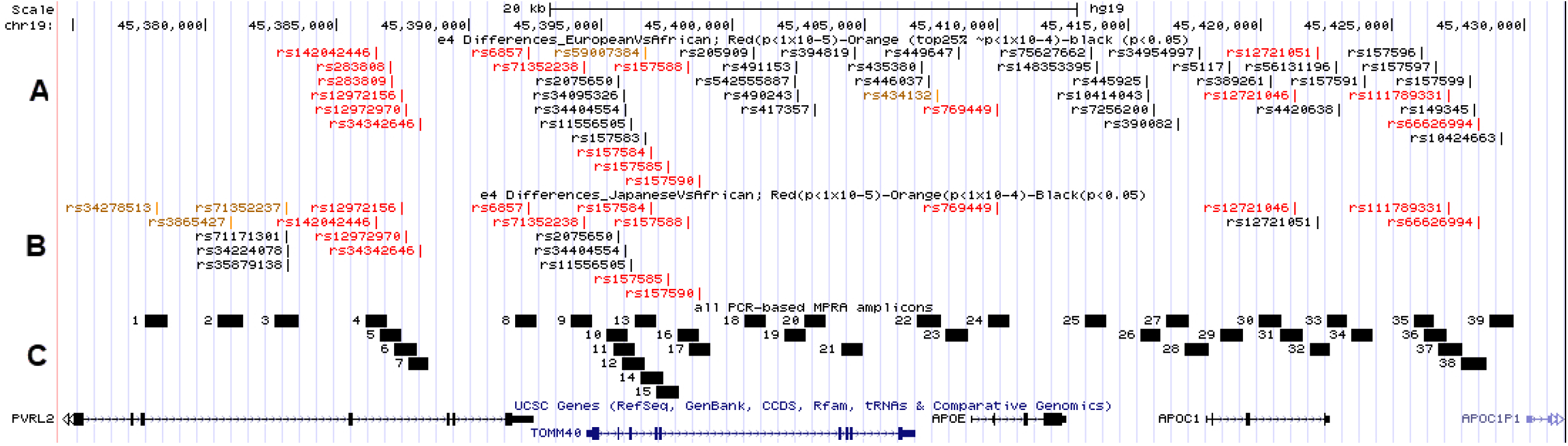
Overview of experimental set-up. A; All variants identified to have differential frequency between European (1000 genomes CEU) vs African (1000 genomes YRI) *APOEe4* haplotypes; color-coded for significance; red; p<1*10-5, orange p<1*10-4 (corresponding to the top 25% results), rest p<0.05. B; All variants identified to have differential frequency between Japanese (1000 genomes JPT) vs African (1000 genomes YRI) *APOEe4* haplotypes; color-coded for significance; red; p<1*10-5, orange p<1*10-4, rest p<0.05. C; All 39 designed amplicons for inclusion in the PCR-based MPRA.

#### 1.1.2. Probe-based MPRA design

In the second assay, we utilized shorter, synthetically made probes. This allows for high-throughput inclusion of variants and single variant assessment but is more limited in the inclusion of adjacent variants since the probe size is restricted to ∼200bp. This approach was utilized to identify regulatory variants as part of a larger design including several AD loci, totaling ∼24,000 variants. For each sTV, 180bp of reference sequence (90bp on either side of variant) was extracted for the “ref” allele fragment. An alternate (“alt”) allele fragment was created by replacing just the position of interest with the alternate allele. The same variants as used for the PCR-based MPRA design were included in this approach, except for the five indel variants, which were excluded for technical reasons in the synthesis. Using these sequences, DNA probes were synthesized (Twist Biosciences) after which 20bp barcodes were attached to these fragments by PCR using a barcoded primer library. Combined fragments were cloned into a previously published vector^27^ and sequenced for variant to barcode link determination. Once validated, the reporter gene with a minimal promoter was cloned in between the fragments and barcodes. More information on the library creation, cell culture and sequencing can be found in **Supplementary Material**.

### 2.2. Luciferase reporter assays

MPRA results were supplemented by single variant/haplotype reporter assays using the Dual-Glo® Luciferase System (Promega Corporate, Madison, WI, USA) for confirmation or when MPRA assays failed to generate sufficient data for analyses. For each region, fragments of both haplotypes were amplified using the same primers as the PCR-based MPRA design (**Supplementary Table 2 and 3**) and cloned separately into the pGL4.24 plasmid to test for enhancer activity. Additionally, we amplified a smaller size fragment 9 into pGL4.10 to test for promoter activity.

### 2.3. Statistics

#### 2.3.1. MPRA

We performed the exact binomial test to find whether the proportion of the two alleles in the expressed material (RNA-Alt, RNA-Ref) is significantly different from the expected ratio (observed in MPRA library; DNA-Alt, DNA-Ref)^28^. We implemented analysis using in-house R scripts. Raw data can be found in **Supplementary Table 1**. For the PCR-based MPRA, we required at least one out of two replicates to survive multiple testing, with the other meeting p<0.05 for confirmation. For the probe-based MPRA, we required at least one out of three replicates to survive multiple testing, with a second replicate also surviving multiple testing, or both replicates meeting p<0.05, for confirmation.

#### 2.3.2. Luciferase reporter assays

Results obtained from three independent experiments were evaluated by *t*-test. A p-value of <0.05 was considered significant in these analyses.

### 2.4. Capture C data

We performed high resolution (i.e., using the 4 bp-cutter DpnII, as opposed to the more commonly used 6 bp-cutter HindIII) genome-wide promoter-focused Capture C analyses on NPC, astrocyte, microglia and neuronal cell lines, following the previously described protocol^29^. NPC and neurons were derived from two iPSC lines from control individuals (CHOPWT10 and CHOPWT14) as described in Su et al^30^.

Libraries were generated in triplicates for each cell type and donor and paired-end sequenced on the Illumina Novaseq 6000 platform (51bp read length) at the Center for Spatial and Functional Genomics at Children’s Hospital of Philadelphia (CHOP). Reads were pre-processed using the HiCUP pipeline (v0.5.9^31^), with the bowtie aligner and hg19 as the reference genome. Significant interactions at 4-DpnII fragment resolution were called using CHiCAGO^32^.

### 2.5. In-silico annotation

For each variant included in the probe-based MPRA design, we evaluated their presence in ChIP-seq transcription factor binding sites (TFBS) from ENCODE, in putative enhancer or promoter regions (based on RoadMap Epigenome data) and in predicted TFBS (using the RegulomeDB Jaspar query and MotifBreakR). Although ENCODE^33^ has ChIP-seq data for many transcription factors in many cell types, data on transcription factors in brain cell types is very limited; neural cell (SMC3, CTCF, EP300, MXI1, RAD21), astrocytes (CTCF), SH-SY5Y (GATA2, GATA3), and NPC (CTCF, EZH2). PrimaryHMM results from RoadMap Epigenome project^34^ of brain regions (hippocampus, dorsolateral prefrontal cortex, singular and angulate gyrus, substantia nigra, and anterior caudate), as well as adrenal gland and liver cells were evaluated. Jaspar^35^ uses position frequency matrices (PFM) to evaluate TFBS. MotifbreakR^36^ was used to further evaluate the effects of variants on transcription factor binding motifs. We used the HOmo sapiens COmprehensive MOdel COllection (HOCOMOCO^37^) for transcription factor binding site identification with a p-value significance threshold of p<5×10^−5^. Further, we evaluated eQTL status of the variants in brain or non-brain tissues using GTEx. Lastly, PsychENCODE H3K27ac peak analysis was used to identify putative enhancer regions based on prefrontal cortex tissue^38^.

## 3. Results

The initial analysis using 1000 Genomes data of *APOEε4* carriers suggested that there were 56 sTV in the *APOE* subTAD (**Table 1**). Overall results on enhancer activity - assayed in representatives of microglia, astrocytes, and neurons - across all methods can be found in the **overview of Table 1** and are visualized in **Figure 2**. Detailed results can be found in **Supplementary Table 1 and Material & Methods**.

**Figure 2.**
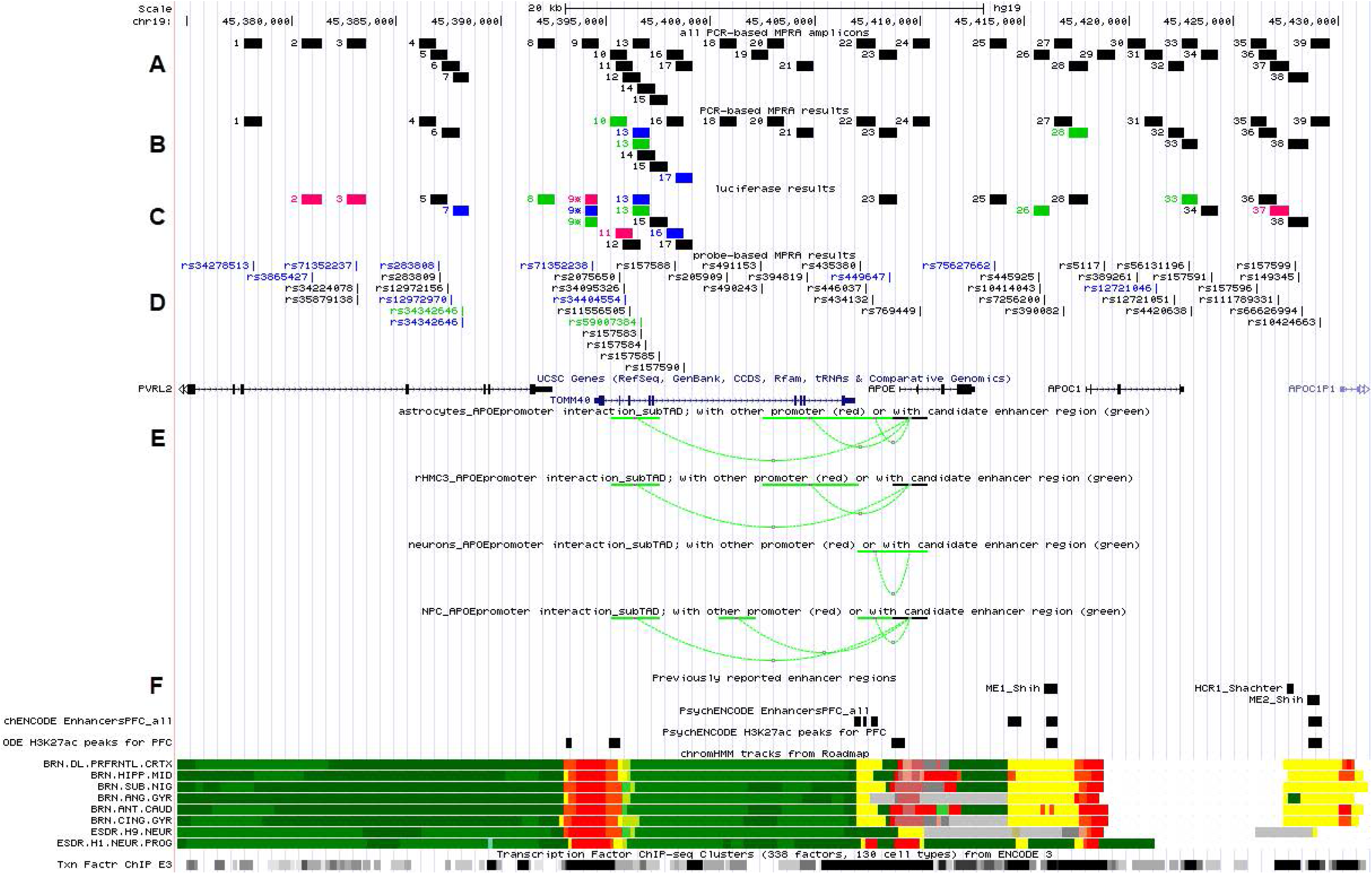
Overview of results and regulatory evidence in the region of interest. A; All 39 designed amplicons for inclusion in the PCR-based MPRA. B; Representation of results of successful fragments in the PCR-based MPRA C; Representation of results of successful fragments in the luciferase experiments. D; Representation of results of successful variant analyses in the probe-based MPRA. Blue; astrocytes. Green; microglia. Pink; neurons. E; Significant interactions of *APOE* promoter within the subTAD based on the promoter Capture C data at 4-DpnII fragment resolution; interactions between promoter and candidate regulatory element (green), interaction between promoters (red). F; Available in-silico annotation data; including previously reported enhancers, enhancers and H3K27ac marks identified in prefrontal cortex from the PsychENCODE project, chromHMM tracks from the RoadMap Epigenome Project (primary HMM) and Transcription factor ChIP-Seq cluster data from ENCODE freeze 3; using data from 338 factors in 130 cell types.

Specifically, we identified a significant regulatory region in microglia and astrocytes supported by several levels of evidence. We identified differential activity in microglia and astrocytes between sTV haplotypes for fragments 10 and 13, both located in the first introns of *TOMM40 (***Table 2**). Higher enhancer activity is observed for the European/Japanese *APOEε4* haplotype in fragment 10 (containing two sTV alleles: rs2075650-G and rs34404554-G) and the European *APOEε4* haplotype in fragment 13 (containing two sTV alleles: rs59007384-T and rs157583-G) (**Table 2**). Haplotype fragment 13 showed consistently higher activity driven by the sTV rs59007384 European allele T in the PCR-based MPRA and luciferase assays (both glia cell lines), and the probe-based analyses (in microglia), supporting the T allele variant as the driving signal in fragment 13. Besides the two sTV SNPs (rs2075650 and rs34404554), European/Japanese and African haplotypes in the PCR-based fragment 10 also contained two additional TV that did not meet Bonferroni significance in the local ancestry comparison (European/Japanese haplotype; rs157580-G and rs157581-T) (**Table 2**). Luciferase analyses of overlapping fragment 11, centering on sTV rs34095326 and rs34404554, did not support differential activity driven by these variants in microglia and astrocytes, suggesting that rs2075650 is the driving factor for the regulatory activity seen with fragment 10. Although 2 out of 3 replicates showed increased enhancer activity for the European/Japanese sTV rs2075650-G allele, this effect could not be replicated in the probe-based analyses (**Supplementary Table 1**). Rs2075650 and rs157581 are both predicted by the JASPAR transcription factor binding database to be located in transcription factor binding sites and are located within 100bp of each other potentially indicating a synergistic effect of these two TV that was missed in the probe-based single variant MPRA. Additionally, according to MotifbreakR, both rs2075650 and rs59007384 are predicted to significantly affect binding sites for SP1 and PLAG, respectively.

Promoter Capture C data in microglia, astrocytes, and NPC consistently show physical interaction of the region encompassing fragments 10-14 with the *APOE* promoter; within subTAD interaction are shown in **Figure 2**. Additional available *in-silico* annotation data supports a regulatory function of this region, with RoadMap Epigenome chromatin mark analyses indicating enhancer activity in the region overlapping with fragments 10-12 (*TOMM40* introns 2-3) in AD-relevant brain regions (hippocampus, frontal cortex), and fragment 13 (*TOMM40* intron 4) in other brain regions (e.g., cingulate gyrus, substantia nigra). Additionally, H3K27ac peaks from prefrontal cortex tissue in PsychENCODE support active regulatory activity in the *TOMM40* introns 1-2 region (overlapping fragments 10-11). No evidence for effect on expression of *APOE* (or any other gene) was observed in GTEx for these variants in the brain.

Although additional chromatin interactions of genomic regions within the subTAD with promoters or candidate regulatory elements outside the subTAD were observed (**Supplementary Figure 2**), these did not overlap with any variants identified to have a significant frequency difference between the European/Japanese and African *APOEε4* haplotypes.

## 4. Discussion

Recently, using single nuclei RNAseq on frozen frontal cortex, we found that *APOEε4* homozygotes lying on a European local ancestry had significantly higher expression of *APOEε4* than *APOEε4* homozygotes lying on an African local ancestry^39^, suggesting a significant difference in the regulatory composition of this regions between ancestries. This supports the importance of determining potential regulatory variants differing in the regulatory architecture between local ancestries. In the brain, *APOE* is primarily produced by glial cells (i.e. microglia and astrocytes), but can also be found to be expressed in neurons, especially under stress conditions^40,41^. Previous descriptions of the regulatory control of the *APOE* gene have suggested it is complex, with different promoter variants identified as well as many regulatory elements, such as Multi-enhancer 1 and 2 (ME1 or 2)^18,19^ and a brain control region (BCR)^20^ having been shown to influence *APOE* expression. However, none of these elements have been studied in the context of local ancestry. Here we report on the analyses of differential enhancer activity of variants significantly different on the African versus European or Japanese local ancestry in the immediate vicinity of *APOE*; including ME1 and 2, overlapping with parts of fragments 26-27 and 39 respectively, but not BCR, as it was located outside the subTAD region with sTV convergence.

Our data suggest there is a major cluster of variants with regulatory potential lying in introns 2-3 of *TOMM40* for microglia and astrocytes. As ApoE is mostly produced in glial cells, the convergence of both differential enhancer activity and *APOE* promoter-focused Capture C data in microglia and astrocytes, but not neurons, strongly supports the importance of this *TOMM40* intronic region in *APOE* regulation in the brain. Probe-based analyses further pinpointed the effect of the higher activity on European haplotypes that is likely due to a few variants (rs2075650/rs157581 and rs59007384). Of these, rs2075650/rs157581 were also significantly different in frequency on Japanese versus African *APOEε4* haplotypes. Interestingly, rs2075650 was predicted to affect a binding site for transcription factor SP1, which was previously identified to be dysregulated in AD^42,43^. Taken together with stronger in-silico data indicating enhancer activity in this region, all the above data support a role of the rs2075650/rs157581 variants in mediating the differences in risk seen for *APOEε4* in European/Japanese versus African ancestries.

Interestingly, a previous study identified higher enhancer activity of the African haplotype acting upon the *APOE* promoter in neuronal cell lines^44^ similar to what we have seen for fragment 11 (rs34404554) though not consistently across methods and cell types. However, the effect of this variant was not or was incompletely tested in microglia and astrocytes in this earlier study. Analyses of the haplotypes used in their study indicated the ‘alternate’ haplotype in the enhancer does not represent the most common haplotype on African *APOEε4* while the promoter haplotypes are a mixture of European/African alleles, complicating comparisons of results. The likely driving variants identified here (rs2075650/rs157581 and rs59007384) have been studied before in several settings, i.e. disease risk, ApoE levels in brain and CSF, or Aβ1-42 levels in CSF^45-47^, though never while also considering *APOE* status and/or full local ancestry haplotypes; again complicating direct comparisons of results.

Previously, other variants in the proximity of *APOE* have been reported to potentially regulate *APOE* expression; most commonly promoter variants rs449647 (“-491A>T”) and rs405509 (“-291G>T”)^13-17^. Although both variants have been reported to significantly influence overall *APOE* expression, and rs405509 was even suggested to be potentially underlying the risk difference for AD between Europeans and Koreans^14^, rs405509 was not significantly different in frequency between European or Japanese and African *APOEε4* haplotypes^11^ and rs449647 did not consistently affect expression in the current analyses (PCR-based MPRA fragment 23 or probe-based MPRA). Therefore, these variants are unlikely to explain the risk difference between African and European/Japanese *APOEε4*. Interestingly, none of the above-described promoter variants nor the potential enhancer activity-affecting variants reported here were identified as eQTLs for *APOE* in the brain in GTEx. In contrast, those variants that were associated with differences in *APOE* expression (**Table 1**), did not show differential enhancer activity in the analyses presented here. These data would suggest that the variants identified as eQTL in GTEx may be tagging a functional SNP currently not picked up in the GTEx analyses. Alleles more common on the African ancestral background are significantly less common in the GTEx dataset, which could explain a loss of power to identify eQTL signals.

Given its close proximity, *TOMM40* and more specifically its poly T repeat SNP rs10524523, has also been implicated in risk for AD^48^. This repeat was not included in the analyses here as we have previously shown that the frequency of this repeat was not observed to be significantly different between African and European *APOEε4* haplotypes, though it might tag protective variants on *APOEε3* haplotypes carrying the ‘very long’ size of the repeat^49^.

We acknowledge that additional significant variants^11^ and chromatin interactions were observed outside the subTAD region (**Supplementary Figure 2**). These however did not overlap with one another and, as such, had a reduced priority for further analyses. We further recognize that the sequence differences between the two *APOEε4* backgrounds could also influence methylation activity in this region. Although differences in methylation between *APOEε4* and the other *APOE* statuses has been studied^21-23,50^, the effect of different ancestral backgrounds has not been fully assessed^51^. Preliminary in-house comparison of CpG methylation in the subTAD on African versus European *APOEε4* haplotypes identified one methylation site with lower methylation levels on European versus African background at the 5’end of fragment 10 (unpublished data). This position does not overlap with any sTVs. These data would again support a regulatory role of this region affected by local ancestry, seemingly independent of sequence variation itself. The effect of methylation on enhancers can be both activating or repressing^52^ so further follow-up would be needed to assess the effect of the methylation in this region on *APOE* expression.

Overall, our use of multiple complementary study designs made it possible to identify clusters of differential enhancer activity potentially driving *APOE* regulation. We acknowledge each design has advantages and pitfalls. Although the probe-based design allows for much easier, high-throughput inclusion of variants and assessment of singular variants, the PCR-based MPRA provides insight in potential synergistic effects of variants on joint haplotypes. Practically, the one-step incorporation of fragment and barcode to the vector in the probe-based MPRA proved to be significantly more efficient at identifying unique barcodes than the two-step incorporation in the PCR-based MPRA.

Finally, our results identify a major locus driving differential enhancer activity between African and European/Japanese *APOEε4* haplotypes in AD-relevant cell types are an important step forward in understanding the differential risk between these *APOEε4* haplotypes. It has long been a matter of debate as to whether increasing or decreasing *APOEε4* would have therapeutic effects^53^. Further evaluation of the regulatory elements identified here will help provide insight in the complex regulation of *APOE* and pinpoint potential efficacious therapies.

## 5. Acknowledgements

We thank Dr. Ryan Tewhey for consulting on the probe-based MPRA library creation. This research was supported by the National Institute on Aging (AG059018 – PI JV; KN, ML, LW, AG, KC, OO, JY, DD-, AG054074 – PI MP; LW, AG, NH, SR, DD, CS, FR, DB, JV-, AG072547 – PI MP; JV-, AG057659 – PI MP, HG009658-PI FJ; SZ-, AG057516 – PI SG; MA, AC-) as well as Alzheimer Association (ZEN-19-591585-PI JV; KN, ML, LW, KC, OO, JY, DD) and BrightFocus (A2018425S – PI JV; KN, LW, AG, FR, JY).

## 8. Supplemental data

**Supplement Table 1. Raw data and p-values for all experiments**

**Supplement Table 2. Full list of TV overlapping with tested fragments**.

**Supplement Table 3. List of primers used in MPRA analyses**.

**Supplementary Figure 1. Representation of 2MB local ancestry and TAD structure surrounding *APOE***.

A; Bonferroni correct p-values for pairwise comparisons of allele frequencies in 1000 genome data between (top) CEU versus YRI, and (bottom) JPT versus YRI populations^11^.

B; Genome-wide Hi-C chromatin interaction analyses at 40k resolution^25^ across the 2MB region in available relevant cell types/tissues.

**Supplementary Figure 2. All promoter Capture C data within *APOE* subTAD region**.

Significant interactions at 4-DpnII fragment resolution between promoter and candidate regulatory elements (green) or interaction between promoters (red). Red boxes display gene name of promoter not in subTAD. Green boxes display genomic positions of candidate regulatory elements not in subTAD. No non-*APOE* promoter interactions were observed in the microglia lines.

## Supplemental Material

Detailed information on methods and results.

## Browser

https://genome.ucsc.edu/hg19browser_Nuytemans

https://genome.ucsc.edu/hg38browser_Nuytemans

---

| Fragment | centered TV SNPs | Location | Allele most common in ε4 |  |
| --- | --- | --- | --- | --- |
|  |  |  | AFR | EUR/JPT |
| 1 | rs34278513 | intron PVRL2 | C | T |
| 2 | rs3865427 | intron PVRL2 | C | A |
| 3 | rs71352237* | intron PVRL2 | T | C |
|  | rs71171301* | intron PVRL2 | - | C |
|  | rs34224078* | intron PVRL2 | A | G |
|  | rs35879138* | intron PVRL2 | T | A |
| 4 | rs142042446 | intron PVRL2 | - | TAA |
| 5 | rs283808 | intron PVRL2 | C | A |
|  | rs283809 | intron PVRL2 | G | A |
| 6 | rs12972156 | intron PVRL2 | C | G |
|  | rs12972970 | intron PVRL2 | G | A |
| 7 | rs34342646 | intron PVRL2 | G | A |
| 8 | rs6857 | 3'UTR PVRL2 | C | T |
| 9 | rs71352238 | Up TOMM40 | T | C |
| 10 | rs2075650* | intron TOMM40 | A | G |
| 11 | rs34095326 | intron TOMM40 | G | A |
|  | rs34404554 | intron TOMM40 | C | G |
| 12 | rs11556505 | exonic TOMM40 | C | T |
| 13 | rs59007384* | intron TOMM40 | G | T |
|  | rs157583 | intron TOMM40 | T | G |
| 14 | rs157584 | intron TOMM40 | C | T |
| 15 | rs157585 | intron TOMM40 | C | A |
| 16 | rs157588 | intron TOMM40 | T | C |
| 17 | rs157590 | intron TOMM40 | C | A |
| 18 | rs205909 | intron TOMM40 | G | T |
| 19 | rs542555887 | intron TOMM40 | T | - |
|  | rs491153 | intron TOMM40 | T | C |
|  | rs490243 | intron TOMM40 | T | C |
| 20 | rs417357 | intron TOMM40 | T | C |
| 21 | rs394819 | intron TOMM40 | T | G |
| 22 | rs435380 | up APOE | A | G |
|  | rs446037 | up APOE | T | G |
|  | rs434132 | up APOE | G | C |
| 23 | rs449647 | up APOE | T | A |
| 24 | rs769449 | intron APOE | G | A |
| 25 | rs75627662 | intergenic | C | T |
|  | rs148353395* | intergenic | - | del |
| 26 | rs445925 | up APOC1 | A | G |
|  | rs10414043* | up APOC1 | G | A |
|  | rs7256200* | up APOC1 | G | T |
| 27 | rs390082 | up APOC1 | G | T |
| 28 | 34954997 =rs1156882 | 5' UTR APOC1 | - | CGTT |
| 29 | rs5117* | intron APOC1 | T | C |
| 30 | rs389261 | intron APOC1 | A | G |
| 31 | rs12721046 | intron APOC1 | G | A |
| 32 | rs12721051 | intron APOC1 | C | G |
| 33 | rs56131196* | down APOC1 | G | A |
|  | rs4420638* | down APOC1 | A | G |
| 34 | rs157591 | intergenic | A | G |
| 35 | rs157596 | intergenic | T | C |
| 36 | rs157597 | intergenic | C | A |
| 37 | rs111789331 | intergenic | T | A |
| 38 | rs157599 | intergenic | G | A |
|  | rs149345 | up APOC1P1 | G | T |
|  | rs66626994 | up APOC1P1 | G | A |
| 39 | rs10424663 | up APOC1P1 | A | G |

NA = not assessed/low coverage/failed. Up = upstream. Down = downstream. EUR: E
\* = frequency different general population versus APOEε4 haplotypes

| Enhancer activity (highest activity, cell type) |  |  |
| --- | --- | --- |
| PCR-based MPRA | Luciferase | Probe-based MPRA |
| Negative | NA | JPT astrocytes |
| NA | JPT neurons | JPT astrocytes |
| NA | JPT neurons | EUR astrocytes<br>NA<br>negative<br>negative |
| Negative | NA | NA |
| NA | negative | AFR astrocytes<br>negative |
| negative | NA | negative<br>AFR astrocytes |
| NA | EUR astrocytes | AFR microglia, astrocytes |
| NA | AFR microglia | - |
| NA | NA | AFR astrocytes |
| EUR microglia | NA | negative |
| NA | AFR neurons | negative<br>AFR astrocytes |
| NA | negative | negative |
| EUR microglia, astrocytes | EUR microglia | EUR microglia<br>negative |
| negative | NA | negative |
| NA | negative | negative |
| NA | AFR astrocytes | negative |
| AFR astrocytes | negative | negative |
| negative | NA | negative |
| NA | NA | NA<br>negative<br>negative |
| negative | NA | NA |
| negative | NA | negative |
| negative | NA | negative<br>negative<br>negative |
| NA | negative | AFR astrocytes |
| negative | NA | negative |
| NA | negative | EUR astrocytes<br>NA |
| NA | EUR microglia | negative<br>negative<br>negative |
| negative | NA | negative |
| AFR microglia | negative | NA |
| NA | NA | negative |
| NA | NA | negative |
| NA | NA | EUR astrocytes |
| NA | NA | negative |
| negative | negative | negative<br>negative |
| NA | negative | negative |
| negative | NA | negative |
| NA | negative | NA |
| NA | AFR neurons | negative |
| NA | negative | negative<br>negative<br>negative |
| negative | NA | negative |

European, AFR: African, JPT: Japanese

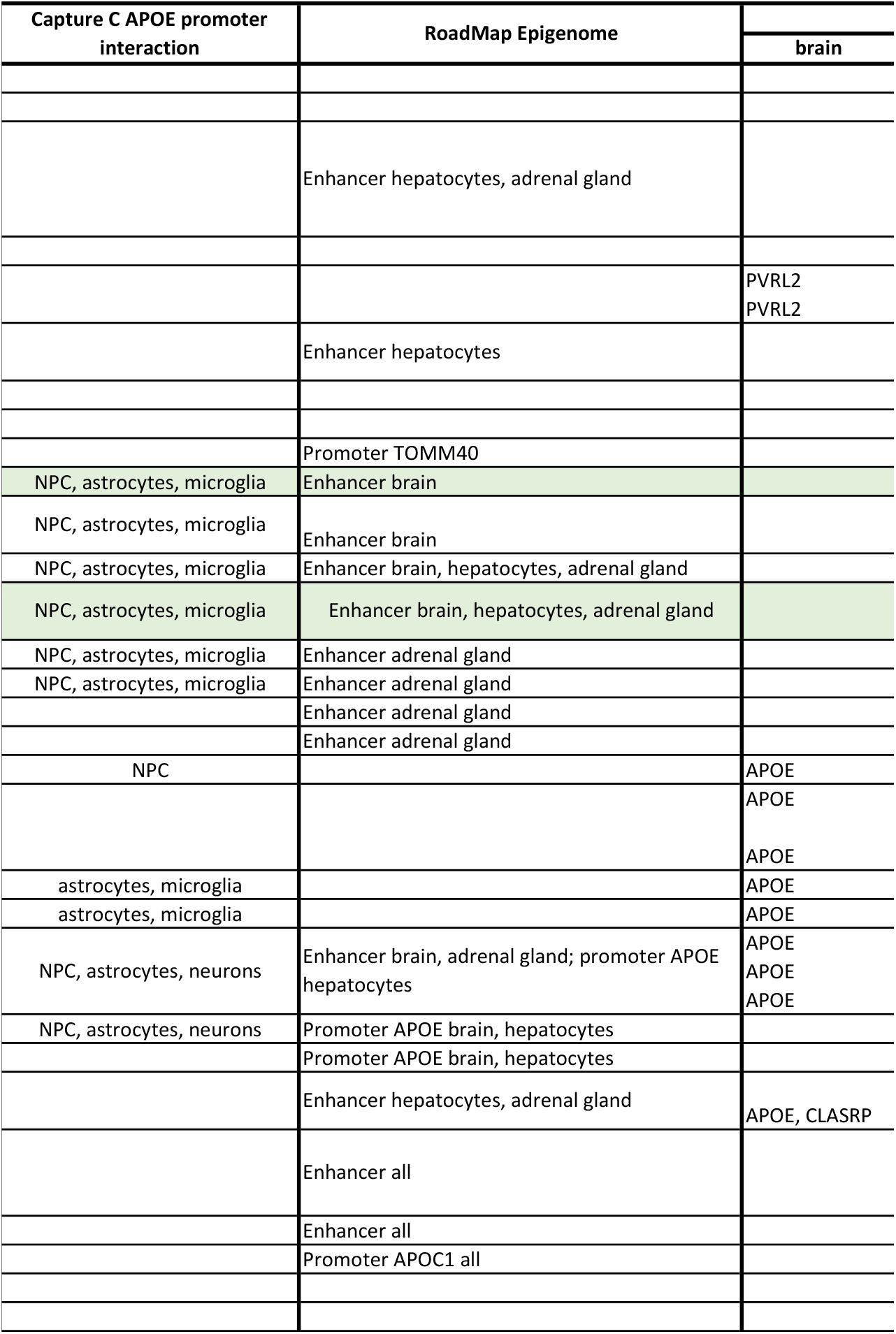

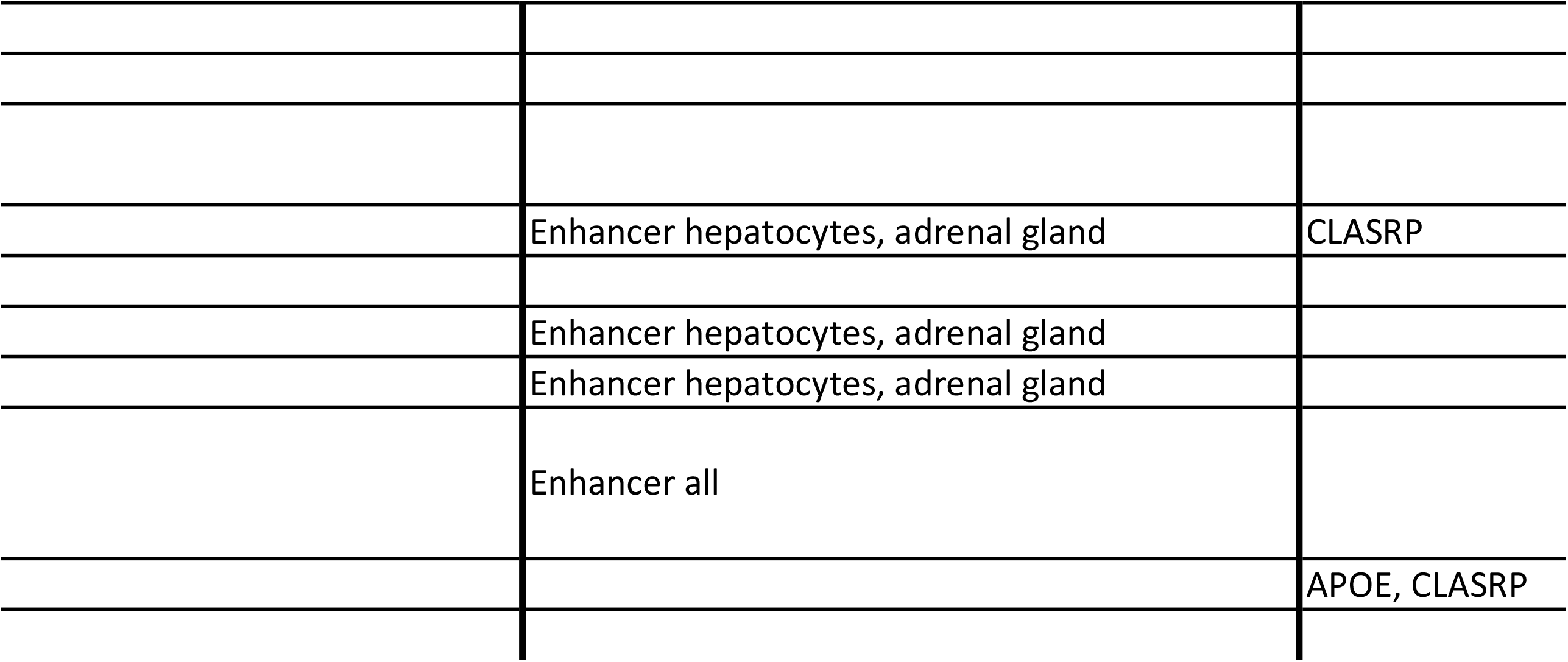

| GTEx | MotifbreakR |
| --- | --- |
| non-brain |  |
| BCAM, PVRL2 |  |
| BCAM, PVRL2 |  |
| BCAM, PVRL2 | RARB & RARG<br>NA |
| BCAM, PVRL2 |  |
| BCAM, PVRL2 |  |
| BCAM, PVRL2 |  |
| BCAM |  |
| TOMM40 |  |
| TOMM40 |  |
| BCAM | TFAIP2A |
| BCAM | FOXD1 |
| BCAM, TOMM40 |  |
| BCAM, APOC1 |  |
| BCAM, TOMM40 |  |
| BCAM | SP1 |
| BCAM | HMGA2 & EHF |
| BCAM |  |
| APOC1 | PLAG |
| APOC1P1, APOE, APOC1 |  |
|  | MAFK |
| APOC1P1, APOE |  |
| APOC1P1, APOE | FEV & EOMES |
| CBLC |  |
| CBLC |  |
| CBLC |  |
| CBLC |  |
| CBLC |  |
| CBLC |  |
| CBLC | ZFX |
| CBLC |  |
| CBLC |  |
| PVRL2 |  |
| APOC1 |  |
| APOC1, APOE | many |
| CBLC |  |
| APOE | PPARG |
| APOC1 |  |
| APOC1 |  |
| APOE |  |
| APOC1, APOE |  |
| APOC1, APOE |  |
| APOE |  |
| APOC1, APOE<br>APOC1, APOE |  |
| CBLC |  |
| CBLC |  |
| CBLC |  |
| CBLC |  |

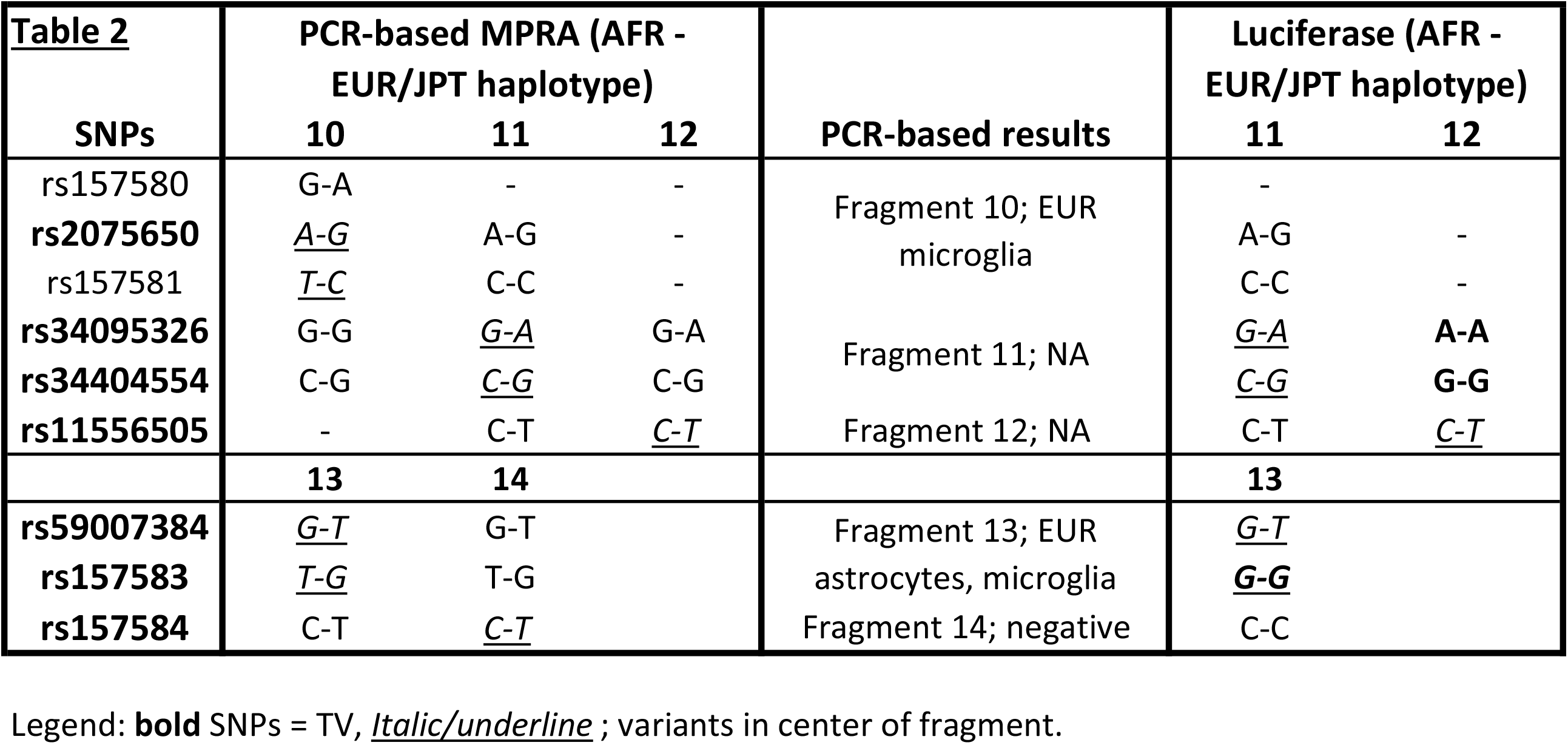

Legend: **bold** SNPs = TV, Italic/underline ; variants in center of fragment.
| Luciferase results | Probe-based MPRA results | ChIP-seq (HaploReg) | score | RegulomeDB JASPAR | MotifBreakR |
| --- | --- | --- | --- | --- | --- |
| Fragment 10; NA | NA | - | 1f | - | - |
|  | negative | - | 1b | RREB1 | SP1 |
|  | NA | - | 2b | ZEB1 | - |
| Fragment 11; AFR neurons | negative | POL2 (K562) | 4 | - | HMGA2 & EHF |
|  | AFR astrocytes | POL2 (K562) | 4 | - | - |
| Fragment 12; negative | negative | POL2 (K562) | 4 | - | - |
| Fragment 13; EUR microglia | EUR microglia | GR (A549) | 4 | - | PLAG |
|  | negative | GR (A549) | 4 | - | - |
| Fragment 14; NA | negative | GR (A549) | 4 | - | - |

| GTE<sub>x</sub> |
| --- |
| APOE skin, APOC1P1 cultured fibroblasts and skin, APOC1 skin<br>BCAM tibial nerve<br>APOC1 mucosa esophagus<br>BCAM tibial nerve<br>BCAM tibial nerve<br>- |
| APOC1 mucosa esophagus and adrenal gland<br>- |
| APOE skin and atrial appendage heart, APOC1 atrial appendage heart |

