## Supplementary material for "Converging evidence for differential regulatory control of *APOEε4* on African versus European haplotypes": Suppl text-figures-tables

**Results.**

2.1. Outcome PCR-based MPRA analyses.

Design.

We designed 39 fragments of ~900bp encompassing a total of 56 variants with differential frequency in European or Japanese versus African haplotypes; with 11 fragments harboring either two or more of the identified sTV variants. These fragments were amplified in heterozygous individuals generating 39 pairs of fragments for inclusion in the library. Of the 39 projected amplicons in the PCR-based MPRA, three were excluded as we were unable to optimize amplification of a 900bp fragment in that region centering around the variant(s) of interest due to repetitive elements.

Sequencing.

Over 143 million paired-end reads were generated for both the HindIII or BglII induced libraries of fragment to barcode ligated vectors; which totaled over 147,000 unique barcodes in either sample. Only barcodes unique across both HindIII and BglII samples and unique for European or African haplotype were used in the further analyses; these amounted to ~51,000 unique barcodes. We generated on average 19 million, 56 million and 28 million single end reads for both replicates of RNA expressed barcodes in microglia, astrocytes and neurons respectively. Of the 51,000 unique barcodes originally identified in the library, we detected ~17,000, ~15,800 and ~14,500 in RNA of microglia, astrocytes and neurons to be used in the differential expression analyses.

Outcomes.

Fragments 2 and 3 showed consistent low incorporation in the library (evidenced by low read numbers in the library of fragment to barcode ligated vectors), indicating Gibson assembly for these two fragments did not perform optimally. Of the remaining 34 fragments, we were able to obtain data on unique barcodes for 24 fragments, though seven of these (15, 16, 23, 31, 32, 36, 38) had low total read counts for the barcodes in the DNA library or expressed RNA samples in the final analyses. For those with sufficient data (**Figure 2B / Table 1 overview / Supplemental Table 1**), we did not identify a significant allele difference for 13 fragments (1, 4, 6, 14, 18, 20, 21, 22, 24, 27, 33, 35 and 39), but observed (nominally) significant differences in both replicates for fragments 13, 17 in astrocytes, fragments 10, 13, 28 in microglia. For both fragment 10 and 13, the haplotype with alleles more common on the European ε4 background induce higher expression, whereas for fragments 17 and 28 the alleles more common on the African ε4 background display higher expression.

### 2.2. Luciferase reporter gene analyses

Of 19 fragments tested in all three cell types (**Figure 2C** / **Table 1 overview / Supplemental Table 1**), we observed very high enhancer activity values compared to the base vector with minimal promotor for fragments 2, 28 (all cell types), 3, 5, 11, 15, 25, 39 (microglia), 7 (neurons), 23 (astrocytes) and 37 (astrocytes, neurons). We detected significant differences ($p<0.05$) in enhancer activity driven by opposite haplotypes for fragments 16 in astrocytes, fragments 8, 13 in microglia and fragments 2, 3, 37 in neurons with >25% activity level difference. Lower fold difference at $p<0.05$ were detected for fragment 7, 13 in astrocytes, 26, 33 in microglia and 11 in neurons. Higher activity of alleles more common on European *APOEε4* was observed for fragments 2, 3, 7, 13, 26 and 33, whereas more common alleles on African *APOEε4* drove higher activity for fragments 2, 8, 11, 16 and 37.

Based on ENCODE ChIP-Seq data indicating many transcription factor binding sites –besides POL2- in fragment 9 overlaying *TOMM40*'s promoter, we evaluated the effect of haplotypes in this fragment in context of promoter activity using pGL4.10 as base vector. We observed a significant higher expression in microglia compared to neuronal or astrocyte cell lines overall as expected and a 2-fold higher expression level of the haplotype more common on European *APOEε4* (driven by rs71352238 C) than the African *APOEε4* background in all three cell types.

### 2.3. Outcome probe-based MPRA analyses

Close to 6 million paired-end reads were generated of the library without reporter gene, in which 3.5 million unique barcodes were detected. We generated on average 118 million, 74 million, 94 million and 164 million single end reads for all three replicates of DNA and RNA expressed barcodes in neurons, microglia and astrocytes respectively. We used 2081 barcodes that were uniquely linked to one allele of one variant in the *APOE* region and were detected in all cell experiments in the differential expression analyses.

Of the 56 variants in the region of interest, we included all non-indel variants (N=51) in the probe-based MPRA design. Of those, three failed the analyses due to insufficient coverage in DNA or expression libraries. We observed significant differences for variants rs34278513 (included in larger fragment 1), rs3865427 (fragment 2), rs71352237 (fragment 3), rs283808 (fragment 5), rs12972970 (fragment 6), rs34342646 (fragment 7), rs71352238 (fragment 9), rs34404554 (fragment 11), rs449647 (fragment 23), rs75627662 (fragment 25) in astrocytes, and rs34342646 (fragment 7) and rs59007384 (fragment 13) in microglia (**Figure 2D**). Alleles more common on either *APOEε4* haplotype show higher activity; European/Japanese rs34278513 (fragment 1), rs3865427 (fragment 2), rs71352237 (fragment 3), rs59007384 (fragment 13), rs75627662 (fragment 25) and rs12721046 (fragment 31) versus African rs283808 (fragment 5), rs12972970 (fragment 6), rs34342646 (fragment 7), rs71352238 (fragment 9), rs34404554 (fragment 11) and rs449647 (fragment 23).

**Methods.**

PCR-based MPRA library creation.

Protocol was adjusted from Trizzino et al.[26]

a. Genomic fragment creation. As per Trizzino et al. [26], we set out to design ~900bp-1kb fragments for inclusion in the library. We used data available through the RoadMap Epigenome project of Frontal Cortex and hippocampus to extract information on functional potential surrounding the 56 SNPs. Primers were designed to amplify the region fully encompassing potential regulatory elements within the ~900bp-1kb fragment regardless of location of SNP therein. Most fragments included two or more of the significant SNPs. Of the final 39 designed fragments, 36 were amplified in a heterozygous carrier previously identified, to ensure equal representation of both alleles. All primers were designed to include 16-18bp overhang with the MCS of pGL4.23 for Gibson assembly (**Supplemental Table 3)**. Concentrations of amplicons was measured through

NanoDrop and qPCR to create an equimolar pool of the fragments (primers available upon request).

b. Barcode fragment creation. A 200bp probe pool was designed to include a XboI cut site, a BglII cut site, a HindIII cut site, a 20bp random barcode and sequence complimentary to Illumina's sequencing primers and a EagI cut site in that order (IDT) (**Supplemental Table 3)**. The probe include sequence complementary to the 3'UTR region of pGL4.23 to allow for Gibson assembly. The probe pool was made doublestranded by 1 PCR cycle using a reverse primer complementary to the end of the fragment in 6 separate reactions (0.5ug probe, 1ul 10mM primer, 5ul 2mM dNTPs, 10ul 5x OneTaq standard buffer and 0.25ul OneTaq (New England Biolabs, NEB) in a 50ul reaction). After reaction clean-up using ethanol purification, ssDNA was removed through ExoI (NEB) digest for 1h at 37C and purified using MinElute in 2x 10ul H20.

c. Library creation. We performed in-vitro mutagenesis to remove a EagI cut site of pGL4.24 right upstream of the Amp gene. This plasmid (~6ug) was digested with EagI-HF and XbaI (NEB) and CIP treated at 37C overnight. After gel extraction and additional CIP treatment, vector was purified using ethanol purification and reconstituted in 40ul H20. We performed 12 parallel Gibson assembly reactions using HIFI builder mastermix, 100ng digested vector and 23ng dsTag in a 20ul reaction for 40min at 50C. Reactions were purified using MinElute in 40ul TE total. We performed transformation of 20ul MegaX-DH10B cells with 2ul of ligation product through electroporation, in 16 parallel reactions; recovered in 1mL of medium for 1h before being combined 2x 8tubes in 100mL cultures each for overnight growth. Vector (pGL4.24+Tag) pools were recovered using a Qiagen maxi prep and eluted in 1mL of TE. For incorporation of the APOE fragment pool, we digested a total of 9ug of pGL4.24+Tag pool with NheI-HF and XhoI and CIP treated overnight at 37C in 3 separate 20ul reactions. After gel extraction and elution in H20, the digested vector pool was ligated with the APOE fragment pool using Gibson assembly in a 1:5 ratio (HIFI builder mastermix 10ul, 100ng digested pGL4.24+Tag, 120ng APOE pool in a 10ul reaction) for 1h at 50C. After Qiagen clean-up, 5 transformations in MegaX-DH10B cells

(2ul ligation product in 20ul cells) were combined in an overnight 100mL culture. The final MPRA vector pool was recovered using Qiagen Maxi prep and elution in 1mL of TE.

d. Fragment to barcode link. To establish the relationship from fragment to barcode we digested out the reporter gene to be able to sequence across barcode to fragment. To prevent the restriction enzyme potentially cutting the APOE fragments, we performed to two separate overnight digestions of 6ug of MPRA pool with either HindIII or BglII. On gel, we extracted bands corresponding to reporter gene-less vector with full length fragments and reporter gene-less vector with shorter (digested) fragments. Extracted fragments were ligated together (100ng vector, 1ul T4 ligase in 10ul reaction) overnight at 4C and purified using Qiagen MinElute. PCR amplicons across APOE fragment and barcode are created using xx primers and standard OneTaq 25ul reaction (5ul 5x OneTaq standard buffer, 2.5mM 2mM dNTPs, 0.25ul OneTaq, 1ul 10mM primer each and 2ul ligation product). PCR products are extracted from gel and included in the tagmentation and sequencing prep. Optimization of the tagmentation protocol (Nextera XT DNA library prep) indicated best results for 50ng of DNA to be tagmentated with 5ul of enzyme for 5min. 10ul of the tagmented product was prepped for sequencing using 2ul of 10uM Illumina index primers, 15ul of Nextera Mastermix, 3ul of 50mM MgCl2, 16ul Betaine and 2.5ul of DMSO in a 50ul reaction.

Probe-based MPRA library creation

We utilized the protocol previously described by Tewhey et al.[27], with the following adjustments. 1146 fragments covering each allele of all 573 single nucleotide variants in the 2MB region surrounding *APOE* were sent to Twist Biosciences for chip manufacturing and guaranteed equal representation. Fragments holding 20bp barcodes and additional adapter sequences were added by regular PCR using the same reagents as reported in Tewhey et al. mpraΔorf library was SPRI purified and eluted in TE to allow for electroporation of 10-beta e.coli cells (NEB, C3020K). Transformed cells were allowed to recover in medium with 100ug/mL Ampicilin. 4.5ug of the mpraΔorf library was digested with AsiSI in a total volume of 200ul. 2ug of the digested library was ligated with ~6ug of GFP amplicon in 250ul volume, after which the product was digested to remove remaining uncut vectors at concentrations corresponding to available amounts of DNA (~1/5 of reported in Tewhey et al). Transformations reached an estimated transformation efficiency of >$10^7$ cfu.

Cell culture

a. Cell types. To evaluate the regulatory potential of haplotypes and single variants in cell types relevant to AD, we performed MPRA and reporter assay analyses in immortalized neuronal (SH-SY5Y), microglia (HMC3), and astrocyte (U-118) cell lines. All were obtained from ATCC (CRL-2266, CRL-3304, HTB-15). For high resolution promoter-focused Capture C experiments, we used HMC3 cells, primary human astrocytes (NHA, Lonza cat#CC-2565), and healthy control iPSC-derived neuroprogenitor cells (NPCs) and neuronal cells. All commercial lines were cultured according to manufacturer's instructions. Cell culture details of the Capture C NPC and neuronal lines are described in Su et al[30].
b. MPRA transfections. For the transfections of cells with the PCR-based vector library, we used 50ug of plasmid library; in a 3:1 ratio of TransIT-LT (MIR2304, MirusBio) to DNA for the microglia and astrocytes, and 1:3 ratio of Lipofectamine 2000 to DNA for the neuronal cell line. We used cells at 90% confluency in a 75$cm^2$ flask in duplicate. For the probe-based fragment vector library we used 65 ug of material with the same reagent ratios and amounts of cells, in triplicate.
c. Luciferase reporter gene assays. The three cell types were seeded into 24-well culture plates at a cell density of $3 \times 10^4$ cells/well. The following day, the cells were co-transfected with the TV fragment bearing firefly luciferase reporter pGL4.24 plasmid (containing either African or European/Japanese haplotype) and the Renilla luciferase regulated by the SV40 promoter (pRL, E2231, Promega) using TransIT®-LT1 (MirusBio) in triplicate at 1: 1/250 :4 ratio. The pRL plasmid served as an internal, non-inducible reporter standard to monitor transfection efficiency.

All firefly luciferase values were normalized to the Renilla control levels. Haplotype pairs were included on the same cell culture plate with same reagent conditions. Twenty-four-hour post-transfection, luciferase activities were measured using the Dual-Glo® Luciferase Assay (E2940, Promega). Results were expressed as the ratio of firefly luciferase activity to renilla luciferase activity. For testing differential promoter activity, haplotypes were cloned into pGL4.10 (E6651, Promega); the same experimental conditions were used as for the enhancer activity assays.

RNA preparations MPRA

a. PCR-based design. RNA was extracted from cell lines, using Qiagen RNeasy kits, with on-column DNAse treatment. cDNA encompassing the barcode/tag from the library expressed RNAs (2ug) was derived using the Qiagen OneStep RT-PCR kit (212012, Qiagen) and primers including Illumina indices and adaptors (**Supplementary Table 3**) according to the manufacturer's protocol. cDNA was purified using the PCR reaction clean-up kit (28004, Qiagen).
b. Probe-based design. RNA was extracted as described in Tewhey et al[29]. In brief, GFP RNA is isolated using biotin labelled probes, converted to cDNA and the sequencing library prepared using primers that include Illumina indices and adaptors (**Supplementary Table 3**).

Sequencing

We performed 2x100bp paired-end sequencing on the Illumina HiSeq3000- for the determination of barcode to fragment link in the tagmented library of the PCR-based MPRA. Sequencing was performed at the Center for Genome Technology (CGT) at the John P. Hussman Institute for Human Genomics (HIHG). Using the subassembly method described in Trizzino et al[28], paired reads were grouped per barcode in read 1 after which sequences of read 2 were used to reassemble the larger fragment and determine variant allele presence. For the probe-based MPRA we performed 2x150bp sequencing on the Illumina NovaSeq at the CGT allowing for complete read-through of the total assembled amplicon harboring the fragment with the variant allele of interest and barcode. Barcodes matching multiple fragments or opposite alleles in the same fragment within either set-up were excluded from further analyses.

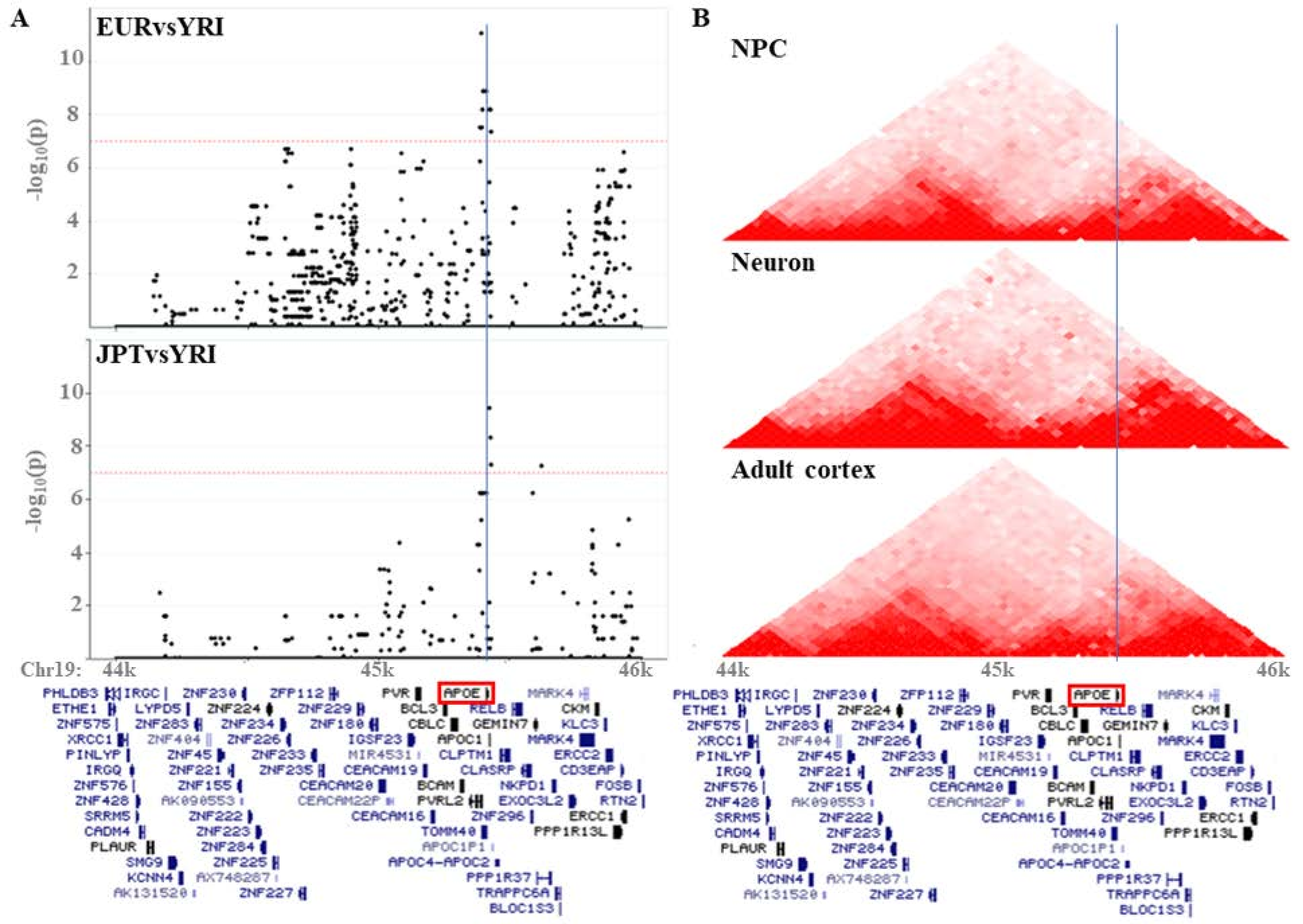
A
EURvsYRI
JPTvsYRI
-log10(p)
10
8
6
4
2
Chr19:
44k
45k
46k
B
NPC
Neuron
Adult cortex
PHLDB3
IRGC
ZNF230
ZFP112
PVR
APOE
MARK4
ETHE1
LYPD5
ZNF224
ZNF229
BCL3
RELB
CKM
ZNF575
ZNF283
ZNF234
ZNF180
CBLC
GEMIN7
KLC3
XRCC1
ZNF404
ZNF226
IGSF23
APOC1
MARK4
PINLYP
ZNF45
ZNF233
MIR4531
CLPTM1
ERCC2
IRGQ
ZNF221
ZNF235
CEACAM19
CLASRP
CD3EAP
ZNF576
ZNF155
CEACAM20
BCAM
NKPD1
FOSB
ZNF428
AK090553
CEACAM22P
PVRL2
EXOC3L2
RTN2
SRRM5
ZNF222
CEACAM16
ZNF296
ERCC1
CADM4
ZNF223
TOMM40
PPP1R13L
PLAUR
ZNF284
APOC1P1
SMG9
ZNF225
APOC4-APOC2
KCNN4
AX748287
PPP1R37
AK131520
ZNF227
TRAPPC6A
BLOC1S3

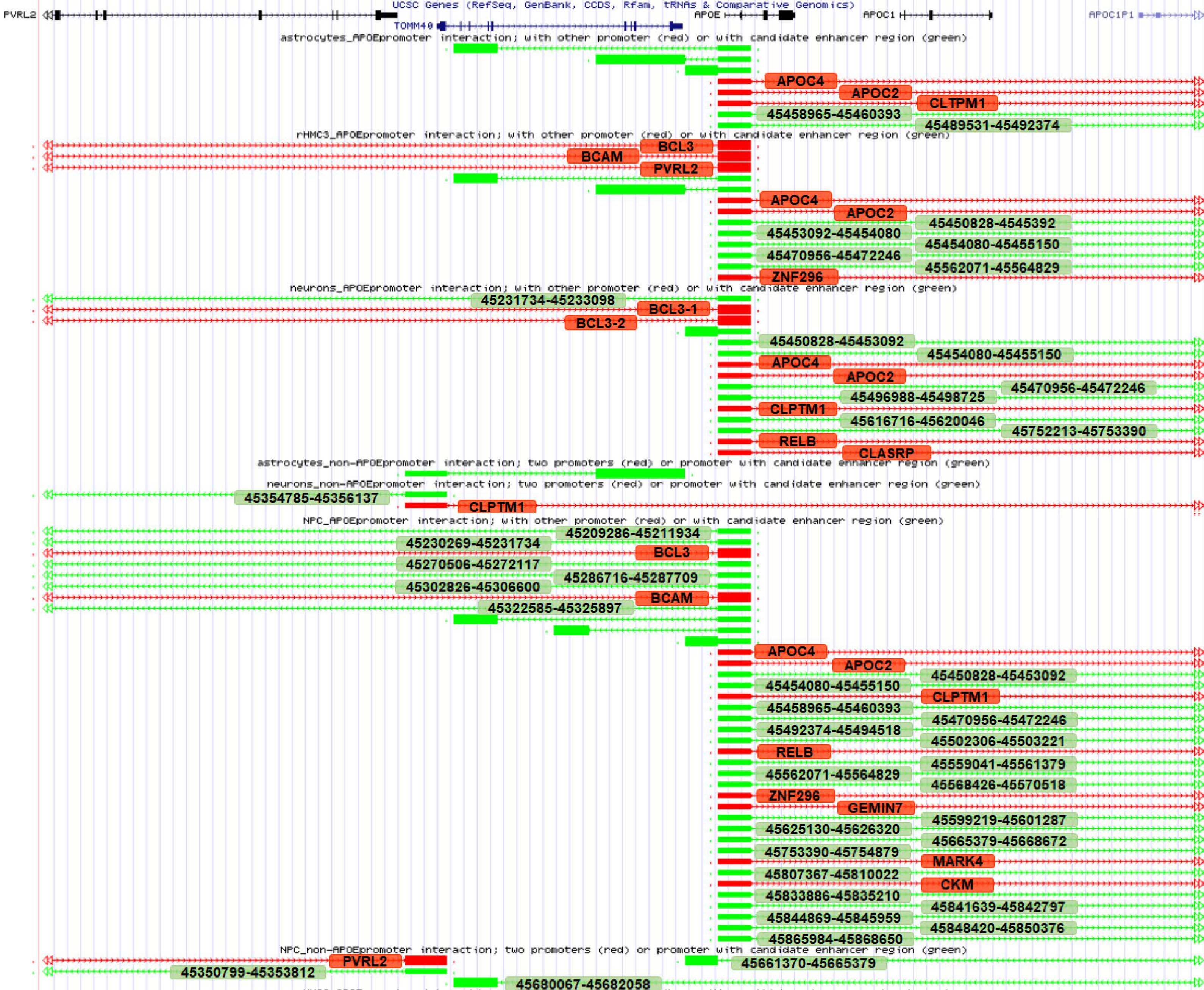
UCSC Genes (RefSeq, GenBank, CCDS, Rfam, tRNAs & Comparative Genomics)
PVRL2
TOMM40
APOE
APOC1
APOC1P1
astrocytes_APOEpromoter interaction; with other promoter (red) or with candidate enhancer region (green)
APOC4
APOC2
CLTPM1
45458965-45460393
45489531-45492374
rHMC3_APOEpromoter interaction; with other promoter (red) or with candidate enhancer region (green)
BCL3
BCAM
PVRL2
APOC4
APOC2
45450828-4545392
45453092-45454080
45454080-45455150
45470956-45472246
45562071-45564829
ZNF296
neurons_APOEpromoter interaction; with other promoter (red) or with candidate enhancer region (green)
45231734-45233098
BCL3-1
BCL3-2
45450828-45453092
45454080-45455150
APOC4
APOC2
45470956-45472246
45496988-45498725
CLPTM1
45616716-45620046
45752213-45753390
RELB
CLASRP
astrocytes_non-APOEpromoter interaction; two promoters (red) or promoter with candidate enhancer region (green)
neurons_non-APOEpromoter interaction; two promoters (red) or promoter with candidate enhancer region (green)
45354785-45356137
CLPTM1
NPC_APOEpromoter interaction; with other promoter (red) or with candidate enhancer region (green)
45209286-45211934
45230269-45231734
BCL3
45270506-45272117
45286716-45287709
45302826-45306600
BCAM
45322585-45325897
APOC4
APOC2
45450828-45453092
45454080-45455150
CLPTM1
45458965-45460393
45470956-45472246
45492374-45494518
45502306-45503221
RELB
45559041-45561379
45562071-45564829
45568426-45570518
ZNF296
GEMIN7
45599219-45601287
45625130-45626320
45665379-45668672
45753390-45754879
MARK4
45807367-45810022
CKM
45833886-45835210
45841639-45842797
45844869-45845959
45848420-45850376
45865984-45868650
NPC_non-APOEpromoter interaction; two promoters (red) or promoter with candidate enhancer region (green)
PVRL2
45661370-45665379
45350799-45353812
45680067-45682058

**Suppl Table 1: PCR-based MPRA results**

| Fragment | A_APOEe4 | R_APOEe4 | DNA_A | DNA_R | A/(A+R) | Experiment | RNA_A | RNA_R |
|---|---|---|---|---|---|---|---|---|
| **Fragments with significant replicates** | | | | | | | | |
| 10 | EUR | AFR | 104 | 618 | 0.144044 | microglia-rep1 | 158 | 751 |
| 10 | | | 104 | 618 | | microglia-rep2 | 157 | 616 |
| 13 | EUR | AFR | 652 | 2629 | 0.19872 | microglia-rep1 | 919 | 3270 |
| 13 | | | 652 | 2629 | | microglia-rep2 | 826 | 2624 |
| 13 | EUR | AFR | 652 | 2629 | 0.19872 | astrocytes-rep1 | 1953 | 7005 |
| 13 | | | 652 | 2629 | | astrocytes-rep2 | 1630 | 5423 |
| 17 | AFR | EUR | 325 | 273 | 0.543478 | astrocytes-rep1 | 1050 | 693 |
| 17 | | | 325 | 273 | | astrocytes-rep2 | 762 | 540 |
| 28 | EUR | AFR | 186 | 450 | 0.292453 | microglia-rep1 | 165 | 596 |
| 28 | | | 186 | 450 | | microglia-rep2 | 142 | 463 |
| **Fragments with negative results** | | | | | | | | |
| 1 | EUR | AFR | 418 | 774 | 0.350671 | microglia-rep1 | 501 | 888 |
| 1 | | | 418 | 774 | | microglia-rep2 | 432 | 682 |
| 1 | | | 418 | 774 | | neurons-rep1 | 569 | 980 |
| 1 | | | 418 | 774 | | neurons-rep2 | 706 | 1111 |
| 1 | | | 418 | 774 | | astrocytes-rep1 | 1038 | 1834 |
| 1 | | | 418 | 774 | | astrocytes-rep2 | 856 | 1389 |
| 4 | EUR | AFR | 212 | 28698 | 0.007333 | microglia-rep1 | 282 | 36203 |
| 4 | | | 212 | 28698 | | microglia-rep2 | 231 | 29011 |
| 4 | | | 212 | 28698 | | neurons-rep1 | 310 | 40367 |
| 4 | | | 212 | 28698 | | neurons-rep2 | 462 | 49215 |
| 4 | | | 212 | 28698 | | astrocytes-rep1 | 754 | 85073 |
| 4 | | | 212 | 28698 | | astrocytes-rep2 | 525 | 69186 |
| 6 | EUR | AFR | 1032 | 619 | 0.625076 | microglia-rep1 | 1197 | 775 |
| 6 | | | 1032 | 619 | | microglia-rep2 | 997 | 663 |
| 6 | | | 1032 | 619 | | neurons-rep1 | 1465 | 880 |
| 6 | | | 1032 | 619 | | neurons-rep2 | 1628 | 1039 |
| 6 | | | 1032 | 619 | | astrocytes-rep1 | 2855 | 1750 |
| 6 | | | 1032 | 619 | | astrocytes-rep2 | 2373 | 1385 |
| 10 | EUR | AFR | 104 | 618 | 0.144044 | neurons-rep1 | 117 | 860 |
| 10 | | | 104 | 618 | | neurons-rep2 | 213 | 1001 |
| 10 | | | 104 | 618 | | astrocytes-rep1 | 364 | 1918 |
| 10 | | | 104 | 618 | | astrocytes-rep2 | 324 | 1487 |
| 13 | EUR | AFR | 652 | 2629 | 0.19872 | neurons-rep1 | 967 | 3465 |
| 13 | | | 652 | 2629 | | neurons-rep2 | 1074 | 4005 |
| 14 | AFR | EUR | 695 | 284 | 0.709908 | microglia-rep1 | 802 | 341 |
| 14 | | | 695 | 284 | | microglia-rep2 | 710 | 307 |
| 14 | | | 695 | 284 | | neurons-rep1 | 970 | 326 |
| 14 | | | 695 | 284 | | neurons-rep2 | 1132 | 471 |
| 14 | | | 695 | 284 | | astrocytes-rep1 | 1883 | 755 |
| 14 | | | 695 | 284 | | astrocytes-rep2 | 1385 | 512 |
| 17 | AFR | EUR | 325 | 273 | 0.543478 | microglia-rep1 | 402 | 307 |
| 17 | | | 325 | 273 | | microglia-rep2 | 325 | 235 |

| | | | | | | | | |
|---|---|---|---|---|---|---|---|---|
| 17 | | | 325 | 273 | | neurons-rep1 | 480 | 288 |
| 17 | | | 325 | 273 | | neurons-rep2 | 573 | 428 |
| 18 | AFR | EUR | 793 | 1336 | 0.372475 | microglia-rep1 | 966 | 1732 |
| 18 | | | 793 | 1336 | | microglia-rep2 | 801 | 1280 |
| 18 | | | 793 | 1336 | | neurons-rep1 | 1130 | 1833 |
| 18 | | | 793 | 1336 | | neurons-rep2 | 1325 | 2173 |
| 18 | | | 793 | 1336 | | astrocytes-rep1 | 2118 | 3807 |
| 18 | | | 793 | 1336 | | astrocytes-rep2 | 1973 | 2832 |
| 20 | AFR | EUR | 324 | 369 | 0.467532 | microglia-rep1 | 374 | 456 |
| 20 | | | 324 | 369 | | microglia-rep2 | 268 | 340 |
| 20 | | | 324 | 369 | | neurons-rep1 | 375 | 520 |
| 20 | | | 324 | 369 | | neurons-rep2 | 516 | 567 |
| 20 | | | 324 | 369 | | astrocytes-rep1 | 835 | 936 |
| 20 | | | 324 | 369 | | astrocytes-rep2 | 681 | 722 |
| 21 | AFR | EUR | 816 | 192 | 0.809524 | microglia-rep1 | 1076 | 263 |
| 21 | | | 816 | 192 | | microglia-rep2 | 881 | 213 |
| 21 | | | 816 | 192 | | neurons-rep1 | 1326 | 260 |
| 21 | | | 816 | 192 | | neurons-rep2 | 1489 | 342 |
| 21 | | | 816 | 192 | | astrocytes-rep1 | 2333 | 533 |
| 21 | | | 816 | 192 | | astrocytes-rep2 | 2001 | 438 |
| 22 | AFR | EUR | 530 | 204 | 0.722071 | microglia-rep1 | 692 | 219 |
| 22 | | | 530 | 204 | | microglia-rep2 | 542 | 253 |
| 22 | | | 530 | 204 | | neurons-rep1 | 702 | 237 |
| 22 | | | 530 | 204 | | neurons-rep2 | 800 | 318 |
| 22 | | | 530 | 204 | | astrocytes-rep1 | 1289 | 540 |
| 22 | | | 530 | 204 | | astrocytes-rep2 | 1164 | 380 |
| 24 | EUR | AFR | 1242 | 1329 | 0.483081 | microglia-rep1 | 1630 | 1727 |
| 24 | | | 1242 | 1329 | | microglia-rep2 | 1296 | 1434 |
| 24 | | | 1242 | 1329 | | neurons-rep1 | 1723 | 1868 |
| 24 | | | 1242 | 1329 | | neurons-rep2 | 2039 | 2185 |
| 24 | | | 1242 | 1329 | | astrocytes-rep1 | 3670 | 3935 |
| 24 | | | 1242 | 1329 | | astrocytes-rep2 | 3070 | 3126 |
| 27 | AFR | EUR | 654 | 842 | 0.437166 | microglia-rep1 | 812 | 1031 |
| 27 | | | 654 | 842 | | microglia-rep2 | 651 | 901 |
| 27 | | | 654 | 842 | | neurons-rep1 | 990 | 1231 |
| 27 | | | 654 | 842 | | neurons-rep2 | 965 | 1496 |
| 27 | | | 654 | 842 | | astrocytes-rep1 | 1855 | 2475 |
| 27 | | | 654 | 842 | | astrocytes-rep2 | 1422 | 2132 |
| 28 | EUR | AFR | 186 | 450 | 0.292453 | neurons-rep1 | 147 | 753 |
| 28 | | | 186 | 450 | | neurons-rep2 | 259 | 721 |
| 28 | | | 186 | 450 | | astrocytes-rep1 | 494 | 1279 |
| 28 | | | 186 | 450 | | astrocytes-rep2 | 319 | 1177 |
| 32 | EUR | AFR | 72 | 179 | 0.286853 | microglia-rep1 | 90 | 183 |
| 32 | | | 72 | 179 | | microglia-rep2 | 65 | 139 |
| 32 | | | 72 | 179 | | neurons-rep1 | 122 | 246 |
| 32 | | | 72 | 179 | | neurons-rep2 | 105 | 262 |
| 32 | | | 72 | 179 | | astrocytes-rep1 | 239 | 540 |

| | | | | | | | | |
|---|---|---|---|---|---|---|---|---|
| 32 | | | 72 | 179 | | astrocytes-rep2 | 217 | 350 |
| 33 | EUR | AFR | 899 | 1673 | 0.349533 | microglia-rep1 | 1268 | 2024 |
| 33 | | | 899 | 1673 | | microglia-rep2 | 782 | 1732 |
| 33 | | | 899 | 1673 | | neurons-rep1 | 1290 | 2319 |
| 33 | | | 899 | 1673 | | neurons-rep2 | 1391 | 2879 |
| 33 | | | 899 | 1673 | | astrocytes-rep1 | 2209 | 4770 |
| 33 | | | 899 | 1673 | | astrocytes-rep2 | 1997 | 3711 |
| 35 | AFR | EUR | 1832 | 2877 | 0.389042 | microglia-rep1 | 2453 | 3591 |
| 35 | | | 1832 | 2877 | | microglia-rep2 | 2103 | 3145 |
| 35 | | | 1832 | 2877 | | neurons-rep1 | 2779 | 4039 |
| 35 | | | 1832 | 2877 | | neurons-rep2 | 3254 | 4832 |
| 35 | | | 1832 | 2877 | | astrocytes-rep1 | 5638 | 7607 |
| 35 | | | 1832 | 2877 | | astrocytes-rep2 | 4216 | 6479 |
| 39 | AFR | EUR | 2758 | 4090 | 0.402745 | microglia-rep1 | 3436 | 4721 |
| 39 | | | 2758 | 4090 | | microglia-rep2 | 2506 | 4004 |
| 39 | | | 2758 | 4090 | | neurons-rep1 | 3755 | 5413 |
| 39 | | | 2758 | 4090 | | neurons-rep2 | 4421 | 6599 |
| 39 | | | 2758 | 4090 | | astrocytes-rep1 | 7757 | 11342 |
| 39 | | | 2758 | 4090 | | astrocytes-rep2 | 6291 | 8978 |
| **Fragments with low coverage** | | | | | | | | |
| 15 | AFR | EUR | 34 | 3 | 0.918919 | microglia-rep1 | 29 | 10 |
| 15 | | | 34 | 3 | | microglia-rep2 | 20 | 8 |
| 15 | | | 34 | 3 | | neurons-rep1 | 33 | 5 |
| 15 | | | 34 | 3 | | neurons-rep2 | 37 | 11 |
| 15 | | | 34 | 3 | | astrocytes-rep1 | 63 | 16 |
| 15 | | | 34 | 3 | | astrocytes-rep2 | 68 | 21 |
| 16 | AFR | EUR | 27 | 56 | 0.325301 | microglia-rep1 | 31 | 67 |
| 16 | | | 27 | 56 | | microglia-rep2 | 19 | 49 |
| 16 | | | 27 | 56 | | neurons-rep1 | 31 | 63 |
| 16 | | | 27 | 56 | | neurons-rep2 | 30 | 82 |
| 16 | | | 27 | 56 | | astrocytes-rep1 | 28 | 142 |
| 16 | | | 27 | 56 | | astrocytes-rep2 | 36 | 105 |
| 23 | AFR | EUR | 8 | 22 | 0.266667 | microglia-rep1 | 10 | 23 |
| 23 | | | 8 | 22 | | microglia-rep2 | 6 | 13 |
| 23 | | | 8 | 22 | | neurons-rep1 | 7 | 26 |
| 23 | | | 8 | 22 | | neurons-rep2 | 6 | 37 |
| 23 | | | 8 | 22 | | astrocytes-rep1 | 27 | 95 |
| 23 | | | 8 | 22 | | astrocytes-rep2 | 25 | 28 |
| 31 | EUR | AFR | 28 | 25 | 0.528302 | microglia-rep1 | 54 | 52 |
| 31 | | | 28 | 25 | | microglia-rep2 | 73 | 74 |
| 31 | | | 28 | 25 | | neurons-rep1 | 36 | 42 |
| 31 | | | 28 | 25 | | neurons-rep2 | 82 | 37 |
| 31 | | | 28 | 25 | | astrocytes-rep1 | 254 | 253 |
| 31 | | | 28 | 25 | | astrocytes-rep2 | 109 | 83 |
| 36 | AFR | EUR | 50 | 54 | 0.480769 | microglia-rep1 | 145 | 107 |
| 36 | | | 50 | 54 | | microglia-rep2 | 128 | 125 |
| 36 | | | 50 | 54 | | neurons-rep1 | 162 | 101 |

| | | | | | | | | |
|---|---|---|---|---|---|---|---|---|
| 36 | | | 50 | 54 | | neurons-rep2 | 147 | 118 |
| 36 | | | 50 | 54 | | astrocytes-rep1 | 2327 | 1609 |
| 36 | | | 50 | 54 | | astrocytes-rep2 | 574 | 399 |
| 38 | AFR | EUR | 6 | 7 | 0.461538 | microglia-rep1 | 1 | 15 |
| 38 | | | 6 | 7 | | microglia-rep2 | 3 | 12 |
| 38 | | | 6 | 7 | | neurons-rep1 | 0 | 15 |
| 38 | | | 6 | 7 | | neurons-rep2 | 8 | 10 |
| 38 | | | 6 | 7 | | astrocytes-rep1 | 29 | 24 |
| 38 | | | 6 | 7 | | astrocytes-rep2 | 23 | 24 |

| A/(A+R) | conf.low | conf.high | P-value |
|---|---|---|---|
| 0.173817 | 0.149721 | 0.200048 | **0.034001** |
| 0.203105 | 0.175274 | 0.233221 | **5.71E-05** |
| 0.219384 | 0.206932 | 0.232229 | **0.004218** |
| 0.23942 | 0.225262 | 0.254022 | **1.04E-07** |
| 0.218017 | 0.209504 | 0.226715 | **5.24E-05** |
| 0.231107 | 0.221311 | 0.241126 | **5.62E-10** |
| 0.60241 | 0.578991 | 0.625483 | **1.02E-05** |
| 0.585253 | 0.557941 | 0.612181 | **0.009024** |
| 0.21682 | 0.188029 | 0.247824 | **3.16E-05** |
| 0.234711 | 0.201477 | 0.270558 | **0.00655** |
| 0.360691 | 0.335395 | 0.386578 | 0.572535 |
| 0.387792 | 0.359059 | 0.417119 | **0.028916** |
| 0.367334 | 0.343274 | 0.391897 | 0.31538 |
| 0.388553 | 0.366061 | 0.411405 | **0.003984** |
| 0.361421 | 0.343825 | 0.379299 | 0.367866 |
| 0.381292 | 0.361144 | 0.40175 | **0.009024** |
| 0.007729 | 0.006856 | 0.008682 | 0.518708 |
| 0.0079 | 0.006917 | 0.008982 | 0.390744 |
| 0.007621 | 0.006799 | 0.008515 | 0.617223 |
| 0.0093 | 0.008475 | 0.010184 | **1.02E-05** |
| 0.008785 | 0.008172 | 0.009432 | **1.47E-05** |
| 0.007531 | 0.006903 | 0.008201 | 0.664349 |
| 0.606998 | 0.585041 | 0.628636 | 0.192235 |
| 0.600602 | 0.576579 | 0.624269 | *0.095353* |
| 0.624733 | 0.604777 | 0.644378 | 0.996268 |
| 0.610424 | 0.591616 | 0.628989 | 0.225559 |
| 0.619978 | 0.605778 | 0.634026 | 0.60825 |
| 0.631453 | 0.615799 | 0.646902 | 0.572535 |
| 0.119754 | 0.100054 | 0.141778 | *0.07895* |
| 0.175453 | 0.154452 | 0.198039 | **0.008791** |
| 0.159509 | 0.144712 | 0.175186 | *0.089273* |
| 0.178907 | 0.16151 | 0.19735 | **0.000272** |
| 0.218186 | 0.206102 | 0.230642 | **0.005417** |
| 0.211459 | 0.200299 | 0.222952 | *0.060995* |
| 0.701662 | 0.674206 | 0.728077 | 0.664349 |
| 0.698132 | 0.668882 | 0.72623 | 0.555165 |
| 0.748457 | 0.723901 | 0.77188 | **0.008028** |
| 0.706176 | 0.683202 | 0.728392 | 0.811603 |
| 0.713798 | 0.69613 | 0.730991 | 0.782755 |
| 0.7301 | 0.709515 | 0.749971 | 0.122011 |
| 0.566996 | 0.529603 | 0.603829 | 0.360009 |
| 0.580357 | 0.538256 | 0.621606 | 0.170544 |

| | | | |
|---|---|---|---|
| 0.625 | 0.589682 | 0.659354 | **5.24E-05** |
| 0.572428 | 0.541105 | 0.603323 | 0.148907 |
| 0.358043 | 0.339929 | 0.376465 | 0.231905 |
| 0.384911 | 0.36394 | 0.406207 | 0.386844 |
| 0.38137 | 0.363837 | 0.399138 | 0.461664 |
| 0.378788 | 0.362677 | 0.395102 | 0.581169 |
| 0.357468 | 0.345254 | 0.369824 | **0.04339** |
| 0.410614 | 0.396658 | 0.424679 | **9.22E-07** |
| 0.450602 | 0.416378 | 0.485179 | 0.467465 |
| 0.440789 | 0.400874 | 0.481283 | 0.333579 |
| 0.418994 | 0.386424 | 0.4521 | **0.011364** |
| 0.476454 | 0.446346 | 0.506691 | 0.681008 |
| 0.471485 | 0.448017 | 0.495047 | 0.811603 |
| 0.485388 | 0.458934 | 0.511904 | 0.319573 |
| 0.803585 | 0.781275 | 0.824553 | 0.693259 |
| 0.805302 | 0.780572 | 0.828376 | 0.811603 |
| 0.836066 | 0.816912 | 0.853966 | **0.019687** |
| 0.813217 | 0.794596 | 0.830829 | 0.811603 |
| 0.814027 | 0.799289 | 0.82812 | 0.678858 |
| 0.820418 | 0.804598 | 0.835465 | 0.319573 |
| 0.759605 | 0.730494 | 0.787026 | **0.033571** |
| 0.681761 | 0.648124 | 0.714044 | **0.034308** |
| 0.747604 | 0.718538 | 0.775108 | 0.172242 |
| 0.715564 | 0.688128 | 0.74186 | 0.756214 |
| 0.704757 | 0.683262 | 0.725593 | 0.192572 |
| 0.753886 | 0.731606 | 0.775196 | **0.016386** |
| 0.485553 | 0.468516 | 0.502615 | 0.832256 |
| 0.474725 | 0.455846 | 0.493659 | 0.535107 |
| 0.479811 | 0.463353 | 0.496301 | 0.800057 |
| 0.482718 | 0.467546 | 0.497913 | 0.982823 |
| 0.482577 | 0.471289 | 0.493879 | 0.968263 |
| 0.495481 | 0.482956 | 0.508011 | 0.119509 |
| 0.440586 | 0.417759 | 0.463602 | 0.832256 |
| 0.419459 | 0.394758 | 0.444465 | 0.305581 |
| 0.445745 | 0.424932 | 0.466702 | 0.562734 |
| 0.392117 | 0.372763 | 0.411728 | **5.31E-05** |
| 0.428406 | 0.413604 | 0.443305 | 0.387534 |
| 0.400113 | 0.383953 | 0.416437 | **5.71E-05** |
| 0.163333 | 0.13976 | 0.189134 | **2.08E-17** |
| 0.264286 | 0.23691 | 0.293085 | 0.121484 |
| 0.278624 | 0.257854 | 0.300129 | 0.358252 |
| 0.213235 | 0.192725 | 0.234878 | **1.63E-10** |
| 0.32967 | 0.274197 | 0.38889 | 0.231905 |
| 0.318627 | 0.255301 | 0.387318 | 0.458647 |
| 0.331522 | 0.283592 | 0.382189 | 0.141209 |
| 0.286104 | 0.240393 | 0.335304 | 1 |
| 0.306804 | 0.27456 | 0.340517 | 0.361985 |

| | | | |
|---|---|---|---|
| 0.382716 | 0.342529 | 0.424132 | **1.23E-05** |
| 0.385176 | 0.368509 | 0.402048 | **0.000133** |
| 0.311058 | 0.292992 | 0.329565 | **0.000295** |
| 0.35744 | 0.341786 | 0.373324 | 0.461253 |
| 0.325761 | 0.31171 | 0.340051 | **0.004946** |
| 0.316521 | 0.305616 | 0.32758 | **1.14E-07** |
| 0.34986 | 0.337479 | 0.362394 | 0.981915 |
| 0.405857 | 0.393442 | 0.418363 | **0.022645** |
| 0.400724 | 0.387429 | 0.41413 | 0.170544 |
| 0.407598 | 0.395902 | 0.419372 | **0.00655** |
| 0.402424 | 0.39171 | 0.413208 | **0.03685** |
| 0.42567 | 0.417229 | 0.434144 | **3.88E-16** |
| 0.394203 | 0.384925 | 0.403539 | 0.413009 |
| 0.421233 | 0.410487 | 0.432036 | **0.003513** |
| 0.384946 | 0.373103 | 0.396892 | **0.011** |
| 0.409577 | 0.399486 | 0.419725 | 0.319581 |
| 0.40118 | 0.39201 | 0.410401 | 0.811603 |
| 0.406147 | 0.39917 | 0.413152 | 0.473238 |
| 0.412011 | 0.404189 | 0.419867 | *0.050808* |
| 0.74359 | 0.578726 | 0.869623 | **0.004068** |
| 0.714286 | 0.513332 | 0.867763 | **0.005417** |
| 0.868421 | 0.719136 | 0.955863 | 0.367866 |
| 0.770833 | 0.62688 | 0.87967 | **0.005417** |
| 0.797468 | 0.692043 | 0.879573 | **0.002868** |
| 0.764045 | 0.662212 | 0.847619 | **5.46E-05** |
| 0.316327 | 0.226059 | 0.41804 | 0.952449 |
| 0.279412 | 0.177343 | 0.401459 | 0.6524 |
| 0.329787 | 0.236248 | 0.434385 | 0.952449 |
| 0.267857 | 0.188579 | 0.359803 | 0.367866 |
| 0.164706 | 0.112324 | 0.229172 | **3.54E-05** |
| 0.255319 | 0.18567 | 0.335539 | 0.172242 |
| 0.30303 | 0.155917 | 0.487111 | 0.800057 |
| 0.315789 | 0.125761 | 0.565502 | 0.725066 |
| 0.212121 | 0.089804 | 0.389081 | 0.681008 |
| 0.139535 | 0.052977 | 0.279325 | 0.130856 |
| 0.221311 | 0.151196 | 0.305421 | 0.449554 |
| 0.471698 | 0.333025 | 0.613645 | **0.006164** |
| 0.509434 | 0.410485 | 0.607843 | 0.800057 |
| 0.496599 | 0.41317 | 0.580167 | 0.592119 |
| 0.461538 | 0.347927 | 0.578158 | 0.390744 |
| 0.689076 | 0.59774 | 0.770737 | **0.002464** |
| 0.500986 | 0.456581 | 0.54538 | 0.367866 |
| 0.567708 | 0.494444 | 0.638849 | 0.414695 |
| 0.575397 | 0.511799 | 0.637199 | **0.009934** |
| 0.505929 | 0.442592 | 0.569125 | 0.587437 |
| 0.61597 | 0.554248 | 0.675046 | **9.52E-05** |

| | | | |
|---|---|---|---|
| 0.554717 | 0.492664 | 0.615532 | **0.04339** |
| 0.591209 | 0.575661 | 0.606622 | **1.44E-41** |
| 0.589928 | 0.558276 | 0.621034 | **3.23E-10** |
| 0.0625 | 0.001581 | 0.302321 | **0.003984** |
| 0.2 | 0.043312 | 0.480891 | 0.142212 |
| 0 | 0 | 0.218019 | **0.000588** |
| 0.444444 | 0.215302 | 0.692428 | 1 |
| 0.54717 | 0.404498 | 0.684385 | 0.361985 |
| 0.489362 | 0.340757 | 0.639355 | 0.832256 |

| Suppl Table 1 | | luciferase/renilla | | | | | | |
|---|---|---|---|---|---|---|---|---|
| Luciferase results | | Microglia | | | Astrocytes | | | |
| **Fragments** | **Experiment** | **AFR** | **EUR** | **t-test** | **AFR** | **EUR** | **t-test** | **AFR** |
| **2** | **rep1** | 6.432 | 4.226 | 0.589 | 2.051 | 2.024 | 0.051 | 3.139 |
| | **rep2** | 6.252 | 6.456 | | 2.031 | 2.183 | | 3.384 |
| | **rep3** | 5.726 | 7.316 | | 1.797 | 2.106 | | 3.052 |
| | **rep4** | 6.300 | 5.103 | | 1.752 | 2.142 | | - |
| **3** | **rep1** | 3.571 | 2.575 | 0.124 | 1.384 | 1.273 | 0.489 | 0.860 |
| | **rep2** | 5.348 | 3.699 | | 1.025 | 1.214 | | 0.794 |
| | **rep3** | 4.342 | 3.404 | | 1.276 | 1.770 | | 0.722 |
| | **rep4** | - | - | | 1.517 | 1.424 | 0.550 | 0.810 |
| **5** | **rep1** | 3.297 | 3.411 | 0.804 | 0.832 | 1.018 | 0.763 | 1.188 |
| | **rep2** | 3.004 | 2.894 | | 1.410 | 0.996 | | 1.010 |
| | **rep3** | - | 2.954 | | 1.184 | 1.339 | | 1.162 |
| | **rep4** | - | - | | 1.206 | 1.096 | | 1.209 |
| **7** | **rep1** | 1.158 | 1.134 | 0.406 | 0.993 | 1.068 | **0.016** | 2.536 |
| | **rep2** | 1.202 | 1.121 | | 1.008 | 1.295 | | 2.725 |
| | **rep3** | 1.042 | 0.888 | | 0.991 | 1.119 | 0.885 | 2.365 |
| | **rep4** | - | - | | 0.961 | 1.322 | | - |
| **8** | **rep1** | 0.644 | 0.351 | **0.020** | 0.881 | 1.064 | 0.868 | 1.387 |
| | **rep2** | 0.769 | 0.603 | | 1.586 | 1.177 | | 1.567 |
| | **rep3** | 0.853 | 0.402 | | 1.286 | 1.506 | | 1.414 |
| | **rep4** | 0.560 | 0.361 | | 1.388 | - | | 1.500 |
| **9** | **rep1** | 754.276 | 2008.088 | **0.00004** | 131.724 | 287.862 | **0.000013** | 109.010 |
| **promoter** | **rep2** | 942.963 | 1898.087 | | 118.513 | 307.294 | | 108.805 |
| | **rep3** | 832.015 | 1617.989 | | 132.702 | 279.972 | | 116.746 |
| | **rep4** | 826.890 | 2004.584 | | 106.334 | - | | 99.501 |
| **11** | **rep1** | 3.095 | 3.200 | 0.904 | 1.491 | 1.433 | 0.572 | 2.276 |
| | **rep2** | 3.451 | 4.154 | | 1.458 | 1.417 | | 2.321 |
| | **rep3** | 2.753 | 3.042 | | 1.686 | 1.572 | | 2.419 |
| | **rep4** | 4.023 | 2.719 | | - | 1.570 | | - |
| **12** | **rep1** | 1.235 | 1.002 | 0.217 | 0.783 | 0.576 | 0.106 | 0.465 |
| | **rep2** | 1.097 | 1.210 | | 0.719 | 0.673 | | 0.676 |
| | **rep3** | 1.174 | 1.138 | | 0.836 | 0.755 | | 0.593 |
| | **rep4** | 1.233 | 1.089 | | 0.669 | 0.552 | | - |
| **13** | **rep1** | 0.349 | 0.785 | **0.001** | 0.686 | 0.856 | **0.0003** | 0.832 |
| | **rep2** | 0.300 | 0.745 | | 0.647 | 0.863 | | 0.781 |
| | **rep3** | 0.286 | 0.657 | | 0.741 | 0.824 | 0.899 | 0.728 |
| | **rep4** | - | - | | 0.694 | 0.868 | | 0.888 |
| **15** | **rep1** | 2.284 | 2.342 | 0.234 | 0.981 | 0.974 | 0.983 | 0.901 |
| | **rep2** | 2.159 | 2.472 | | 0.974 | 1.085 | | 0.919 |
| | **rep3** | 1.722 | 1.867 | | 1.178 | 0.878 | | 0.794 |
| | **rep4** | 2.060 | 2.609 | | 0.941 | 1.129 | | 0.928 |
| **16** | **rep1** | 0.373 | 0.359 | 0.261 | 0.694 | 0.701 | **0.007** | 0.723 |
| | **rep2** | 0.495 | 0.690 | | 0.835 | 0.571 | | 0.668 |
| | **rep3** | 0.529 | 0.779 | | 0.860 | 0.510 | 1.686 | 0.694 |

| | | | | | | | | |
|---|---|---|---|---|---|---|---|---|
| | **rep4** | 0.425 | 0.495 | | 0.859 | 0.505 | | 0.630 |
| **17** | **rep1** | 0.440 | 0.451 | 0.137 | 1.188 | 0.904 | 0.699 | 0.655 |
| | **rep2** | 0.570 | 0.584 | | 1.524 | 1.262 | | 0.426 |
| | **rep3** | 0.515 | 0.646 | | 1.533 | 1.458 | | 0.416 |
| | **rep4** | 0.416 | 0.694 | | 1.056 | 1.397 | | - |
| **23** | **rep1** | 0.908 | 0.819 | 0.093 | 2.512 | 2.976 | 0.649 | 1.830 |
| | **rep2** | 1.031 | 0.886 | | 2.947 | 2.730 | | 2.143 |
| | **rep3** | 1.123 | 0.915 | | 2.731 | 2.508 | | 1.659 |
| | **rep4** | 1.331 | - | | 2.413 | - | | 1.686 |
| **25** | **rep1** | 5.070 | 4.181 | 0.776 | 1.878 | 1.272 | 0.422 | 1.898 |
| | **rep2** | 4.048 | 4.625 | | 1.453 | 1.556 | | 2.186 |
| | **rep3** | 4.827 | 4.495 | | 1.792 | 1.774 | | 1.764 |
| | **rep4** | 3.609 | 4.678 | | - | - | | - |
| **26** | **rep1** | 1.263 | 1.613 | **0.013** | 1.515 | 2.097 | 0.726 | 0.954 |
| | **rep2** | 1.286 | 1.637 | | 2.028 | 1.978 | | 1.100 |
| | **rep3** | 1.239 | 1.655 | | 1.646 | 1.696 | | 1.333 |
| | **rep4** | 1.592 | 1.838 | | 2.191 | - | | 1.238 |
| **28** | **rep1** | 13.440 | 14.832 | 0.671 | 7.918 | 8.605 | 0.756 | 4.334 |
| | **rep2** | 20.553 | 18.519 | | 7.646 | 6.327 | | 4.081 |
| | **rep3** | 18.752 | 14.037 | | 7.553 | 9.029 | | 3.599 |
| | **rep4** | - | 18.780 | | - | - | | 3.474 |
| **33** | **rep1** | 0.703 | 0.877 | **0.033** | 0.798 | 1.032 | 0.775 | 0.817 |
| | **rep2** | 0.732 | 0.902 | | 1.011 | 1.034 | | 0.851 |
| | **rep3** | 0.723 | 1.118 | | 1.012 | 0.842 | | 1.041 |
| | **rep4** | - | - | | - | - | | - |
| **36** | **rep1** | 1.583 | 1.429 | 0.303 | 1.167 | 1.163 | 0.674 | 0.905 |
| | **rep2** | 1.424 | 1.274 | | 1.182 | 1.384 | | 0.993 |
| | **rep3** | - | - | | 1.337 | 1.201 | | 0.993 |
| | **rep4** | - | - | | 1.452 | 1.242 | | 0.976 |
| **37** | **rep1** | 1.244 | 1.446 | 0.230 | 1.857 | 2.274 | 0.672 | 5.679 |
| | **rep2** | 1.610 | 1.593 | | 2.153 | 1.623 | | 5.587 |
| | **rep3** | 1.498 | 1.647 | | 3.516 | 2.791 | | 5.929 |
| | **rep4** | 1.125 | - | | - | - | | 6.114 |
| **38** | **rep1** | 4.706 | 5.714 | 0.274 | 0.762 | 0.646 | 0.052 | 0.957 |
| | **rep2** | 5.815 | 5.604 | | 1.215 | 0.699 | | 0.863 |
| | **rep3** | 4.724 | 4.853 | | 1.088 | 0.868 | | 1.023 |
| | **rep4** | 5.224 | 6.047 | | 0.916 | 0.701 | | - |

| Neurons | |
|---|---|
| EUR | t-test |
| 2.006 | **0.0005** |
| 1.781 | |
| 1.973 | |
| - | |
| 1.180 | **0.00001** |
| 1.225 | |
| 1.240 | |
| 1.202 | |
| 1.090 | 0.214 |
| 1.044 | |
| 0.983 | |
| 1.140 | |
| 2.384 | 0.286 |
| 3.121 | |
| 3.089 | |
| - | |
| 1.369 | 0.071 |
| 1.342 | |
| 1.373 | |
| 1.405 | |
| 281.780 | **0.0000001** |
| 283.194 | |
| 271.135 | |
| 267.642 | |
| 1.956 | **0.009** |
| 1.979 | |
| 1.744 | |
| 2.101 | |
| 0.719 | 0.187 |
| 0.641 | |
| 0.686 | |
| - | |
| 0.849 | 0.726 |
| 0.819 | |
| 0.801 | |
| 0.812 | |
| 0.887 | 0.267 |
| 0.814 | |
| 0.721 | |
| 0.878 | |
| 0.576 | 0.569 |
| 0.663 | |
| 0.611 | |

| | |
|---|---|
| 0.759 | |
| 0.438 | 0.815 |
| 0.479 | |
| 0.519 | |
| - | |
| 1.660 | 0.550 |
| 1.854 | |
| 1.789 | |
| 1.714 | |
| 2.297 | 0.104 |
| 2.104 | |
| 2.397 | |
| - | |
| 0.877 | 0.068 |
| 0.955 | |
| 1.045 | |
| 0.951 | |
| 4.198 | 0.356 |
| 3.879 | |
| 4.349 | |
| - | |
| 1.012 | 0.881 |
| 0.839 | |
| 0.813 | |
| - | |
| 0.945 | 0.585 |
| 0.915 | |
| 0.967 | |
| 1.196 | |
| 2.728 | **0.000003** |
| 3.122 | |
| 3.248 | |
| 2.917 | |
| 0.989 | 0.114 |
| 1.185 | |
| 1.126 | |
| 1.009 | |

**Suppl Table 1: probe-based MPRA results**

| | Allele (APOEε4 most common) | | | | | |
|---|---|---|---|---|---|---|
| **Position** | **A_allele** | **R_allele** | **DNA_A** | **DNA_R** | **A/(A+R)** | **Experiment** |
| **Variants with significant results** | | | | | | |
| rs34278513 | | | 1161 | 1107 | | astocytes-rep1 |
| rs34278513 | T (EUR/JPT) | C (AFR) | 1161 | 1107 | 0.511905 | astocytes-rep2 |
| rs34278513 | | | 1161 | 1107 | | astocytes-rep3 |
| rs3865427 | | | 654 | 545 | | astocytes-rep1 |
| rs3865427 | A (EUR/JPT) | C (AFR) | 654 | 545 | 0.545455 | astocytes-rep2 |
| rs3865427 | | | 654 | 545 | | astocytes-rep3 |
| rs71352237 | | | 440 | 1522 | | astocytes-rep1 |
| rs71352237 | C (EUR/JPT) | T (AFR) | 440 | 1522 | 0.224261 | astocytes-rep2 |
| rs71352237 | | | 440 | 1522 | | astocytes-rep3 |
| rs283808 | | | 2869 | 733 | | astocytes-rep1 |
| rs283808 | C (AFR) | A (EUR/JPT) | 2869 | 733 | 0.796502 | astocytes-rep2 |
| rs283808 | | | 2869 | 733 | | astocytes-rep3 |
| rs12972970 | | | 4350 | 3156 | | astocytes-rep1 |
| rs12972970 | A (EUR/JPT) | G (AFR) | 4350 | 3156 | 0.579536 | astocytes-rep2 |
| rs12972970 | | | 4350 | 3156 | | astocytes-rep3 |
| rs34342646 | | | 1037 | 1397 | | microglia-rep1 |
| rs34342646 | A (EUR/JPT) | G (AFR) | 1037 | 1397 | 0.426048 | microglia-rep2 |
| rs34342646 | | | 1037 | 1397 | | microglia-rep3 |
| rs34342646 | | | 892 | 1222 | | astocytes-rep1 |
| rs34342646 | A (EUR/JPT) | G (AFR) | 892 | 1222 | 0.421949 | astocytes-rep2 |
| rs34342646 | | | 892 | 1222 | | astocytes-rep3 |
| rs71352238 | | | 625 | 1200 | | astocytes-rep1 |
| rs71352238 | C (EUR/JPT) | T (AFR) | 625 | 1200 | 0.342466 | astocytes-rep2 |
| rs71352238 | | | 625 | 1200 | | astocytes-rep3 |
| rs34404554 | | | 1167 | 1152 | | astocytes-rep1 |
| rs34404554 | G (EUR/JPT) | C (AFR) | 1167 | 1152 | 0.503234 | astocytes-rep2 |
| rs34404554 | | | 1167 | 1152 | | astocytes-rep3 |
| rs59007384 | | | 3003 | 1827 | | microglia-rep1 |
| rs59007384 | T (EUR/JPT) | G (AFR) | 3003 | 1827 | 0.621739 | microglia-rep2 |
| rs59007384 | | | 3003 | 1827 | | microglia-rep3 |
| rs449647 | | | 1232 | 1910 | | astocytes-rep1 |
| rs449647 | T (AFR) | A (EUR/JPT) | 1232 | 1910 | 0.392107 | astocytes-rep2 |
| rs449647 | | | 1232 | 1910 | | astocytes-rep3 |
| rs75627662 | | | 568 | 255 | | astocytes-rep1 |
| rs75627662 | T (EUR/JPT) | C (AFR) | 568 | 255 | 0.690158 | astocytes-rep2 |
| rs75627662 | | | 568 | 255 | | astocytes-rep3 |
| rs12721046 | | | 1489 | 1463 | | astocytes-rep1 |
| rs12721046 | A (EUR/JPT) | G (AFR) | 1489 | 1463 | 0.504404 | astocytes-rep2 |
| rs12721046 | | | 1489 | 1463 | | astocytes-rep3 |
| **Variants with negative results** | | | | | | |
| rs34278513 | | | 1417 | 1348 | | microglia-rep1 |
| rs34278513 | C (AFR) | T (EUR/JPT) | 1417 | 1348 | 0.512477 | microglia-rep2 |

| | | | | | | |
|---|---|---|---|---|---|---|
| rs34278513 | | | 1417 | 1348 | | microglia-rep3 |
| rs34278513 | | | 1161 | 1107 | | neurons-rep1 |
| rs34278513 | C (AFR) | T (EUR/JPT) | 1161 | 1107 | 0.511905 | neurons-rep2 |
| rs34278513 | | | 1161 | 1107 | | neurons-rep3 |
| rs3865427 | | | 779 | 601 | | microglia-rep1 |
| rs3865427 | C (AFR) | A (EUR/JPT) | 779 | 601 | 0.564493 | microglia-rep2 |
| rs3865427 | | | 779 | 601 | | microglia-rep3 |
| rs3865427 | | | 654 | 545 | | neurons-rep1 |
| rs3865427 | C (AFR) | A (EUR/JPT) | 654 | 545 | 0.545455 | neurons-rep2 |
| rs3865427 | | | 654 | 545 | | neurons-rep3 |
| rs34224078 | | | 2132 | 821 | | microglia-rep1 |
| rs34224078 | G (EUR/JPT) | A (AFR) | 2132 | 821 | 0.721978 | microglia-rep2 |
| rs34224078 | | | 2132 | 821 | | microglia-rep3 |
| rs34224078 | | | 1776 | 654 | | astocytes-rep1 |
| rs34224078 | G (EUR/JPT) | A (AFR) | 1776 | 654 | 0.730864 | astocytes-rep2 |
| rs34224078 | | | 1776 | 654 | | astocytes-rep3 |
| rs34224078 | | | 1776 | 654 | | neurons-rep1 |
| rs34224078 | G (EUR/JPT) | A (AFR) | 1776 | 654 | 0.730864 | neurons-rep2 |
| rs34224078 | | | 1776 | 654 | | neurons-rep3 |
| rs35879138 | | | 2324 | 1683 | | microglia-rep1 |
| rs35879138 | A (EUR/JPT) | T (AFR) | 2324 | 1683 | 0.579985 | microglia-rep2 |
| rs35879138 | | | 2324 | 1683 | | microglia-rep3 |
| rs35879138 | | | 1982 | 1402 | | astocytes-rep1 |
| rs35879138 | A (EUR/JPT) | T (AFR) | 1982 | 1402 | 0.585697 | astocytes-rep2 |
| rs35879138 | | | 1982 | 1402 | | astocytes-rep3 |
| rs35879138 | | | 1982 | 1402 | | neurons-rep1 |
| rs35879138 | A (EUR/JPT) | T (AFR) | 1982 | 1402 | 0.585697 | neurons-rep2 |
| rs35879138 | | | 1982 | 1402 | | neurons-rep3 |
| rs71352237 | | | 575 | 1692 | | microglia-rep1 |
| rs71352237 | T (AFR) | C (EUR/JPT) | 575 | 1692 | 0.253639 | microglia-rep2 |
| rs71352237 | | | 575 | 1692 | | microglia-rep3 |
| rs71352237 | | | 440 | 1522 | | neurons-rep1 |
| rs71352237 | T (AFR) | C (EUR/JPT) | 440 | 1522 | 0.224261 | neurons-rep2 |
| rs71352237 | | | 440 | 1522 | | neurons-rep3 |
| rs283808 | | | 3231 | 747 | | microglia-rep1 |
| rs283808 | C (AFR) | A (EUR/JPT) | 3231 | 747 | 0.812217 | microglia-rep2 |
| rs283808 | | | 3231 | 747 | | microglia-rep3 |
| rs283808 | | | 2869 | 733 | | neurons-rep1 |
| rs283808 | C (AFR) | A (EUR/JPT) | 2869 | 733 | 0.796502 | neurons-rep2 |
| rs283808 | | | 2869 | 733 | | neurons-rep3 |
| rs283809 | | | 2106 | 706 | | microglia-rep1 |
| rs283809 | G (AFR) | A (EUR/JPT) | 2106 | 706 | 0.748933 | microglia-rep2 |
| rs283809 | | | 2106 | 706 | | microglia-rep3 |
| rs283809 | | | 1746 | 589 | | astocytes-rep1 |
| rs283809 | G (AFR) | A (EUR/JPT) | 1746 | 589 | 0.747752 | astocytes-rep2 |
| rs283809 | | | 1746 | 589 | | astocytes-rep3 |
| rs283809 | | | 1746 | 589 | | neurons-rep1 |

| | | | | | | |
|---|---|---|---|---|---|---|
| rs283809 | G (AFR) | A (EUR/JPT) | 1746 | 589 | 0.747752 | neurons-rep2 |
| rs283809 | | | 1746 | 589 | | neurons-rep3 |
| rs12972156 | | | 1429 | 2135 | | microglia-rep1 |
| rs12972156 | G (EUR/JPT) | C (AFR) | 1429 | 2135 | 0.400954 | microglia-rep2 |
| rs12972156 | | | 1429 | 2135 | | microglia-rep3 |
| rs12972156 | | | 1194 | 1854 | | astocytes-rep1 |
| rs12972156 | G (EUR/JPT) | C (AFR) | 1194 | 1854 | 0.391732 | astocytes-rep2 |
| rs12972156 | | | 1194 | 1854 | | astocytes-rep3 |
| rs12972156 | | | 1194 | 1854 | | neurons-rep1 |
| rs12972156 | G (EUR/JPT) | C (AFR) | 1194 | 1854 | 0.391732 | neurons-rep2 |
| rs12972156 | | | 1194 | 1854 | | neurons-rep3 |
| rs12972970 | | | 5030 | 3866 | | microglia-rep1 |
| rs12972970 | A (EUR/JPT) | G (AFR) | 5030 | 3866 | 0.565423 | microglia-rep2 |
| rs12972970 | | | 5030 | 3866 | | microglia-rep3 |
| rs12972970 | | | 4350 | 3156 | | neurons-rep1 |
| rs12972970 | A (EUR/JPT) | G (AFR) | 4350 | 3156 | 0.579536 | neurons-rep2 |
| rs12972970 | | | 4350 | 3156 | | neurons-rep3 |
| rs34342646 | | | 892 | 1222 | | neurons-rep1 |
| rs34342646 | A (EUR/JPT) | G (AFR) | 892 | 1222 | 0.421949 | neurons-rep2 |
| rs34342646 | | | 892 | 1222 | | neurons-rep3 |
| rs71352238 | | | 733 | 1379 | | microglia-rep1 |
| rs71352238 | C (EUR/JPT) | T (AFR) | 733 | 1379 | 0.347064 | microglia-rep2 |
| rs71352238 | | | 733 | 1379 | | microglia-rep3 |
| rs71352238 | | | 625 | 1200 | | neurons-rep1 |
| rs71352238 | C (EUR/JPT) | T (AFR) | 625 | 1200 | 0.342466 | neurons-rep2 |
| rs71352238 | | | 625 | 1200 | | neurons-rep3 |
| rs2075650 | | | 266 | 518 | | microglia-rep1 |
| rs2075650 | G (EUR/JPT) | A (AFR) | 266 | 518 | 0.339286 | microglia-rep2 |
| rs2075650 | | | 266 | 518 | | microglia-rep3 |
| rs2075650 | | | 222 | 407 | | astocytes-rep1 |
| rs2075650 | G (EUR/JPT) | A (AFR) | 222 | 407 | 0.352941 | astocytes-rep2 |
| rs2075650 | | | 222 | 407 | | astocytes-rep3 |
| rs2075650 | | | 222 | 407 | | neurons-rep1 |
| rs2075650 | G (EUR/JPT) | A (AFR) | 222 | 407 | 0.352941 | neurons-rep2 |
| rs2075650 | | | 222 | 407 | | neurons-rep3 |
| rs34095326 | | | 4593 | 3105 | | microglia-rep1 |
| rs34095326 | A (EUR/JPT) | G (AFR) | 4593 | 3105 | 0.596648 | microglia-rep2 |
| rs34095326 | | | 4593 | 3105 | | microglia-rep3 |
| rs34095326 | | | 4025 | 2561 | | astocytes-rep1 |
| rs34095326 | A (EUR/JPT) | G (AFR) | 4025 | 2561 | 0.611145 | astocytes-rep2 |
| rs34095326 | | | 4025 | 2561 | | astocytes-rep3 |
| rs34095326 | | | 4025 | 2561 | | neurons-rep1 |
| rs34095326 | A (EUR/JPT) | G (AFR) | 4025 | 2561 | 0.611145 | neurons-rep2 |
| rs34095326 | | | 4025 | 2561 | | neurons-rep3 |
| rs34404554 | | | 1327 | 1440 | | microglia-rep1 |
| rs34404554 | G (EUR/JPT) | C (AFR) | 1327 | 1440 | 0.479581 | microglia-rep2 |
| rs34404554 | | | 1327 | 1440 | | microglia-rep3 |

| | | | | | | |
|---|---|---|---|---|---|---|
| rs34404554 | | | 1167 | 1152 | | neurons-rep1 |
| rs34404554 | G (EUR/JPT) | C (AFR) | 1167 | 1152 | 0.503234 | neurons-rep2 |
| rs34404554 | | | 1167 | 1152 | | neurons-rep3 |
| rs11556505 | | | 358 | 464 | | microglia-rep1 |
| rs11556505 | T (EUR/JPT) | C (AFR) | 358 | 464 | 0.435523 | microglia-rep2 |
| rs11556505 | | | 358 | 464 | | microglia-rep3 |
| rs11556505 | | | 296 | 368 | | astocytes-rep1 |
| rs11556505 | T (EUR/JPT) | C (AFR) | 296 | 368 | 0.445783 | astocytes-rep2 |
| rs11556505 | | | 296 | 368 | | astocytes-rep3 |
| rs11556505 | | | 296 | 368 | | neurons-rep1 |
| rs11556505 | T (EUR/JPT) | C (AFR) | 296 | 368 | 0.445783 | neurons-rep2 |
| rs11556505 | | | 296 | 368 | | neurons-rep3 |
| rs59007384 | | | 2476 | 1412 | | astocytes-rep1 |
| rs59007384 | T (EUR/JPT) | G (AFR) | 2476 | 1412 | 0.636831 | astocytes-rep2 |
| rs59007384 | | | 2476 | 1412 | | astocytes-rep3 |
| rs59007384 | | | 2476 | 1412 | | neurons-rep1 |
| rs59007384 | T (EUR/JPT) | G (AFR) | 2476 | 1412 | 0.636831 | neurons-rep2 |
| rs59007384 | | | 2476 | 1412 | | neurons-rep3 |
| rs157583 | | | 3355 | 2934 | | microglia-rep1 |
| rs157583 | T (AFR) | G (EUR/JPT) | 3355 | 2934 | | microglia-rep2 |
| rs157583 | | | 3355 | 2934 | | microglia-rep3 |
| rs157583 | | | 2903 | 2459 | | astocytes-rep1 |
| rs157583 | T (AFR) | G (EUR/JPT) | 2903 | 2459 | 0.541402 | astocytes-rep2 |
| rs157583 | | | 2903 | 2459 | | astocytes-rep3 |
| rs157583 | | | 2903 | 2459 | | neurons-rep1 |
| rs157583 | T (AFR) | G (EUR/JPT) | 2903 | 2459 | 0.541402 | neurons-rep2 |
| rs157583 | | | 2903 | 2459 | | neurons-rep3 |
| rs157584 | | | 1422 | 2953 | | microglia-rep1 |
| rs157584 | C (AFR) | T (EUR/JPT) | 1422 | 2953 | 0.325029 | microglia-rep2 |
| rs157584 | | | 1422 | 2953 | | microglia-rep3 |
| rs157584 | | | 1212 | 2489 | | astocytes-rep1 |
| rs157584 | C (AFR) | T (EUR/JPT) | 1212 | 2489 | 0.327479 | astocytes-rep2 |
| rs157584 | | | 1212 | 2489 | | astocytes-rep3 |
| rs157584 | | | 1212 | 2489 | | neurons-rep1 |
| rs157584 | C (AFR) | T (EUR/JPT) | 1212 | 2489 | 0.327479 | neurons-rep2 |
| rs157584 | | | 1212 | 2489 | | neurons-rep3 |
| rs157585 | | | 1420 | 690 | | microglia-rep1 |
| rs157585 | C (AFR) | A (EUR/JPT) | 1420 | 690 | 0.672986 | microglia-rep2 |
| rs157585 | | | 1420 | 690 | | microglia-rep3 |
| rs157585 | | | 1188 | 565 | | astocytes-rep1 |
| rs157585 | C (AFR) | A (EUR/JPT) | 1188 | 565 | 0.677695 | astocytes-rep2 |
| rs157585 | | | 1188 | 565 | | astocytes-rep3 |
| rs157585 | | | 1188 | 565 | | neurons-rep1 |
| rs157585 | C (AFR) | A (EUR/JPT) | 1188 | 565 | 0.677695 | neurons-rep2 |
| rs157585 | | | 1188 | 565 | | neurons-rep3 |
| rs157588 | | | 81 | 140 | | microglia-rep1 |
| rs157588 | T (AFR) | C (EUR/JPT) | 81 | 140 | 0.366516 | microglia-rep2 |

| | | | | | | |
|---|---|---|---|---|---|---|
| rs157588 | | | 81 | 140 | | microglia-rep3 |
| rs157588 | | | 67 | 135 | | astocytes-rep1 |
| rs157588 | T (AFR) | C (EUR/JPT) | 67 | 135 | 0.331683 | astocytes-rep2 |
| rs157588 | | | 67 | 135 | | astocytes-rep3 |
| rs157588 | | | 67 | 135 | | neurons-rep1 |
| rs157588 | T (AFR) | C (EUR/JPT) | 67 | 135 | 0.331683 | neurons-rep2 |
| rs157588 | | | 67 | 135 | | neurons-rep3 |
| rs157590 | | | 4643 | 3568 | | microglia-rep1 |
| rs157590 | C (AFR) | A (EUR/JPT) | 4643 | 3568 | 0.565461 | microglia-rep2 |
| rs157590 | | | 4643 | 3568 | | microglia-rep3 |
| rs157590 | | | 4090 | 3140 | | astocytes-rep1 |
| rs157590 | C (AFR) | A (EUR/JPT) | 4090 | 3140 | 0.565698 | astocytes-rep2 |
| rs157590 | | | 4090 | 3140 | | astocytes-rep3 |
| rs157590 | | | 4090 | 3140 | | neurons-rep1 |
| rs157590 | C (AFR) | A (EUR/JPT) | 4090 | 3140 | 0.565698 | neurons-rep2 |
| rs157590 | | | 4090 | 3140 | | neurons-rep3 |
| rs205909 | | | 2514 | 2090 | | microglia-rep1 |
| rs205909 | G (AFR) | T (EUR/JPT) | 2514 | 2090 | 0.546047 | microglia-rep2 |
| rs205909 | | | 2514 | 2090 | | microglia-rep3 |
| rs205909 | | | 2083 | 1682 | | astocytes-rep1 |
| rs205909 | G (AFR) | T (EUR/JPT) | 2083 | 1682 | 0.553254 | astocytes-rep2 |
| rs205909 | | | 2083 | 1682 | | astocytes-rep3 |
| rs205909 | | | 2083 | 1682 | | neurons-rep1 |
| rs205909 | G (AFR) | T (EUR/JPT) | 2083 | 1682 | 0.553254 | neurons-rep2 |
| rs205909 | | | 2083 | 1682 | | neurons-rep3 |
| rs491153 | | | 541 | 157 | | microglia-rep1 |
| rs491153 | T (AFR) | C (EUR/JPT) | 541 | 157 | 0.775072 | microglia-rep2 |
| rs491153 | | | 541 | 157 | | microglia-rep3 |
| rs491153 | | | 422 | 127 | | astocytes-rep1 |
| rs491153 | T (AFR) | C (EUR/JPT) | 422 | 127 | 0.76867 | astocytes-rep2 |
| rs491153 | | | 422 | 127 | | astocytes-rep3 |
| rs491153 | | | 422 | 127 | | neurons-rep1 |
| rs491153 | T (AFR) | C (EUR/JPT) | 422 | 127 | 0.76867 | neurons-rep2 |
| rs491153 | | | 422 | 127 | | neurons-rep3 |
| rs490243 | | | 611 | 1623 | | microglia-rep1 |
| rs490243 | T (AFR) | C (EUR/JPT) | 611 | 1623 | 0.2735 | microglia-rep2 |
| rs490243 | | | 611 | 1623 | | microglia-rep3 |
| rs490243 | | | 484 | 1324 | | neurons-rep1 |
| rs490243 | T (AFR) | C (EUR/JPT) | 484 | 1324 | 0.267699 | neurons-rep2 |
| rs490243 | | | 484 | 1324 | | neurons-rep3 |
| rs490243 | | | 484 | 1324 | | astocytes-rep1 |
| rs490243 | T (AFR) | C (EUR/JPT) | 484 | 1324 | 0.267699 | astocytes-rep2 |
| rs490243 | | | 484 | 1324 | | astocytes-rep3 |
| rs394819 | | | 1216 | 1686 | | microglia-rep1 |
| rs394819 | T (AFR) | G (EUR/JPT) | 1216 | 1686 | 0.419021 | microglia-rep2 |
| rs394819 | | | 1216 | 1686 | | microglia-rep3 |
| rs394819 | | | 988 | 1305 | | astocytes-rep1 |

| | | | | | | |
|---|---|---|---|---|---|---|
| rs394819 | T (AFR) | G (EUR/JPT) | 988 | 1305 | 0.430877 | astocytes-rep2 |
| rs394819 | | | 988 | 1305 | | astocytes-rep3 |
| rs394819 | | | 988 | 1305 | | neurons-rep1 |
| rs394819 | T (AFR) | G (EUR/JPT) | 988 | 1305 | 0.430877 | neurons-rep2 |
| rs394819 | | | 988 | 1305 | | neurons-rep3 |
| rs435380 | | | 349 | 81 | | microglia-rep1 |
| rs435380 | A (AFR) | G (EUR/JPT) | 349 | 81 | 0.811628 | microglia-rep2 |
| rs435380 | | | 349 | 81 | | microglia-rep3 |
| rs435380 | | | 307 | 66 | | astocytes-rep1 |
| rs435380 | A (AFR) | G (EUR/JPT) | 307 | 66 | 0.823056 | astocytes-rep2 |
| rs435380 | | | 307 | 66 | | astocytes-rep3 |
| rs435380 | | | 307 | 66 | | neurons-rep1 |
| rs435380 | A (AFR) | G (EUR/JPT) | 307 | 66 | 0.823056 | neurons-rep2 |
| rs435380 | | | 307 | 66 | | neurons-rep3 |
| rs446037 | | | 2963 | 3386 | | microglia-rep1 |
| rs446037 | T (AFR) | G (EUR/JPT) | 2963 | 3386 | 0.466688 | microglia-rep2 |
| rs446037 | | | 2963 | 3386 | | microglia-rep3 |
| rs446037 | | | 2450 | 2789 | | astocytes-rep1 |
| rs446037 | T (AFR) | G (EUR/JPT) | 2450 | 2789 | 0.467646 | astocytes-rep2 |
| rs446037 | | | 2450 | 2789 | | astocytes-rep3 |
| rs446037 | | | 2450 | 2789 | | neurons-rep1 |
| rs446037 | T (AFR) | G (EUR/JPT) | 2450 | 2789 | 0.467646 | neurons-rep2 |
| rs446037 | | | 2450 | 2789 | | neurons-rep3 |
| rs434132 | | | 1599 | 1973 | | microglia-rep1 |
| rs434132 | G (AFR) | C (EUR/JPT) | 1599 | 1973 | 0.447648 | microglia-rep2 |
| rs434132 | | | 1599 | 1973 | | microglia-rep3 |
| rs434132 | | | 1359 | 1635 | | astocytes-rep1 |
| rs434132 | G (AFR) | C (EUR/JPT) | 1359 | 1635 | 0.453908 | astocytes-rep2 |
| rs434132 | | | 1359 | 1635 | | astocytes-rep3 |
| rs434132 | | | 1359 | 1635 | | neurons-rep1 |
| rs434132 | G (AFR) | C (EUR/JPT) | 1359 | 1635 | 0.453908 | neurons-rep2 |
| rs434132 | | | 1359 | 1635 | | neurons-rep3 |
| rs449647 | | | 1506 | 2291 | | microglia-rep1 |
| rs449647 | T (AFR) | A (EUR/JPT) | 1506 | 2291 | 0.396629 | microglia-rep2 |
| rs449647 | | | 1506 | 2291 | | microglia-rep3 |
| rs449647 | | | 1232 | 1910 | | neurons-rep1 |
| rs449647 | T (AFR) | A (EUR/JPT) | 1232 | 1910 | 0.392107 | neurons-rep2 |
| rs449647 | | | 1232 | 1910 | | neurons-rep3 |
| rs769449 | | | 833 | 2621 | | microglia-rep1 |
| rs769449 | A (EUR/JPT) | G (AFR) | 833 | 2621 | 0.24117 | microglia-rep2 |
| rs769449 | | | 833 | 2621 | | microglia-rep3 |
| rs769449 | | | 772 | 2298 | | astocytes-rep1 |
| rs769449 | A (EUR/JPT) | G (AFR) | 772 | 2298 | 0.251466 | astocytes-rep2 |
| rs769449 | | | 772 | 2298 | | astocytes-rep3 |
| rs769449 | | | 772 | 2298 | | neurons-rep1 |
| rs769449 | A (EUR/JPT) | G (AFR) | 772 | 2298 | 0.251466 | neurons-rep2 |
| rs769449 | | | 772 | 2298 | | neurons-rep3 |

| | | | | | | |
|---|---|---|---|---|---|---|
| rs75627662 | | | 638 | 273 | | microglia-rep1 |
| rs75627662 | T (EUR/JPT) | C (AFR) | 638 | 273 | 0.700329 | microglia-rep2 |
| rs75627662 | | | 638 | 273 | | microglia-rep3 |
| rs75627662 | | | 568 | 255 | | neurons-rep1 |
| rs75627662 | T (EUR/JPT) | C (AFR) | 568 | 255 | 0.690158 | neurons-rep2 |
| rs75627662 | | | 568 | 255 | | neurons-rep3 |
| rs445925 | | | 1916 | 930 | | microglia-rep1 |
| rs445925 | A (AFR) | G (EUR/JPT) | 1916 | 930 | 0.673226 | microglia-rep2 |
| rs445925 | | | 1916 | 930 | | microglia-rep3 |
| rs445925 | | | 1594 | 774 | | neurons-rep1 |
| rs445925 | A (AFR) | G (EUR/JPT) | 1594 | 774 | 0.673142 | neurons-rep2 |
| rs445925 | | | 1594 | 774 | | neurons-rep3 |
| rs445925 | | | 1594 | 774 | | astocytes-rep1 |
| rs445925 | A (AFR) | G (EUR/JPT) | 1594 | 774 | 0.673142 | astocytes-rep2 |
| rs445925 | | | 1594 | 774 | | astocytes-rep3 |
| rs10414043 | | | 2963 | 2694 | | microglia-rep1 |
| rs10414043 | A (EUR/JPT) | G (AFR) | 2963 | 2694 | 0.523776 | microglia-rep2 |
| rs10414043 | | | 2963 | 2694 | | microglia-rep3 |
| rs10414043 | | | 2540 | 2236 | | astocytes-rep1 |
| rs10414043 | A (EUR/JPT) | G (AFR) | 2540 | 2236 | 0.531826 | astocytes-rep2 |
| rs10414043 | | | 2540 | 2236 | | astocytes-rep3 |
| rs10414043 | | | 2540 | 2236 | | neurons-rep1 |
| rs10414043 | A (EUR/JPT) | G (AFR) | 2540 | 2236 | 0.531826 | neurons-rep2 |
| rs10414043 | | | 2540 | 2236 | | neurons-rep3 |
| rs7256200 | | | 695 | 195 | | microglia-rep1 |
| rs7256200 | T (EUR/JPT) | G (AFR) | 695 | 195 | 0.780899 | microglia-rep2 |
| rs7256200 | | | 695 | 195 | | microglia-rep3 |
| rs7256200 | | | 585 | 179 | | astocytes-rep1 |
| rs7256200 | T (EUR/JPT) | G (AFR) | 585 | 179 | 0.765707 | astocytes-rep2 |
| rs7256200 | | | 585 | 179 | | astocytes-rep3 |
| rs7256200 | | | 585 | 179 | | neurons-rep1 |
| rs7256200 | T (EUR/JPT) | G (AFR) | 585 | 179 | 0.765707 | neurons-rep2 |
| rs7256200 | | | 585 | 179 | | neurons-rep3 |
| rs390082 | | | 2960 | 1248 | | microglia-rep1 |
| rs390082 | G (AFR) | T (EUR/JPT) | 2960 | 1248 | 0.703422 | microglia-rep2 |
| rs390082 | | | 2960 | 1248 | | microglia-rep3 |
| rs390082 | | | 2389 | 1003 | | astocytes-rep1 |
| rs390082 | G (AFR) | T (EUR/JPT) | 2389 | 1003 | 0.704304 | astocytes-rep2 |
| rs390082 | | | 2389 | 1003 | | astocytes-rep3 |
| rs390082 | | | 2389 | 1003 | | neurons-rep1 |
| rs390082 | G (AFR) | T (EUR/JPT) | 2389 | 1003 | 0.704304 | neurons-rep2 |
| rs390082 | | | 2389 | 1003 | | neurons-rep3 |
| rs5117 | | | 1951 | 2356 | | microglia-rep1 |
| rs5117 | C (EUR/JPT) | T (AFR) | 1951 | 2356 | 0.452984 | microglia-rep2 |
| rs5117 | | | 1951 | 2356 | | microglia-rep3 |
| rs5117 | | | 1614 | 2088 | | astocytes-rep1 |
| rs5117 | C (EUR/JPT) | T (AFR) | 1614 | 2088 | 0.435981 | astocytes-rep2 |

| | | | | | | |
|---|---|---|---|---|---|---|
| rs5117 | | | 1614 | 2088 | | astocytes-rep3 |
| rs5117 | | | 1614 | 2088 | | neurons-rep1 |
| rs5117 | C (EUR/JPT) | T (AFR) | 1614 | 2088 | 0.435981 | neurons-rep2 |
| rs5117 | | | 1614 | 2088 | | neurons-rep3 |
| rs389261 | | | 906 | 1157 | | microglia-rep1 |
| rs389261 | A (AFR) | G (EUR/JPT) | 906 | 1157 | 0.439166 | microglia-rep2 |
| rs389261 | | | 906 | 1157 | | microglia-rep3 |
| rs389261 | | | 782 | 940 | | astocytes-rep1 |
| rs389261 | A (AFR) | G (EUR/JPT) | 782 | 940 | 0.454123 | astocytes-rep2 |
| rs389261 | | | 782 | 940 | | astocytes-rep3 |
| rs389261 | | | 782 | 940 | | neurons-rep1 |
| rs389261 | A (AFR) | G (EUR/JPT) | 782 | 940 | 0.454123 | neurons-rep2 |
| rs389261 | | | 782 | 940 | | neurons-rep3 |
| rs12721046 | | | 1785 | 1675 | | microglia-rep1 |
| rs12721046 | A (EUR/JPT) | G (AFR) | 1785 | 1675 | 0.515896 | microglia-rep2 |
| rs12721046 | | | 1785 | 1675 | | microglia-rep3 |
| rs12721046 | | | 1489 | 1463 | | neurons-rep1 |
| rs12721046 | A (EUR/JPT) | G (AFR) | 1489 | 1463 | 0.504404 | neurons-rep2 |
| rs12721046 | | | 1489 | 1463 | | neurons-rep3 |
| rs12721051 | | | 1885 | 3864 | | microglia-rep1 |
| rs12721051 | G (EUR/JPT) | C (AFR) | 1885 | 3864 | 0.327883 | microglia-rep2 |
| rs12721051 | | | 1885 | 3864 | | microglia-rep3 |
| rs12721051 | | | 1622 | 3204 | | astocytes-rep1 |
| rs12721051 | G (EUR/JPT) | C (AFR) | 1622 | 3204 | 0.336096 | astocytes-rep2 |
| rs12721051 | | | 1622 | 3204 | | astocytes-rep3 |
| rs12721051 | | | 1622 | 3204 | | neurons-rep1 |
| rs12721051 | G (EUR/JPT) | C (AFR) | 1622 | 3204 | 0.336096 | neurons-rep2 |
| rs12721051 | | | 1622 | 3204 | | neurons-rep3 |
| rs56131196 | | | 355 | 574 | | microglia-rep1 |
| rs56131196 | G (AFR) | A (EUR/JPT) | 355 | 574 | 0.382131 | microglia-rep2 |
| rs56131196 | | | 355 | 574 | | microglia-rep3 |
| rs56131196 | | | 316 | 467 | | astocytes-rep1 |
| rs56131196 | G (AFR) | A (EUR/JPT) | 316 | 467 | 0.403576 | astocytes-rep2 |
| rs56131196 | | | 316 | 467 | | astocytes-rep3 |
| rs56131196 | | | 316 | 467 | | neurons-rep1 |
| rs56131196 | G (AFR) | A (EUR/JPT) | 316 | 467 | 0.403576 | neurons-rep2 |
| rs56131196 | | | 316 | 467 | | neurons-rep3 |
| rs4420638 | | | 3540 | 4200 | | microglia-rep1 |
| rs4420638 | G (EUR/JPT) | A (AFR) | 3540 | 4200 | 0.457364 | microglia-rep2 |
| rs4420638 | | | 3540 | 4200 | | microglia-rep3 |
| rs4420638 | | | 3004 | 3567 | | astocytes-rep1 |
| rs4420638 | G (EUR/JPT) | A (AFR) | 3004 | 3567 | 0.45716 | astocytes-rep2 |
| rs4420638 | | | 3004 | 3567 | | astocytes-rep3 |
| rs4420638 | | | 3004 | 3567 | | neurons-rep1 |
| rs4420638 | G (EUR/JPT) | A (AFR) | 3004 | 3567 | 0.45716 | neurons-rep2 |
| rs4420638 | | | 3004 | 3567 | | neurons-rep3 |
| rs157591 | | | 140 | 1046 | | microglia-rep1 |

| | | | | | | |
|---|---|---|---|---|---|---|
| rs157591 | A (AFR) | G (EUR/JPT) | 140 | 1046 | 0.118044 | microglia-rep2 |
| rs157591 | | | 140 | 1046 | | microglia-rep3 |
| rs157591 | | | 112 | 891 | | astocytes-rep1 |
| rs157591 | A (AFR) | G (EUR/JPT) | 112 | 891 | 0.111665 | astocytes-rep2 |
| rs157591 | | | 112 | 891 | | astocytes-rep3 |
| rs157591 | | | 112 | 891 | | neurons-rep1 |
| rs157591 | A (AFR) | G (EUR/JPT) | 112 | 891 | 0.111665 | neurons-rep2 |
| rs157591 | | | 112 | 891 | | neurons-rep3 |
| rs157596 | | | 1258 | 979 | | microglia-rep1 |
| rs157596 | T (AFR) | C (EUR/JPT) | 1258 | 979 | 0.56236 | microglia-rep2 |
| rs157596 | | | 1258 | 979 | | microglia-rep3 |
| rs157596 | | | 979 | 747 | | astocytes-rep1 |
| rs157596 | T (AFR) | C (EUR/JPT) | 979 | 747 | 0.567207 | astocytes-rep2 |
| rs157596 | | | 979 | 747 | | astocytes-rep3 |
| rs157596 | | | 979 | 747 | | neurons-rep1 |
| rs157596 | T (AFR) | C (EUR/JPT) | 979 | 747 | 0.567207 | neurons-rep2 |
| rs157596 | | | 979 | 747 | | neurons-rep3 |
| rs111789331 | | | 111 | 658 | | microglia-rep1 |
| rs111789331 | A (EUR/JPT) | T (AFR) | 111 | 658 | 0.144343 | microglia-rep2 |
| rs111789331 | | | 111 | 658 | | microglia-rep3 |
| rs111789331 | | | 128 | 584 | | astocytes-rep1 |
| rs111789331 | A (EUR/JPT) | T (AFR) | 128 | 584 | 0.179775 | astocytes-rep2 |
| rs111789331 | | | 128 | 584 | | astocytes-rep3 |
| rs111789331 | | | 128 | 584 | | neurons-rep1 |
| rs111789331 | A (EUR/JPT) | T (AFR) | 128 | 584 | 0.179775 | neurons-rep2 |
| rs111789331 | | | 128 | 584 | | neurons-rep3 |
| rs157599 | | | 3495 | 1769 | | microglia-rep1 |
| rs157599 | G (AFR) | A (EUR/JPT) | 3495 | 1769 | 0.663944 | microglia-rep2 |
| rs157599 | | | 3495 | 1769 | | microglia-rep3 |
| rs157599 | | | 2881 | 1597 | | astocytes-rep1 |
| rs157599 | G (AFR) | A (EUR/JPT) | 2881 | 1597 | 0.643368 | astocytes-rep2 |
| rs157599 | | | 2881 | 1597 | | astocytes-rep3 |
| rs157599 | | | 2881 | 1597 | | neurons-rep1 |
| rs157599 | G (AFR) | A (EUR/JPT) | 2881 | 1597 | 0.643368 | neurons-rep2 |
| rs157599 | | | 2881 | 1597 | | neurons-rep3 |
| rs149345 | | | 5267 | 3266 | | microglia-rep1 |
| rs149345 | G (AFR) | T (EUR/JPT) | 5267 | 3266 | 0.617251 | microglia-rep2 |
| rs149345 | | | 5267 | 3266 | | microglia-rep3 |
| rs149345 | | | 4386 | 2732 | | astocytes-rep1 |
| rs149345 | G (AFR) | T (EUR/JPT) | 4386 | 2732 | 0.616184 | astocytes-rep2 |
| rs149345 | | | 4386 | 2732 | | astocytes-rep3 |
| rs149345 | | | 4386 | 2732 | | neurons-rep1 |
| rs149345 | G (AFR) | T (EUR/JPT) | 4386 | 2732 | 0.616184 | neurons-rep2 |
| rs149345 | | | 4386 | 2732 | | neurons-rep3 |
| rs66626994 | | | 3201 | 5700 | | microglia-rep1 |
| rs66626994 | A (EUR/JPT) | G (AFR) | 3201 | 5700 | 0.359623 | microglia-rep2 |
| rs66626994 | | | 3201 | 5700 | | microglia-rep3 |

| | | | | | | |
|---|---|---|---|---|---|---|
| rs66626994 | | | 2692 | 4965 | | astocytes-rep1 |
| rs66626994 | A (EUR/JPT) | G (AFR) | 2692 | 4965 | 0.351574 | astocytes-rep2 |
| rs66626994 | | | 2692 | 4965 | | astocytes-rep3 |
| rs66626994 | | | 2692 | 4965 | | neurons-rep1 |
| rs66626994 | A (EUR/JPT) | G (AFR) | 2692 | 4965 | 0.351574 | neurons-rep2 |
| rs66626994 | | | 2692 | 4965 | | neurons-rep3 |
| rs10424663 | | | 1362 | 1424 | | microglia-rep1 |
| rs10424663 | A (AFR) | G (EUR/JPT) | 1362 | 1424 | 0.488873 | microglia-rep2 |
| rs10424663 | | | 1362 | 1424 | | microglia-rep3 |
| rs10424663 | | | 1184 | 1194 | | astocytes-rep1 |
| rs10424663 | A (AFR) | G (EUR/JPT) | 1184 | 1194 | 0.497897 | astocytes-rep2 |
| rs10424663 | | | 1184 | 1194 | | astocytes-rep3 |
| rs10424663 | | | 1184 | 1194 | | neurons-rep1 |
| rs10424663 | A (AFR) | G (EUR/JPT) | 1184 | 1194 | 0.497897 | neurons-rep2 |
| rs10424663 | | | 1184 | 1194 | | neurons-rep3 |

| RNA_A | RNA_R | A/(A+R) | conf.low | conf.high | p.value |
|---|---|---|---|---|---|
| 1200 | 1005 | 0.544218 | 0.523156 | 0.565162 | **0.002480853** |
| 746 | 557 | 0.572525 | 0.545141 | 0.599581 | **1.18E-05** |
| 959 | 757 | 0.558858 | 0.53499 | 0.582524 | **0.00010037** |
| 521 | 350 | 0.598163 | 0.564743 | 0.630917 | **0.001740904** |
| 388 | 254 | 0.604361 | 0.565348 | 0.64241 | **0.002926925** |
| 478 | 248 | 0.658402 | 0.622626 | 0.692885 | **6.74E-10** |
| 426 | 1130 | 0.273779 | 0.251746 | 0.296669 | **4.99E-06** |
| 164 | 564 | 0.225275 | 0.195405 | 0.257389 | 0.929254144 |
| 304 | 806 | 0.273874 | 0.247817 | 0.301135 | **0.000115368** |
| 2637 | 502 | 0.840076 | 0.826779 | 0.852737 | **5.12E-10** |
| 1565 | 353 | 0.815954 | 0.797872 | 0.833064 | **0.035806254** |
| 2176 | 383 | 0.850332 | 0.835915 | 0.863943 | **2.80E-12** |
| 3657 | 3029 | 0.546964 | 0.534939 | 0.558948 | **7.54E-08** |
| 2670 | 2152 | 0.553712 | 0.539549 | 0.567811 | **0.000296723** |
| 3357 | 2609 | 0.562689 | 0.549989 | 0.575327 | **0.008717261** |
| 347 | 458 | 0.431056 | 0.396531 | 0.466087 | 0.77567446 |
| 416 | 762 | 0.353141 | 0.325819 | 0.381199 | **3.85E-07** |
| 303 | 551 | 0.354801 | 0.322675 | 0.387933 | **2.36E-05** |
| 694 | 1043 | 0.399539 | 0.376404 | 0.423015 | 0.061388435 |
| 438 | 859 | 0.337702 | 0.311971 | 0.364171 | **5.69E-10** |
| 427 | 1028 | 0.293471 | 0.270166 | 0.317612 | **4.67E-24** |
| 456 | 1100 | 0.293059 | 0.27053 | 0.316372 | **3.42E-05** |
| 383 | 984 | 0.280176 | 0.256497 | 0.304803 | **9.16E-07** |
| 420 | 1125 | 0.271845 | 0.249787 | 0.294773 | **2.92E-09** |
| 812 | 937 | 0.464265 | 0.440683 | 0.487967 | **0.001141754** |
| 464 | 748 | 0.382838 | 0.355373 | 0.410874 | **4.50E-17** |
| 699 | 997 | 0.412146 | 0.388595 | 0.436002 | **6.41E-14** |
| 869 | 548 | 0.613267 | 0.587344 | 0.63872 | 0.511060559 |
| 1355 | 684 | 0.664541 | 0.643577 | 0.685032 | **6.38E-05** |
| 1021 | 499 | 0.671711 | 0.64746 | 0.695296 | **5.16E-05** |
| 1168 | 1485 | 0.440256 | 0.421253 | 0.459392 | **4.79E-07** |
| 685 | 965 | 0.415152 | 0.391242 | 0.439364 | 0.058598032 |
| 988 | 1306 | 0.430689 | 0.410307 | 0.451248 | **0.000166772** |
| 602 | 165 | 0.784876 | 0.754076 | 0.813467 | **5.26E-09** |
| 347 | 145 | 0.705285 | 0.662833 | 0.745248 | 0.495038175 |
| 436 | 129 | 0.771681 | 0.734813 | 0.805681 | **1.83E-05** |
| 2053 | 1943 | 0.513764 | 0.498137 | 0.529371 | 0.241728716 |
| 11339 | 7464 | 0.603042 | 0.596007 | 0.610046 | **2.33E-162** |
| 7630 | 5522 | 0.58014 | 0.57165 | 0.588594 | **6.70E-68** |
| 445 | 379 | 0.540049 | 0.50532 | 0.574489 | 0.116866418 |
| 697 | 614 | 0.531655 | 0.504217 | 0.558951 | 0.167231725 |

| | | | | | |
|---|---|---|---|---|---|
| 508 | 498 | 0.50497 | 0.473601 | 0.53631 | 0.636310892 |
| 496 | 433 | 0.533907 | 0.501227 | 0.566373 | 0.189249277 |
| 410 | 337 | 0.548862 | 0.512375 | 0.584962 | **0.044137062** |
| 353 | 321 | 0.523739 | 0.485258 | 0.562011 | 0.563336599 |
| 198 | 159 | 0.554622 | 0.501393 | 0.606937 | 0.709024921 |
| 317 | 227 | 0.582721 | 0.540008 | 0.62453 | 0.411422139 |
| 227 | 139 | 0.620219 | 0.568319 | 0.670156 | **0.034849679** |
| 208 | 181 | 0.534704 | 0.483749 | 0.585128 | 0.683954405 |
| 166 | 147 | 0.530351 | 0.473402 | 0.586721 | 0.609663801 |
| 134 | 136 | 0.496296 | 0.435134 | 0.557541 | 0.112041269 |
| 669 | 240 | 0.735974 | 0.706032 | 0.764377 | 0.354969414 |
| 939 | 359 | 0.723421 | 0.698214 | 0.747611 | 0.925977048 |
| 741 | 277 | 0.727898 | 0.699435 | 0.755036 | 0.700437274 |
| 1747 | 591 | 0.74722 | 0.729086 | 0.764732 | 0.076379117 |
| 1023 | 317 | 0.763433 | 0.739742 | 0.785962 | **0.007344758** |
| 1371 | 493 | 0.735515 | 0.714864 | 0.755423 | 0.676094239 |
| 689 | 272 | 0.716961 | 0.687325 | 0.745262 | 0.326395165 |
| 620 | 224 | 0.734597 | 0.70343 | 0.764116 | 0.84615472 |
| 582 | 191 | 0.752911 | 0.720928 | 0.782951 | 0.180762663 |
| 726 | 508 | 0.588331 | 0.560283 | 0.615956 | 0.564158599 |
| 974 | 768 | 0.559127 | 0.535442 | 0.582613 | 0.08051433 |
| 734 | 551 | 0.571206 | 0.543621 | 0.598465 | 0.534148743 |
| 1721 | 1199 | 0.589384 | 0.571287 | 0.607301 | 0.693268558 |
| 971 | 752 | 0.563552 | 0.539753 | 0.587134 | 0.063114734 |
| 1304 | 1078 | 0.547439 | 0.527193 | 0.567569 | **0.000165911** |
| 742 | 489 | 0.602762 | 0.574802 | 0.63023 | 0.23555771 |
| 638 | 462 | 0.58 | 0.550197 | 0.609374 | 0.713461143 |
| 575 | 423 | 0.576152 | 0.544805 | 0.607049 | 0.541701497 |
| 188 | 568 | 0.248677 | 0.218234 | 0.281095 | 0.770030189 |
| 233 | 734 | 0.240951 | 0.214302 | 0.269188 | 0.37529233 |
| 160 | 576 | 0.217391 | 0.188095 | 0.248973 | **0.024672988** |
| 170 | 531 | 0.242511 | 0.211209 | 0.275997 | 0.257536018 |
| 133 | 510 | 0.206843 | 0.176163 | 0.240241 | 0.298675368 |
| 136 | 423 | 0.243292 | 0.208264 | 0.281057 | 0.286918346 |
| 1156 | 214 | 0.843796 | 0.823472 | 0.862632 | **0.002596356** |
| 1509 | 307 | 0.830947 | 0.812903 | 0.847915 | **0.041036417** |
| 1178 | 245 | 0.827829 | 0.807191 | 0.847103 | 0.13536216 |
| 1136 | 247 | 0.821403 | 0.800179 | 0.841251 | **0.021173737** |
| 907 | 181 | 0.83364 | 0.810157 | 0.855302 | **0.002008086** |
| 808 | 168 | 0.827869 | 0.802694 | 0.851047 | **0.015233757** |
| 600 | 240 | 0.714286 | 0.682425 | 0.744634 | **0.023258225** |
| 945 | 322 | 0.745856 | 0.720935 | 0.769631 | 0.79556613 |
| 690 | 246 | 0.737179 | 0.707729 | 0.765129 | 0.407090575 |
| 1607 | 578 | 0.735469 | 0.716438 | 0.753867 | 0.191734802 |
| 876 | 335 | 0.723369 | 0.69724 | 0.748409 | 0.054928193 |
| 1357 | 490 | 0.734705 | 0.713938 | 0.754724 | 0.198508874 |
| 673 | 219 | 0.754484 | 0.724867 | 0.78241 | 0.671583084 |

| | | | | | |
|---|---|---|---|---|---|
| 636 | 209 | 0.752663 | 0.722129 | 0.781423 | 0.781597659 |
| 443 | 176 | 0.71567 | 0.678355 | 0.750914 | 0.070996833 |
| 400 | 726 | 0.35524 | 0.327251 | 0.383987 | **0.001729219** |
| 661 | 1098 | 0.375782 | 0.353087 | 0.398892 | **0.032277767** |
| 469 | 785 | 0.374003 | 0.347146 | 0.401453 | 0.053538339 |
| 991 | 1760 | 0.360233 | 0.342268 | 0.378495 | **0.000725757** |
| 580 | 1200 | 0.325843 | 0.30409 | 0.34817 | **9.55E-09** |
| 919 | 1535 | 0.374491 | 0.355297 | 0.393984 | 0.082393484 |
| 386 | 742 | 0.342199 | 0.31451 | 0.370713 | **0.000630787** |
| 414 | 684 | 0.377049 | 0.34829 | 0.40647 | 0.32275395 |
| 346 | 645 | 0.349142 | 0.319444 | 0.37974 | **0.006243974** |
| 1565 | 1262 | 0.55359 | 0.535041 | 0.572029 | 0.210533266 |
| 2341 | 1865 | 0.556586 | 0.541418 | 0.571675 | 0.249779371 |
| 1814 | 1410 | 0.562655 | 0.545326 | 0.57987 | 0.76266009 |
| 1599 | 1315 | 0.54873 | 0.53045 | 0.566913 | **0.000781527** |
| 1406 | 1059 | 0.570385 | 0.550567 | 0.590037 | 0.358649324 |
| 1270 | 987 | 0.562694 | 0.541937 | 0.583288 | 0.105202587 |
| 325 | 502 | 0.392987 | 0.359527 | 0.427212 | 0.097937446 |
| 232 | 425 | 0.35312 | 0.316544 | 0.391023 | **0.000370529** |
| 211 | 341 | 0.382246 | 0.341528 | 0.424233 | 0.063795582 |
| 235 | 445 | 0.345588 | 0.309846 | 0.382678 | 0.967875574 |
| 295 | 648 | 0.312831 | 0.283332 | 0.343505 | **0.028536011** |
| 279 | 481 | 0.367105 | 0.332751 | 0.402495 | 0.253044573 |
| 235 | 497 | 0.321038 | 0.287313 | 0.356214 | 0.227492104 |
| 178 | 364 | 0.328413 | 0.288982 | 0.369728 | 0.526336207 |
| 173 | 352 | 0.329524 | 0.289422 | 0.371558 | 0.550175446 |
| 60 | 147 | 0.289855 | 0.229052 | 0.356789 | 0.142241982 |
| 95 | 179 | 0.346715 | 0.29047 | 0.406317 | 0.798754719 |
| 74 | 142 | 0.342593 | 0.279547 | 0.410027 | 0.942738875 |
| 106 | 330 | 0.243119 | 0.203568 | 0.286196 | **1.04E-06** |
| 168 | 162 | 0.509091 | 0.453767 | 0.564251 | **6.54E-09** |
| 139 | 332 | 0.295117 | 0.254263 | 0.338565 | **0.009152301** |
| 63 | 141 | 0.308824 | 0.246177 | 0.377126 | 0.212847895 |
| 68 | 122 | 0.357895 | 0.289807 | 0.430493 | 0.87960241 |
| 47 | 117 | 0.286585 | 0.21878 | 0.362281 | 0.085874011 |
| 1532 | 904 | 0.6289 | 0.609364 | 0.648125 | **0.001182257** |
| 2209 | 1331 | 0.624011 | 0.607817 | 0.640001 | **0.00088787** |
| 1623 | 1072 | 0.602226 | 0.583459 | 0.620771 | 0.56911836 |
| 3480 | 2256 | 0.606695 | 0.593914 | 0.619366 | 0.489798768 |
| 2061 | 1213 | 0.629505 | 0.6127 | 0.646079 | **0.031468519** |
| 2796 | 1870 | 0.599228 | 0.585002 | 0.613331 | 0.095586628 |
| 1598 | 891 | 0.642025 | 0.622832 | 0.660883 | **0.001542102** |
| 1348 | 807 | 0.625522 | 0.604696 | 0.646006 | 0.177721298 |
| 1202 | 777 | 0.607377 | 0.585464 | 0.628973 | 0.729524892 |
| 414 | 427 | 0.492271 | 0.457966 | 0.526631 | 0.468732784 |
| 525 | 608 | 0.463372 | 0.434022 | 0.492912 | 0.28444812 |
| 414 | 425 | 0.493445 | 0.459093 | 0.527843 | 0.426935028 |

| | | | | | |
|---|---|---|---|---|---|
| 363 | 472 | 0.434731 | 0.40079 | 0.469134 | **7.81E-05** |
| 313 | 349 | 0.47281 | 0.434217 | 0.511646 | 0.12010082 |
| 313 | 330 | 0.486781 | 0.447507 | 0.526176 | 0.407844388 |
| 104 | 127 | 0.450216 | 0.384901 | 0.516825 | 0.690647796 |
| 174 | 195 | 0.471545 | 0.419664 | 0.523886 | 0.172271455 |
| 139 | 130 | 0.516729 | 0.455254 | 0.577831 | **0.008103696** |
| 222 | 336 | 0.397849 | 0.356977 | 0.43981 | **0.023945** |
| 133 | 259 | 0.339286 | 0.292509 | 0.388511 | **1.88E-05** |
| 214 | 235 | 0.476615 | 0.42959 | 0.52395 | 0.199901344 |
| 108 | 150 | 0.418605 | 0.357715 | 0.481387 | 0.415608852 |
| 75 | 110 | 0.405405 | 0.333989 | 0.479904 | 0.30047556 |
| 87 | 110 | 0.441624 | 0.371093 | 0.513939 | 0.942920085 |
| 2131 | 1133 | 0.65288 | 0.636264 | 0.669222 | 0.058390771 |
| 1515 | 791 | 0.656982 | 0.637199 | 0.676364 | **0.044078629** |
| 1767 | 1052 | 0.626818 | 0.608663 | 0.64471 | 0.272825869 |
| 886 | 568 | 0.609354 | 0.583731 | 0.634533 | **0.031202017** |
| 846 | 476 | 0.639939 | 0.613397 | 0.665858 | 0.841358787 |
| 712 | 417 | 0.630647 | 0.60174 | 0.65887 | 0.664974569 |
| 1051 | 942 | 0.527346 | 0.50515 | 0.549461 | 0.590056764 |
| 1429 | 1324 | 0.51907 | 0.500213 | 0.537887 | 0.131308351 |
| 1203 | 951 | 0.558496 | 0.537226 | 0.579606 | **0.020830572** |
| 2426 | 2127 | 0.532835 | 0.518216 | 0.547412 | 0.246111789 |
| 1362 | 1540 | 0.469331 | 0.451041 | 0.487684 | **8.36E-15** |
| 2237 | 1943 | 0.535167 | 0.519908 | 0.550378 | 0.419668005 |
| 1141 | 965 | 0.541785 | 0.520222 | 0.563232 | 0.982556619 |
| 951 | 837 | 0.531879 | 0.508435 | 0.555219 | 0.419865021 |
| 874 | 760 | 0.534884 | 0.510349 | 0.559293 | 0.602214307 |
| 410 | 1001 | 0.290574 | 0.266986 | 0.315037 | **0.005818647** |
| 687 | 1431 | 0.324363 | 0.304444 | 0.344768 | 0.963002046 |
| 496 | 1075 | 0.315722 | 0.292782 | 0.339353 | 0.450777342 |
| 1185 | 2283 | 0.341696 | 0.325905 | 0.357753 | 0.076213772 |
| 709 | 1164 | 0.378537 | 0.356509 | 0.400946 | **3.22E-06** |
| 917 | 2002 | 0.314149 | 0.297332 | 0.331339 | 0.12888479 |
| 431 | 972 | 0.307199 | 0.283127 | 0.33208 | 0.111140054 |
| 377 | 864 | 0.303787 | 0.278287 | 0.33022 | 0.079350735 |
| 326 | 786 | 0.293165 | 0.266537 | 0.320893 | **0.015134458** |
| 391 | 194 | 0.668376 | 0.628593 | 0.706449 | 0.825634987 |
| 535 | 270 | 0.664596 | 0.630803 | 0.697179 | 0.625318061 |
| 385 | 215 | 0.641667 | 0.601842 | 0.680089 | 0.107340525 |
| 856 | 418 | 0.6719 | 0.645351 | 0.697653 | 0.653121436 |
| 501 | 271 | 0.648964 | 0.614128 | 0.682656 | 0.090226017 |
| 795 | 376 | 0.678907 | 0.651314 | 0.705598 | 0.95015077 |
| 395 | 221 | 0.641234 | 0.601939 | 0.679167 | 0.057762959 |
| 340 | 152 | 0.691057 | 0.648152 | 0.731647 | 0.562750164 |
| 316 | 134 | 0.702222 | 0.657629 | 0.744133 | 0.289500806 |
| 21 | 47 | 0.308824 | 0.202363 | 0.432561 | 0.378815142 |
| 33 | 61 | 0.351064 | 0.255419 | 0.456376 | 0.830726431 |

| | | | | | |
|---|---|---|---|---|---|
| 25 | 30 | 0.454545 | 0.319701 | 0.594455 | 0.207429027 |
| 57 | 109 | 0.343373 | 0.271527 | 0.420926 | 0.742202213 |
| 24 | 46 | 0.342857 | 0.233482 | 0.465996 | 0.899061981 |
| 94 | 64 | 0.594937 | 0.514041 | 0.672202 | **1.40E-11** |
| 25 | 29 | 0.462963 | 0.326225 | 0.603905 | **0.043723815** |
| 24 | 47 | 0.338028 | 0.22997 | 0.460073 | 0.900217522 |
| 12 | 36 | 0.25 | 0.136372 | 0.395959 | 0.283436265 |
| 1377 | 1099 | 0.556139 | 0.53631 | 0.575835 | 0.351147184 |
| 2124 | 1763 | 0.546437 | 0.530628 | 0.562176 | **0.017382456** |
| 1632 | 1245 | 0.567258 | 0.548919 | 0.58546 | 0.850853873 |
| 3432 | 2808 | 0.55 | 0.537555 | 0.562398 | **0.012763179** |
| 2245 | 1669 | 0.573582 | 0.55791 | 0.589144 | 0.325339891 |
| 3094 | 2547 | 0.548484 | 0.535387 | 0.561531 | **0.009171337** |
| 1486 | 1128 | 0.568477 | 0.549231 | 0.58757 | 0.782398762 |
| 1267 | 991 | 0.561116 | 0.540358 | 0.581715 | 0.671160636 |
| 1099 | 892 | 0.551984 | 0.529826 | 0.573988 | 0.222167727 |
| 776 | 620 | 0.555874 | 0.529357 | 0.582155 | 0.468086113 |
| 1102 | 978 | 0.529808 | 0.508089 | 0.551443 | 0.140118141 |
| 815 | 679 | 0.545515 | 0.519864 | 0.570988 | 0.979273448 |
| 1805 | 1482 | 0.549133 | 0.531933 | 0.566245 | 0.635802669 |
| 1094 | 937 | 0.538651 | 0.516682 | 0.560508 | 0.187982337 |
| 1563 | 1081 | 0.59115 | 0.572129 | 0.609968 | **9.09E-05** |
| 785 | 700 | 0.52862 | 0.502859 | 0.554267 | 0.056780788 |
| 666 | 564 | 0.541463 | 0.51314 | 0.569589 | 0.40579522 |
| 548 | 498 | 0.523901 | 0.493125 | 0.554541 | 0.05783535 |
| 166 | 47 | 0.779343 | 0.7176 | 0.833135 | 0.934704048 |
| 225 | 47 | 0.827206 | 0.776931 | 0.870188 | **0.041762276** |
| 160 | 52 | 0.754717 | 0.691106 | 0.811057 | 0.460201578 |
| 372 | 100 | 0.788136 | 0.74846 | 0.824156 | 0.32645402 |
| 195 | 89 | 0.68662 | 0.629165 | 0.740129 | **0.001496985** |
| 335 | 105 | 0.761364 | 0.718717 | 0.800456 | 0.734450641 |
| 183 | 53 | 0.775424 | 0.716769 | 0.827018 | 0.877335334 |
| 137 | 46 | 0.748634 | 0.679315 | 0.809712 | 0.539363298 |
| 102 | 33 | 0.755556 | 0.674186 | 0.825375 | 0.684371863 |
| 195 | 525 | 0.270833 | 0.238674 | 0.304883 | 0.900231589 |
| 262 | 840 | 0.23775 | 0.212889 | 0.264019 | **0.007572529** |
| 210 | 557 | 0.273794 | 0.242506 | 0.306832 | 1 |
| 203 | 554 | 0.268164 | 0.2369 | 0.301246 | 0.967279195 |
| 170 | 422 | 0.287162 | 0.251011 | 0.325452 | 0.28587785 |
| 142 | 382 | 0.270992 | 0.233365 | 0.311225 | 0.882342724 |
| 429 | 1221 | 0.26 | 0.238975 | 0.281883 | 0.487188048 |
| 337 | 680 | 0.331367 | 0.302469 | 0.361244 | **7.80E-06** |
| 280 | 1008 | 0.217391 | 0.195143 | 0.240937 | **3.67E-05** |
| 376 | 572 | 0.396624 | 0.365319 | 0.428574 | 0.166907245 |
| 535 | 706 | 0.431104 | 0.403339 | 0.459196 | 0.388300874 |
| 413 | 535 | 0.435654 | 0.403805 | 0.467904 | 0.307611186 |
| 933 | 1172 | 0.44323 | 0.421863 | 0.464756 | 0.25255701 |

| | | | | | |
|---|---|---|---|---|---|
| 561 | 691 | 0.448083 | 0.420287 | 0.476123 | 0.219925685 |
| 708 | 1005 | 0.41331 | 0.389864 | 0.437053 | 0.143329 |
| 423 | 511 | 0.452891 | 0.420624 | 0.485456 | 0.175679149 |
| 362 | 449 | 0.446363 | 0.411781 | 0.481335 | 0.375601457 |
| 331 | 345 | 0.489645 | 0.451338 | 0.528043 | **0.002143838** |
| 87 | 19 | 0.820755 | 0.734325 | 0.888494 | 0.90139572 |
| 157 | 22 | 0.877095 | 0.81985 | 0.921349 | **0.027328051** |
| 90 | 30 | 0.75 | 0.662731 | 0.824535 | 0.100959113 |
| 204 | 27 | 0.883117 | 0.834512 | 0.921543 | **0.015547349** |
| 111 | 38 | 0.744966 | 0.667156 | 0.812772 | **0.017598616** |
| 150 | 23 | 0.867052 | 0.807218 | 0.91382 | 0.135888334 |
| 101 | 12 | 0.893805 | 0.821846 | 0.943911 | **0.04847126** |
| 101 | 18 | 0.848739 | 0.771517 | 0.90782 | 0.548099554 |
| 73 | 13 | 0.848837 | 0.755387 | 0.916983 | 0.67114865 |
| 914 | 1014 | 0.474066 | 0.451572 | 0.496639 | 0.522792529 |
| 1390 | 1463 | 0.487206 | 0.468711 | 0.505728 | **0.028147342** |
| 1021 | 1102 | 0.480923 | 0.459473 | 0.502426 | 0.19187688 |
| 2302 | 2472 | 0.482195 | 0.467931 | 0.496481 | **0.045330677** |
| 1302 | 1710 | 0.432271 | 0.414486 | 0.450187 | **0.000100041** |
| 1817 | 2099 | 0.463994 | 0.448278 | 0.479763 | 0.653891706 |
| 939 | 1117 | 0.456712 | 0.435012 | 0.478536 | 0.330844982 |
| 821 | 924 | 0.470487 | 0.446848 | 0.494225 | 0.829071993 |
| 797 | 809 | 0.496264 | 0.471525 | 0.521016 | **0.022845946** |
| 456 | 561 | 0.448378 | 0.417508 | 0.479547 | 0.974850424 |
| 653 | 842 | 0.436789 | 0.411458 | 0.462369 | 0.405367326 |
| 505 | 632 | 0.444151 | 0.415007 | 0.473585 | 0.834661586 |
| 1097 | 1395 | 0.440209 | 0.420598 | 0.45996 | 0.171346805 |
| 617 | 717 | 0.462519 | 0.435496 | 0.489707 | 0.52726913 |
| 917 | 908 | 0.502466 | 0.479264 | 0.525659 | **3.47E-05** |
| 524 | 668 | 0.439597 | 0.41118 | 0.468313 | 0.32284827 |
| 410 | 539 | 0.432034 | 0.400243 | 0.464247 | 0.181381722 |
| 375 | 460 | 0.449102 | 0.414996 | 0.483568 | 0.807805355 |
| 493 | 653 | 0.430192 | 0.401301 | 0.459442 | **0.021715201** |
| 688 | 977 | 0.413213 | 0.389432 | 0.437301 | 0.168350149 |
| 532 | 707 | 0.429379 | 0.401609 | 0.457484 | **0.020142054** |
| 530 | 686 | 0.435855 | 0.407757 | 0.464264 | **0.002031174** |
| 429 | 604 | 0.415295 | 0.385039 | 0.446035 | 0.134184641 |
| 411 | 565 | 0.421107 | 0.389897 | 0.452794 | 0.066355907 |
| 271 | 824 | 0.247489 | 0.222178 | 0.274164 | 0.621042336 |
| 374 | 1143 | 0.246539 | 0.225029 | 0.269035 | 0.631125392 |
| 305 | 811 | 0.273297 | 0.247329 | 0.300467 | **0.012973171** |
| 607 | 1881 | 0.243971 | 0.227204 | 0.261343 | 0.405522919 |
| 423 | 1219 | 0.257613 | 0.236604 | 0.279491 | 0.569491813 |
| 555 | 1401 | 0.283742 | 0.263845 | 0.30429 | **0.001122064** |
| 266 | 868 | 0.234568 | 0.210179 | 0.260341 | 0.193568677 |
| 237 | 778 | 0.233498 | 0.207782 | 0.260768 | 0.192904008 |
| 203 | 682 | 0.229379 | 0.202053 | 0.258519 | 0.140884366 |

| | | | | | |
|---|---|---|---|---|---|
| 211 | 73 | 0.742958 | 0.687997 | 0.79277 | 0.120413565 |
| 311 | 140 | 0.689579 | 0.644627 | 0.732022 | 0.607709462 |
| 269 | 90 | 0.749304 | 0.70113 | 0.793312 | **0.043742237** |
| 241 | 101 | 0.704678 | 0.65324 | 0.752531 | 0.598854549 |
| 193 | 71 | 0.731061 | 0.673274 | 0.783577 | 0.162294996 |
| 184 | 78 | 0.70229 | 0.642942 | 0.756994 | 0.738397069 |
| 615 | 290 | 0.679558 | 0.648055 | 0.709887 | 0.696833366 |
| 916 | 401 | 0.69552 | 0.66987 | 0.720294 | 0.088388211 |
| 731 | 260 | 0.737639 | 0.709065 | 0.764793 | **1.21E-05** |
| 714 | 279 | 0.719033 | 0.689952 | 0.746809 | **0.002064508** |
| 573 | 242 | 0.703067 | 0.670391 | 0.734268 | 0.073010525 |
| 484 | 188 | 0.720238 | 0.684638 | 0.753891 | **0.009525134** |
| 1501 | 530 | 0.739045 | 0.719363 | 0.758034 | **1.16E-10** |
| 807 | 445 | 0.644569 | 0.617341 | 0.671115 | **0.032422** |
| 1232 | 514 | 0.705613 | 0.683615 | 0.726917 | **0.003631926** |
| 1041 | 837 | 0.554313 | 0.531497 | 0.576959 | **0.008432358** |
| 1371 | 1170 | 0.539551 | 0.519938 | 0.559073 | 0.112128728 |
| 1074 | 849 | 0.558502 | 0.535973 | 0.580853 | **0.002386641** |
| 2293 | 2005 | 0.533504 | 0.518455 | 0.548508 | 0.830572368 |
| 1301 | 1220 | 0.516065 | 0.49635 | 0.535743 | 0.114888703 |
| 1877 | 1476 | 0.559797 | 0.5428 | 0.57669 | **0.001209135** |
| 1113 | 838 | 0.570477 | 0.548163 | 0.592578 | **0.000611984** |
| 936 | 728 | 0.5625 | 0.538271 | 0.586508 | **0.012221399** |
| 833 | 656 | 0.559436 | 0.533787 | 0.584849 | **0.033222711** |
| 227 | 65 | 0.777397 | 0.725275 | 0.823793 | 0.887508479 |
| 326 | 90 | 0.783654 | 0.740934 | 0.822285 | 0.952738536 |
| 263 | 76 | 0.775811 | 0.727629 | 0.819101 | 0.843827462 |
| 515 | 112 | 0.821372 | 0.7891 | 0.85058 | **0.00080285** |
| 312 | 74 | 0.80829 | 0.765415 | 0.846361 | **0.047493899** |
| 448 | 152 | 0.746667 | 0.709871 | 0.781014 | 0.267921568 |
| 202 | 52 | 0.795276 | 0.740359 | 0.843158 | 0.299576626 |
| 182 | 57 | 0.761506 | 0.702323 | 0.814078 | 0.878670935 |
| 176 | 42 | 0.807339 | 0.748637 | 0.857478 | 0.173610833 |
| 958 | 401 | 0.70493 | 0.67989 | 0.729081 | 0.929018242 |
| 1345 | 621 | 0.68413 | 0.663064 | 0.704646 | 0.064035998 |
| 1071 | 407 | 0.724628 | 0.701085 | 0.747275 | 0.077449107 |
| 2259 | 925 | 0.709485 | 0.693371 | 0.725213 | 0.534382324 |
| 1597 | 755 | 0.678997 | 0.659702 | 0.697844 | **0.007667226** |
| 1950 | 874 | 0.69051 | 0.673092 | 0.707532 | 0.112361412 |
| 1048 | 381 | 0.73338 | 0.709639 | 0.756158 | **0.016117162** |
| 871 | 341 | 0.718647 | 0.69241 | 0.743819 | 0.284626995 |
| 753 | 292 | 0.720574 | 0.692294 | 0.747606 | 0.263336317 |
| 667 | 719 | 0.481241 | 0.454639 | 0.507922 | **0.035333594** |
| 931 | 1186 | 0.439773 | 0.418489 | 0.461224 | 0.22986412 |
| 725 | 816 | 0.470474 | 0.445305 | 0.495755 | 0.17503583 |
| 1466 | 1614 | 0.475974 | 0.458204 | 0.49379 | **8.45E-06** |
| 863 | 1286 | 0.401582 | 0.380769 | 0.422664 | **0.001283058** |

| | | | | | |
|---|---|---|---|---|---|
| 1359 | 1658 | 0.450447 | 0.432583 | 0.468408 | 0.110264688 |
| 708 | 853 | 0.453555 | 0.428646 | 0.47864 | 0.168150031 |
| 614 | 692 | 0.470138 | 0.442779 | 0.497631 | **0.013020565** |
| 576 | 585 | 0.496124 | 0.466974 | 0.525294 | **3.85E-05** |
| 230 | 325 | 0.414414 | 0.37308 | 0.456665 | 0.248351378 |
| 419 | 466 | 0.473446 | 0.440121 | 0.506948 | **0.042109871** |
| 297 | 380 | 0.4387 | 0.40092 | 0.477017 | 1 |
| 623 | 716 | 0.465273 | 0.438284 | 0.492414 | 0.426116424 |
| 246 | 438 | 0.359649 | 0.323626 | 0.396889 | **6.77E-07** |
| 621 | 614 | 0.502834 | 0.474565 | 0.531089 | **0.00060275** |
| 267 | 337 | 0.442053 | 0.401986 | 0.482689 | 0.567393263 |
| 203 | 285 | 0.415984 | 0.371853 | 0.461139 | 0.092668718 |
| 214 | 246 | 0.465217 | 0.4189 | 0.511984 | 0.639851048 |
| 568 | 544 | 0.510791 | 0.480965 | 0.54056 | 0.741432849 |
| 834 | 860 | 0.492326 | 0.468254 | 0.516424 | 0.054777867 |
| 577 | 595 | 0.492321 | 0.463322 | 0.521358 | 0.108012706 |
| 556 | 571 | 0.493345 | 0.463762 | 0.522963 | 0.474666514 |
| 505 | 505 | 0.5 | 0.468707 | 0.531293 | 0.8012607 |
| 480 | 426 | 0.529801 | 0.496697 | 0.562711 | 0.126591525 |
| 572 | 1253 | 0.313425 | 0.292182 | 0.335269 | 0.194808099 |
| 758 | 1778 | 0.298896 | 0.281123 | 0.317135 | **0.00187042** |
| 585 | 1348 | 0.302638 | 0.282212 | 0.323665 | **0.018752003** |
| 1306 | 2908 | 0.309919 | 0.295973 | 0.324129 | **0.000312785** |
| 850 | 1765 | 0.325048 | 0.307108 | 0.34338 | 0.238062988 |
| 1210 | 2451 | 0.330511 | 0.315277 | 0.346016 | 0.484090038 |
| 562 | 1270 | 0.306769 | 0.285702 | 0.328456 | **0.008122304** |
| 552 | 1075 | 0.339275 | 0.316269 | 0.362862 | 0.793012507 |
| 437 | 1002 | 0.303683 | 0.279997 | 0.328173 | **0.009433606** |
| 112 | 167 | 0.401434 | 0.343443 | 0.461541 | 0.537909869 |
| 171 | 251 | 0.405213 | 0.357998 | 0.453768 | 0.341294075 |
| 97 | 160 | 0.377432 | 0.317944 | 0.439783 | 0.898017398 |
| 220 | 372 | 0.371622 | 0.332575 | 0.411956 | 0.121122711 |
| 171 | 205 | 0.454787 | 0.403656 | 0.506636 | **0.045739865** |
| 221 | 296 | 0.427466 | 0.384376 | 0.47139 | 0.282045511 |
| 103 | 157 | 0.396154 | 0.336269 | 0.458435 | 0.849685001 |
| 96 | 130 | 0.424779 | 0.359489 | 0.492069 | 0.541906257 |
| 64 | 129 | 0.331606 | 0.265654 | 0.402824 | **0.047335997** |
| 1109 | 1432 | 0.436442 | 0.417045 | 0.455986 | **0.034807661** |
| 1613 | 1895 | 0.459806 | 0.443211 | 0.476469 | 0.773314388 |
| 1256 | 1428 | 0.467958 | 0.448939 | 0.487048 | 0.277972496 |
| 2549 | 3076 | 0.453156 | 0.440085 | 0.466275 | 0.555977672 |
| 1848 | 1847 | 0.500135 | 0.483884 | 0.516387 | **1.78E-07** |
| 1994 | 2525 | 0.441248 | 0.426702 | 0.45587 | **0.032742983** |
| 1119 | 1352 | 0.452853 | 0.43309 | 0.472728 | 0.671601726 |
| 949 | 1214 | 0.438742 | 0.417694 | 0.459957 | 0.088197208 |
| 897 | 1090 | 0.451434 | 0.429387 | 0.473625 | 0.620363716 |
| 46 | 304 | 0.131429 | 0.097853 | 0.171391 | 0.455374422 |

| | | | | | |
|---|---|---|---|---|---|
| 58 | 448 | 0.114625 | 0.088199 | 0.145649 | 0.890356244 |
| 49 | 339 | 0.126289 | 0.094908 | 0.163504 | 0.582400485 |
| 120 | 716 | 0.143541 | 0.120465 | 0.169161 | **0.005014387** |
| 53 | 388 | 0.120181 | 0.091334 | 0.154239 | 0.545630577 |
| 90 | 823 | 0.098576 | 0.080013 | 0.119775 | 0.226707506 |
| 41 | 317 | 0.114525 | 0.08345 | 0.152147 | 0.866592002 |
| 33 | 268 | 0.109635 | 0.076679 | 0.150513 | 1 |
| 26 | 253 | 0.09319 | 0.06178 | 0.133561 | 0.39157731 |
| 378 | 333 | 0.531646 | 0.494197 | 0.56883 | 0.104048375 |
| 604 | 394 | 0.60521 | 0.574109 | 0.635689 | **0.006664487** |
| 437 | 342 | 0.560976 | 0.525309 | 0.596179 | 0.942450795 |
| 922 | 761 | 0.547831 | 0.52369 | 0.571806 | 0.109868205 |
| 598 | 463 | 0.563619 | 0.533165 | 0.59372 | 0.828326543 |
| 858 | 665 | 0.563362 | 0.538023 | 0.588456 | 0.776079423 |
| 435 | 333 | 0.566406 | 0.530508 | 0.601793 | 0.970962246 |
| 371 | 271 | 0.577882 | 0.538613 | 0.616431 | 0.604700908 |
| 298 | 227 | 0.567619 | 0.524009 | 0.610464 | 1 |
| 37 | 221 | 0.143411 | 0.103027 | 0.192215 | 1 |
| 44 | 286 | 0.133333 | 0.098579 | 0.174833 | 0.638285376 |
| 51 | 233 | 0.179577 | 0.136728 | 0.229256 | 0.091615892 |
| 82 | 490 | 0.143357 | 0.115667 | 0.174789 | **0.022181493** |
| 45 | 240 | 0.157895 | 0.117561 | 0.205512 | 0.35565156 |
| 61 | 306 | 0.166213 | 0.129589 | 0.208323 | 0.540858244 |
| 31 | 217 | 0.125 | 0.086535 | 0.172714 | **0.025214559** |
| 32 | 165 | 0.162437 | 0.113838 | 0.221504 | 0.578339895 |
| 36 | 167 | 0.17734 | 0.127406 | 0.23696 | 1 |
| 1060 | 534 | 0.664994 | 0.641222 | 0.688156 | 0.936627626 |
| 1654 | 811 | 0.670994 | 0.652048 | 0.689533 | 0.468548766 |
| 1174 | 590 | 0.665533 | 0.642971 | 0.687543 | 0.89973955 |
| 2601 | 1354 | 0.657649 | 0.642622 | 0.672442 | 0.060731108 |
| 1599 | 869 | 0.647893 | 0.62868 | 0.666754 | 0.643967358 |
| 2295 | 959 | 0.705286 | 0.689288 | 0.720914 | **7.66E-14** |
| 1133 | 578 | 0.662186 | 0.63922 | 0.684594 | 0.106310324 |
| 975 | 505 | 0.658784 | 0.633994 | 0.682941 | 0.222110552 |
| 879 | 459 | 0.656951 | 0.630817 | 0.682393 | 0.304395047 |
| 1611 | 1005 | 0.615826 | 0.596874 | 0.634517 | 0.888043397 |
| 2335 | 1511 | 0.607124 | 0.591483 | 0.622603 | 0.201513066 |
| 1688 | 1142 | 0.596466 | 0.578122 | 0.614611 | **0.023659545** |
| 3607 | 2365 | 0.603985 | 0.591449 | 0.61642 | 0.053707033 |
| 2205 | 1380 | 0.615063 | 0.598907 | 0.631031 | 0.890747003 |
| 3128 | 2069 | 0.601886 | 0.588427 | 0.61523 | **0.034786286** |
| 1720 | 1036 | 0.624093 | 0.605704 | 0.642218 | 0.399743388 |
| 1469 | 913 | 0.616709 | 0.59684 | 0.63629 | 0.966398365 |
| 1242 | 773 | 0.616377 | 0.59474 | 0.637675 | 1 |
| 1030 | 1780 | 0.366548 | 0.348701 | 0.384674 | 0.443414163 |
| 1535 | 2727 | 0.36016 | 0.34573 | 0.374781 | 0.949099704 |
| 1125 | 1937 | 0.367407 | 0.350301 | 0.384766 | 0.376177055 |

| | | | | | |
|---|---|---|---|---|---|
| 2347 | 4465 | 0.344539 | 0.333249 | 0.355962 | 0.22806163 |
| 1893 | 3030 | 0.384522 | 0.370903 | 0.398278 | **1.53E-06** |
| 2114 | 3823 | 0.356072 | 0.343881 | 0.368405 | 0.471334798 |
| 1043 | 1774 | 0.370252 | 0.352386 | 0.388388 | **0.038300673** |
| 903 | 1650 | 0.353702 | 0.335136 | 0.372603 | 0.81969128 |
| 887 | 1386 | 0.390233 | 0.370112 | 0.410638 | **0.000131378** |
| 437 | 491 | 0.470905 | 0.438387 | 0.503608 | 0.2786731 |
| 637 | 649 | 0.495334 | 0.467657 | 0.523033 | 0.655428253 |
| 469 | 484 | 0.49213 | 0.459931 | 0.524378 | 0.845917153 |
| 911 | 1174 | 0.43693 | 0.415502 | 0.458537 | **2.57E-08** |
| 608 | 607 | 0.500412 | 0.471914 | 0.528908 | 0.863384842 |
| 983 | 831 | 0.541896 | 0.51864 | 0.565017 | **0.000187507** |
| 457 | 491 | 0.482068 | 0.44983 | 0.514416 | 0.330127998 |
| 392 | 436 | 0.47343 | 0.438964 | 0.508086 | 0.164515965 |
| 349 | 404 | 0.463479 | 0.427401 | 0.499845 | 0.063022728 |

| Suppl Table 2 | | | | | Alleles | | Frequency alt ge |
|---|---|---|---|---|---|---|---|
| Fragment number | | | hg19 position | | ref | alt | EUR/JPT |
| 1 | | | 45378144 | **rs34278513** | C | T | 0.106 / 0.154 |
| 1 | | | 45378357 | rs73556120 | C | T | 0 / 0 |
| 1 | | | 45378540 | rs57907894 | C | CT | 0.182 / 0.476 |
| 2 | | | 45380545 | **rs421812** | G | T | 0.273 / 0.740 |
| 2 | | | 45380961 | **rs3865427** | C | A | 0.106 / 0.159 |
| 2 | | | 45380970 | rs11668861 | G | T | 0.460 / 0.755 |
| 2 | | | 45381292 | rs150639620 | G | T | 0.066 / 0 |
| 2 | | | 45381314 | rs143981846 | CCTCCTCCT | C | 0.061 / 0 |
| 3 | | | 45382675 | rs41290120 | G | A | 0.035 /0 |
| 3 | | | 45382717 | rs406456 | G | A | 0.576 / 0.909 |
| 3 | | | 45382966 | **rs3852860** | C | T | 0.404 / 0.760 |
| 3 | | | 45382984 | rs11669338 | T | G | 0.091 / 0 |
| 3 | | | 45383037 | rs11673139 | A | T | 0.091 / 0 |
| 3 | | | 45383061 | **rs3852861** | G | T | 0.409 / 0.760 |
| 3 | | | 45383079 | **rs71352237** | T | C | 0.106 / 0.159 |
| 3 | | | 45383095 | **rs71171301** | A | AC | 0.111 / 0.159 |
| 3 | | | 45383115 | **rs34224078** | A | G | 0.111 / 0.159 |
| 3 | | | 45383139 | **rs35879138** | T | A | 0.111 / 0.159 |
| 4 | | | 45386460 | rs76725281 | A | G | 0.010 / 0.010 |
| 4 | | | 45386470 | **rs142042446** | G | GTAA | 0.182 / 0.135 |
| 4 | | | 45386634 | rs147636938 | C | T | 0.035 / 0 |
| 4 | 5 | | 45386855 | **rs166907** | A | G | 0 / 0.014 |
| 5 | | | 45387034 | **rs283808** | A | C | 0.030 / 0.072 |
| 5 | | | 45387057 | **rs283809** | A | G | 0.030 / 0.072 |
| 6 | | | 45387459 | **rs12972156** | C | G | 0.187 / 0.115 |
| 6 | | | 45387596 | **rs12972970** | G | A | 0.187 / 0.115 |
| 7 | | | 45388130 | **rs34342646** | G | A | 0.192 / 0.115 |
| 7 | | | 45388241 | rs283810 | T | G | 0.066 / 0.072 |
| 8 | | | 45392254 | **rs6857** | C | T | 0.197 / 0.120 |
| 9 | | | 45394336 | **rs71352238** | T | C | 0.172 / 0.120 |
| 10 | | | 45395266 | **rs157580** | G | A | 0.626 / 0.457 |
| 10 | | | 45395330 | rs2075649 | A | G | 0.389 / 0.260 |
| 10 | 11 | | 45395619 | **rs2075650** | A | G | 0.167 / 0.120 |
| 10 | 11 | | 45395714 | **rs157581** | T | C | 0.237 / 0.260 |
| 10 | 11 | 12 | 45395816 | rs73936968 | G | A | 0.020 / 0 |
| 10 | 11 | 12 | 45395844 | **rs34095326** | G | A | 0.126 / 0 |
| 10 | 11 | 12 | 45395909 | **rs34404554** | C | G | 0.167 / 0.120 |
| 11 | | 12 | 45396144 | **rs11556505** | C | T | 0.167 / 0.120 |
| 11 | | 12 | 45396219 | rs157582 | C | T | 0.237 / 0.202 |
| 13 | 14 | | 45396665 | **rs59007384** | G | T | 0.232 / 0.183 |
| 13 | 14 | | 45396673 | **rs157583** | G | T | 0 / 0.014 |

| | | | | | | |
|---|---|---|---|---|---|---|
| 13 | 14 | 45396899 | **rs157584** | T | C | 0.455 / 0.293 |
| 13 | 14 | 45396973 | rs77301115 | G | A | 0.030 / 0 |
| 14 | 15 | 45397171 | rs73936970 | G | C | 0.010 / 0.014 |
| 14 | 15 | 45397229 | rs1160983 | G | A | 0.025 / 0.053 |
| 14 | 15 | 45397307 | rs112849259 | C | T | 0.030 / 0 |
| 15 | | 45397512 | **rs157585** | A | C | 0.450 / 0.303 |
| 16 | | 45397952 | rs116881820 | T | C | 0.030 / 0 |
| 16 | | 45397965 | rs115676124 | G | A | 0 / 0 |
| 16 | | 45398206 | **rs157587** | A | G | 0 / 0.014 |
| 16 | | 45398264 | **rs157588** | C | T | 0.450 / 0.293 |
| 16 | 17 | 45398457 | rs77726367 | G | A | 0 / 0 |
| 16 | 17 | 45398633 | rs11668327 | G | C | 0.202 / 0 |
| 17 | | 45398716 | **rs157590** | A | C | 0.465 / 0.293 |
| 17 | | 45398785 | rs79398853 | C | T | 0.030 / 0 |
| 17 | | 45398817 | **rs2238681** | C | T | 0.384 / 0.202 |
| 18 | | 45400491 | rs150006531 | T | TG | 0.030 / 0 |
| 18 | | 45400725 | rs114536010 | C | T | 0.030 / 0 |
| 18 | | 45400747 | rs61679753 | T | A | 0.025 / 0.053 |
| 18 | | 45400775 | **rs205909** | T | G | 0 / 0.014 |
| 18 | | 45400871 | rs116874600 | G | A | 0.010 / 0.014 |
| 20 | | 45402718 | rs35568738 | G | C | 0.046 / 0 |
| 20 | | 45403119 | **rs417357** | C | T | 0 / 0.014 |
| 20 | | 45403216 | rs115881343 | C | T | 0.030 / 0 |
| 20 | | 45403412 | rs1160985 | C | T | 0.419 / 0.284 |
| 20 | | 45403458 | rs77100236 | C | T | 0.005 / 0 |
| 21 | | 45404431 | rs741780 | T | C | 0.419 / 0.284 |
| 21 | | 45404432 | rs117264457 | G | A | 0.025 / 0 |
| 21 | | 45404579 | **rs394819** | G | T | 0 / 0.014 |
| 21 | | 45404691 | rs405697 | A | G | 0.763 / 0.385 |
| 21 | | 45404721 | rs116977783 | C | T | 0.010 / 0.014 |
| 21 | | 45404830 | rs113492558 | TGAAACTCCG | A | 0.030 / 0 |
| 21 | | 45404857 | rs112019714 | T | C | 0.030 / 0 |
| 22 | | 45407118 | **rs435380** | G | A | 0 / 0.014 |
| 22 | | 45407226 | rs66916483 | C | A | 0 / 0 |
| 22 | | 45407437 | **rs446037** | G | T | 0 / 0.014 |
| 22 | | 45407655 | rs7256173 | C | T | 0.010 / 0.014 |
| 22 | | 45407720 | **rs434132** | C | G | 0 / 0.014 |
| 22 | | 45407788 | rs7259620 | G | A | 0.419 / 0.284 |
| 23 | | 45408475 | **rs439382** | A | G | 0 / 0.014 |
| 23 | | 45408564 | **rs449647** | A | T | 0.202 / 0.034 |
| 23 | | 45408628 | rs769446 | T | C | 0.086 / 0.048 |
| 23 | | 45408836 | rs405509 | T | G | 0.490 / 0.269 |
| 24 | | 45410002 | **rs769449** | G | A | 0.146 / 0.067 |

| | | | | | | |
|---|---|---|---|---|---|---|
| 24 | | 45410444 | rs769450 | G | A | 0.384 / 0.212 |
| 25 | | 45413366 | rs1081106 | T | C | 0.106 / 0 |
| 25 | | 45413576 | **rs75627662** | C | T | 0.207 / 0.125 |
| 25 | | 45413788 | **rs148353395** | TCCCCGGCCT( | C | 0 / 0.014 |
| 25 | | 45414109 | rs72654471 | T | C | 0 / 0 |
| 26 | | 45415491 | rs117650633 | T | C | 0 / 0.115 |
| 26 | | 45415640 | **rs445925** | G | A | 0.096 / 0.058 |
| 26 | | 45415713 | **rs10414043** | G | A | 0.141 / 0.077 |
| 26 | | 45415935 | **rs7256200** | G | T | 0.141 / 0.077 |
| 26 | | 45416099 | rs72654437 | G | A | 0.035 / 0 |
| 26 | | 45416178 | rs483082 | G | T | 0.237 / 0.135 |
| 27 | | 45416478 | rs584007 | A | G | 0.636 / 0.370 |
| 27 | | 45416741 | rs438811 | C | T | 0.237 / 0.135 |
| 27 | | 45416831 | **rs390082** | T | G | 0.096 / 0.058 |
| 28 | | 45417640 | **rs34954997** | C | CTTCG | 0.237 / 0.120 |
| 28 | | 45417744 | rs184802815 | G | A | 0 / 0 |
| 31 | | 45421100 | rs3925681 | G | A | 0.404 / 0.226 |
| 31 | | 45421204 | rs150966173 | C | T | 0.030 / 0 |
| 31 | | 45421254 | **rs12721046** | G | A | 0.187 / 0.091 |
| 32 | | 45421877 | rs484195 | A | G | 0.616 / 0.385 |
| 32 | | 45421926 | rs112528434 | G | T | 0 / 0 |
| 32 | | 45421973 | rs12721052 | AG | A | 0.338 / 0.231 |
| 32 | | 45422160 | **rs12721051** | C | G | 0.217 / 0.091 |
| 32 | | 45422561 | rs1064725 | T | G | 0.076 / 0 |
| 32 | | 45422587 | rs12721054 | A | G | 0 /0 |
| 32 | 33 | 45422561 | rs1064725 | T | G | 0.076 / 0 |
| 32 | 33 | 45422587 | rs12721054 | A | G | 0 / 0 |
| 33 | | 45422846 | **rs56131196** | G | A | 0.217 / 0.091 |
| 33 | | 45422946 | **rs4420638** | A | G | 0.217 / 0.091 |
| 34 | | 45423636 | rs78959900 | G | A | 0.343 / 0.231 |
| 34 | | 45423934 | **rs157591** | G | A | 0 / 0 |
| 35 | | 45426105 | **rs157596** | C | T | 0 / 0 |
| 36 | | 45426637 | **rs157597** | A | C | 0 / 0 |
| 36 | 37 | 45426792 | rs141622900 | G | A | 0.046 / 0.053 |
| 36 | 37 | 45426813 | **rs157598** | C | T | 0 / 0 |
| 37 | | 45427074 | rs138682750 | A | T | 0 /0 |
| 37 | | 45427125 | **rs111789331** | T | A | 0.187 / 0.096 |
| 37 | | 45427162 | rs113086796 | G | A | 0 / 0 |
| 37 | | 45427213 | rs112490204 | G | T | 0 / 0 |
| 37 | | 45427353 | rs4803770 | C | G | 0.364 / 0.216 |
| 38 | | 45427931 | **rs157599** | A | G | 0 / 0 |
| 38 | | 45428083 | **rs149345** | T | G | 0 / 0 |
| 38 | | 45428234 | **rs66626994** | G | A | 0.187 / 0.096 |

| | | | | | |
|---|---|---|---|---|---|
| 38 | 45428459 | rs4803772 | C | T | 0.343 / 0.221 |
| 39 | 45429099 | **rs10424663** | G | A | 0 / 0.019 |
| 39 | 45429177 | rs147378144 | T | C | 0 / 0 |
| 39 | 45429543 | rs71352239 | C | T | 0.318 / 0.236 |

Green highlight; sTV

| neral population | Frequency alt on ε4 | | | Differences on ε4 | | |
|---|---|---|---|---|---|---|
| AFR | EUR | JPT | AFR | EURvsAFR | p-value | JPTvsAFR |
| 0 | 0.229 | 0.706 | 0.000 | -0.22857143 | 1 | -0.70588235 |
| 0 | 0.000 | 0.000 | 0.000 | 0 | 1 | 0 |
| 0.213 | 0.086 | 0.059 | 0.102 | 0.01632653 | 1 | 0.04321729 |
| 0.417 | 0.314 | 0.941 | 0.327 | 0.0122449 | 1 | -0.61464586 |
| 0 | 0.229 | 0.706 | 0.000 | -0.22857143 | 1 | -0.70588235 |
| 0.482 | 0.171 | 0.118 | 0.449 | 0.27755102 | 1 | 0.33133253 |
| 0.032 | 0.000 | 0.000 | 0.082 | 0.08163265 | 1 | 0.08163265 |
| 0.046 | 0.000 | 0.000 | 0.163 | 0.16326531 | 1 | 0.16326531 |
| 0 | 0.000 | 0.000 | 0.000 | 0 | 1 | 0 |
| 0.759 | 0.343 | 1.000 | 0.673 | 0.33061224 | 1 | -0.32653061 |
| 0.329 | 0.143 | 0.118 | 0.184 | 0.04081633 | 1 | 0.06602641 |
| 0.111 | 0.029 | 0.000 | 0.306 | 0.27755102 | 1 | 0.30612245 |
| 0.111 | 0.029 | 0.000 | 0.306 | 0.27755102 | 1 | 0.30612245 |
| 0.329 | 0.143 | 0.118 | 0.184 | 0.04081633 | 1 | 0.06602641 |
| 0.111 | 0.229 | 0.706 | 0.000 | -0.22857143 | 1 | -0.70588235 |
| 0.125 | 0.229 | 0.706 | 0.020 | -0.20816327 | 1 | -0.68547419 |
| 0.125 | 0.229 | 0.706 | 0.020 | -0.20816327 | 1 | -0.68547419 |
| 0.125 | 0.229 | 0.706 | 0.020 | -0.20816327 | 1 | -0.68547419 |
| 0 | 0.000 | 0.000 | 0.000 | 0 | 1 | 0 |
| 0 | 0.771 | 0.824 | 0.000 | -0.77142857 | 3.0185E-08 | -0.82352941 |
| 0.032 | 0.171 | 0.000 | 0.020 | -0.15102041 | 1 | 0.02040816 |
| 0.153 | 0.000 | 0.176 | 0.388 | 0.3877551 | 1 | 0.21128451 |
| 0.361 | 0.000 | 0.176 | 0.776 | 0.7755102 | 5.6316E-07 | 0.59903962 |
| 0.361 | 0.000 | 0.176 | 0.776 | 0.7755102 | 5.6316E-07 | 0.59903962 |
| 0 | 0.771 | 0.824 | 0.000 | -0.77142857 | 3.0185E-08 | -0.82352941 |
| 0 | 0.771 | 0.824 | 0.000 | -0.77142857 | 3.0185E-08 | -0.82352941 |
| 0 | 0.771 | 0.824 | 0.000 | -0.77142857 | 3.0185E-08 | -0.82352941 |
| 0.370 | 0.171 | 0.176 | 0.653 | 0.48163265 | 1 | 0.47659064 |
| 0.037 | 0.943 | 0.824 | 0.020 | -0.92244898 | 8.4354E-12 | -0.80312125 |
| 0 | 0.771 | 0.824 | 0.000 | -0.77142857 | 3.0185E-08 | -0.82352941 |
| 0.889 | 0.971 | 1.000 | 0.980 | 0.00816327 | 1 | -0.02040816 |
| 0.107 | 0.029 | 0.000 | 0.061 | 0.03265306 | 1 | 0.06122449 |
| 0.139 | 0.771 | 0.824 | 0.122 | -0.64897959 | 0.00046176 | -0.70108043 |
| 0.495 | 0.943 | 1.000 | 0.918 | -0.0244898 | 1 | -0.08163265 |
| 0.107 | 0.000 | 0.000 | 0.061 | 0.06122449 | 1 | 0.06122449 |
| 0 | 0.543 | 0.000 | 0.000 | -0.54285714 | 0.00131796 | 0 |
| 0.097 | 0.771 | 0.824 | 0.122 | -0.64897959 | 0.00046176 | -0.70108043 |
| 0.125 | 0.771 | 0.824 | 0.122 | -0.64897959 | 0.00046176 | -0.70108043 |
| 0.514 | 0.943 | 1.000 | 0.918 | -0.0244898 | 1 | -0.08163265 |
| 0.319 | 0.943 | 0.824 | 0.224 | -0.71836735 | 2.0957E-05 | -0.59903962 |
| 0.157 | 0.000 | 0.176 | 0.612 | 0.6122449 | 0.00181752 | 0.43577431 |

| | | | | | | |
|---|---|---|---|---|---|---|
| 0.875 | 0.200 | 0.176 | 1.000 | 0.8 | 6.4548E-09 | 0.82352941 |
| 0.051 | 0.171 | 0.000 | 0.184 | 0.0122449 | 1 | 0.18367347 |
| 0.065 | 0.000 | 0.000 | 0.020 | 0.02040816 | 1 | 0.02040816 |
| 0.097 | 0.000 | 0.000 | 0.041 | 0.04081633 | 1 | 0.04081633 |
| 0.046 | 0.171 | 0.000 | 0.184 | 0.0122449 | 1 | 0.18367347 |
| 0.894 | 0.171 | 0.176 | 1.000 | 0.82857143 | 1.3069E-09 | 0.82352941 |
| 0.046 | 0.171 | 0.000 | 0.184 | 0.0122449 | 1 | 0.18367347 |
| 0.065 | 0.000 | 0.000 | 0.000 | 0 | 1 | 0 |
| 0.093 | 0.000 | 0.176 | 0.367 | 0.36734694 | 1 | 0.19087635 |
| 0.894 | 0.171 | 0.176 | 1.000 | 0.82857143 | 1.3069E-09 | 0.82352941 |
| 0.065 | 0.000 | 0.000 | 0.000 | 0 | 1 | 0 |
| 0 | 0.029 | 0.000 | 0.000 | -0.02857143 | 1 | 0 |
| 0.898 | 0.171 | 0.176 | 1.000 | 0.82857143 | 1.3069E-09 | 0.82352941 |
| 0.046 | 0.171 | 0.000 | 0.184 | 0.0122449 | 1 | 0.18367347 |
| 0.222 | 0.000 | 0.000 | 0.000 | 0 | 1 | 0 |
| 0.056 | 0.171 | 0.000 | 0.184 | 0.0122449 | 1 | 0.18367347 |
| 0.046 | 0.171 | 0.000 | 0.184 | 0.0122449 | 1 | 0.18367347 |
| 0.116 | 0.000 | 0.000 | 0.041 | 0.04081633 | 1 | 0.04081633 |
| 0.139 | 0.000 | 0.176 | 0.551 | 0.55102041 | 0.02135444 | 0.37454982 |
| 0 | 0.000 | 0.000 | 0.000 | 0 | 1 | 0 |
| 0 | 0.000 | 0.000 | 0.000 | 0 | 1 | 0 |
| 0.134 | 0.000 | 0.176 | 0.551 | 0.55102041 | 0.02135444 | 0.37454982 |
| 0.046 | 0.171 | 0.000 | 0.184 | 0.0122449 | 1 | 0.18367347 |
| 0.662 | 0.000 | 0.000 | 0.102 | 0.10204082 | 1 | 0.10204082 |
| 0 | 0.000 | 0.000 | 0.000 | 0 | 1 | 0 |
| 0.662 | 0.000 | 0.000 | 0.102 | 0.10204082 | 1 | 0.10204082 |
| 0 | 0.000 | 0.000 | 0.000 | 0 | 1 | 0 |
| 0.13 | 0.000 | 0.176 | 0.531 | 0.53061224 | 0.04599548 | 0.35414166 |
| 0.884 | 0.971 | 1.000 | 1.000 | 0.02857143 | 1 | 0 |
| 0 | 0.000 | 0.000 | 0.000 | 0 | 1 | 0 |
| 0.065 | 0.171 | 0.000 | 0.245 | 0.07346939 | 1 | 0.24489796 |
| 0.065 | 0.171 | 0.000 | 0.245 | 0.07346939 | 1 | 0.24489796 |
| 0.130 | 0.000 | 0.176 | 0.531 | 0.53061224 | 0.04599548 | 0.35414166 |
| 0.120 | 0.000 | 0.000 | 0.000 | 0 | 1 | 0 |
| 0.130 | 0.000 | 0.176 | 0.531 | 0.53061224 | 0.04599548 | 0.35414166 |
| 0.023 | 0.000 | 0.000 | 0.000 | 0 | 1 | 0 |
| 0.167 | 0.000 | 0.176 | 0.694 | 0.69387755 | 4.3786E-05 | 0.51740696 |
| 0.449 | 0.000 | 0.000 | 0.102 | 0.10204082 | 1 | 0.10204082 |
| 0.107 | 0.000 | 0.176 | 0.429 | 0.42857143 | 1 | 0.25210084 |
| 0.333 | 0.000 | 0.176 | 0.612 | 0.6122449 | 0.00181752 | 0.43577431 |
| 0.018 | 0.114 | 0.000 | 0.000 | -0.11428571 | 1 | 0 |
| 0.769 | 0.171 | 0.000 | 0.469 | 0.29795918 | 1 | 0.46938776 |
| 0 | 0.829 | 0.824 | 0.000 | -0.82857143 | 1.3069E-09 | -0.82352941 |

| | | | | | | |
|---|---|---|---|---|---|---|
| 0.375 | 0.000 | 0.000 | 0.000 | 0 | 1 | 0 |
| 0 | 0.000 | 0.000 | 0.000 | 0 | 1 | 0 |
| 0.148 | 0.800 | 0.824 | 0.143 | -0.65714286 | 0.00040833 | -0.68067227 |
| 0.148 | 0.000 | 0.176 | 0.612 | 0.6122449 | 0.00181752 | 0.43577431 |
| 0.162 | 0.000 | 0.000 | 0.061 | 0.06122449 | 1 | 0.06122449 |
| 0 | 0.000 | 0.000 | 0.000 | 0 | 1 | 0 |
| 0.282 | 0.171 | 0.176 | 0.755 | 0.58367347 | 0.02627691 | 0.57863145 |
| 0.190 | 0.800 | 0.824 | 0.163 | -0.63673469 | 0.0013832 | -0.66026411 |
| 0.19 | 0.800 | 0.824 | 0.163 | -0.63673469 | 0.0013832 | -0.66026411 |
| 0 | 0.000 | 0.000 | 0.000 | 0 | 1 | 0 |
| 0.44 | 0.971 | 1.000 | 0.776 | -0.19591837 | 1 | -0.2244898 |
| 0.889 | 1.000 | 1.000 | 1.000 | 0 | 1 | 0 |
| 0.482 | 0.971 | 1.000 | 0.939 | -0.03265306 | 1 | -0.06122449 |
| 0.292 | 0.171 | 0.176 | 0.776 | 0.60408163 | 0.00959686 | 0.59903962 |
| 0.264 | 0.971 | 0.824 | 0.388 | -0.58367347 | 0.01000056 | -0.43577431 |
| 0 | 0.000 | 0.000 | 0.000 | 0 | 1 | 0 |
| 0.273 | 0.029 | 0.000 | 0.041 | 0.0122449 | 1 | 0.04081633 |
| 0.046 | 0.171 | 0.000 | 0.204 | 0.03265306 | 1 | 0.20408163 |
| 0 | 0.800 | 1.000 | 0.000 | -0.8 | 6.4548E-09 | -1 |
| 0.894 | 1.000 | 1.000 | 1.000 | 0 | 1 | 0 |
| 0.056 | 0.000 | 0.000 | 0.061 | 0.06122449 | 1 | 0.06122449 |
| 0.245 | 0.029 | 0.000 | 0.041 | 0.0122449 | 1 | 0.04081633 |
| 0.060 | 0.971 | 1.000 | 0.224 | -0.74693878 | 3.6165E-06 | -0.7755102 |
| 0 | 0.000 | 0.000 | 0.000 | 0 | 1 | 0 |
| 0.111 | 0.000 | 0.000 | 0.102 | 0.10204082 | 1 | 0.10204082 |
| 0 | 0.000 | 0.000 | 0.000 | 0 | 1 | 0 |
| 0.111 | 0.000 | 0.000 | 0.102 | 0.10204082 | 1 | 0.10204082 |
| 0.153 | 0.971 | 1.000 | 0.327 | -0.64489796 | 0.00068417 | -0.67346939 |
| 0.153 | 0.971 | 1.000 | 0.327 | -0.64489796 | 0.00068417 | -0.67346939 |
| 0.319 | 0.029 | 0.000 | 0.041 | 0.0122449 | 1 | 0.04081633 |
| 0.130 | 0.000 | 0.000 | 0.531 | 0.53061224 | 0.04599548 | 0.53061224 |
| 0.319 | 0.000 | 0.000 | 0.551 | 0.55102041 | 0.02135444 | 0.55102041 |
| 0.319 | 0.000 | 0.000 | 0.551 | 0.55102041 | 0.02135444 | 0.55102041 |
| 0.083 | 0.000 | 0.000 | 0.000 | 0 | 1 | 0 |
| 0.116 | 0.000 | 0.000 | 0.469 | 0.46938776 | 0.39865202 | 0.46938776 |
| 0.032 | 0.000 | 0.000 | 0.041 | 0.04081633 | 1 | 0.04081633 |
| 0 | 0.800 | 0.941 | 0.000 | -0.8 | 6.4548E-09 | -0.94117647 |
| 0.106 | 0.000 | 0.000 | 0.000 | 0 | 1 | 0 |
| 0.051 | 0.000 | 0.000 | 0.102 | 0.10204082 | 1 | 0.10204082 |
| 0.282 | 0.029 | 0.000 | 0.102 | 0.07346939 | 1 | 0.10204082 |
| 0.361 | 0.000 | 0.000 | 0.551 | 0.55102041 | 0.02135444 | 0.55102041 |
| 0.329 | 0.000 | 0.000 | 0.551 | 0.55102041 | 0.02135444 | 0.55102041 |
| 0.093 | 0.800 | 0.941 | 0.020 | -0.77959184 | 4.3691E-08 | -0.92076831 |

| | | | | | | |
|---|---|---|---|---|---|---|
| 0.241 | 0.029 | 0.000 | 0.041 | 0.0122449 | 1 | 0.04081633 |
| 0.134 | 0.000 | 0.059 | 0.531 | 0.53061224 | 0.04599548 | 0.47178872 |
| 0.056 | 0.000 | 0.000 | 0.041 | 0.04081633 | 1 | 0.04081633 |
| 0.083 | 0.086 | 0.118 | 0.041 | -0.04489796 | 1 | -0.07683073 |

| | PCR-based MPRA fragments | | Luciferase | |
|---|---|---|---|---|
| p-value | AFR | EUR/JPT | AFR | EUR/JPT |
| 5.1282E-05 | C | T | | |
| 1 | C | C | | |
| 1 | - | - | | |
| 1 | T | G | T | G |
| 5.1282E-05 | C | A | C | A |
| 1 | G | G | G | G |
| 1 | G | G | G | G |
| 1 | CTCCTCCTC | CTCCTCCTC | CTCCTCCTC | CTCCTCCTC |
| 1 | G | G | A | G |
| 1 | A | A | A | A |
| 1 | T | C | T | C |
| 1 | T | T | T | T |
| 1 | A | A | A | A |
| 1 | T | G | T | G |
| 5.1282E-05 | T | C | T | C |
| 0.00048541 | - | C | - | C |
| 0.00048541 | A | G | A | G |
| 0.00048541 | T | A | T | A |
| 1 | A | A | | |
| 5.8546E-07 | - | TAA | | |
| 1 | C | C | | |
| 1 | G | A | G | A |
| 1 | C | A | C | A |
| 1 | G | A | G | A |
| 5.8546E-07 | C | G | | |
| 5.8546E-07 | G | A | | |
| 5.8546E-07 | G | A | G | A |
| 1 | G | G | G | G |
| 5.8754E-06 | C | T | C | T |
| 5.8546E-07 | T | C | | |
| 1 | A | G | | |
| 1 | A | A | | |
| 0.01929632 | A | G | A | G |
| 1 | T | C | C | C |
| 1 | G | G | G | G |
| 1 | G | 10-G 11/12-A | 11-G 12-A | A |
| 0.01929632 | C | G | 11-C 12-G | G |
| 0.01929632 | C | T | C | T |
| 1 | T | T | T | T |
| 1 | G | T | G | T |
| 1 | T | G | G | G |

| | | | | |
|---|---|---|---|---|
| 5.8546E-07 | **C** | **T** | **C** | **C** |
| 1 | G | G | G | G |
| 1 | G | G | G | G |
| 1 | G | G | G | G |
| 1 | C | C | C | C |
| 5.8546E-07 | **C** | **A** | **C** | **A** |
| 1 | T | T | T | T |
| 1 | G | G | G | G |
| 1 | G | A | G | A |
| 5.8546E-07 | **T** | **C** | **T** | **C** |
| 1 | G | G | G | G |
| 1 | G | G | G | G |
| 5.8546E-07 | **C** | **A** | **C** | **A** |
| 1 | C | C | C | C |
| 1 | T | C | T | C |
| 1 | - | - | | |
| 1 | C | C | | |
| 1 | T | T | | |
| 1 | **G** | **T** | | |
| 1 | G | G | | |
| 1 | G | G | | |
| 1 | **T** | **C** | | |
| 1 | C | C | | |
| 1 | C | C | | |
| 1 | C | C | | |
| 1 | T | T | | |
| 1 | G | G | | |
| 1 | **T** | **G** | | |
| 1 | G | G | | |
| 1 | C | C | | |
| 1 | CGGTGAAACTCCGTCTC | GGTGAAACTCCGTCTC | | |
| 1 | T | T | | |
| 1 | **A** | **G** | | |
| 1 | C | C | | |
| 1 | **T** | **G** | | |
| 1 | C | C | | |
| 1 | **G** | **C** | | |
| 1 | G | G | | |
| 1 | G | A | G | A |
| 1 | **T** | **A** | **T** | **A** |
| 1 | T | T | T | T |
| 1 | T | T | T | T |
| 5.8546E-07 | **G** | **A** | | |

| | | | | |
|---|---|---|---|---|
| 1 | G | G | | |
| 1 | A | A | A | A |
| 0.06160456 | **C** | **T** | **C** | **T** |
| 1 | **-** | **TTCTCCCCGGCCTGCTT** | **-** | **TCTCCCCGGCCTGCT** |
| 1 | T | T | T | T |
| 1 | C | C | C | C |
| 1 | **A** | **G** | **A** | **G** |
| 0.1771741 | **G** | **A** | **G** | **A** |
| 0.1771741 | **G** | **T** | **G** | **T** |
| 1 | G | G | G | G |
| 1 | T | T | T | T |
| 1 | G | G | | |
| 1 | T | T | | |
| 1 | **G** | **T** | | |
| 1 | **-** | **TTCG** | **-** | **TTCG** |
| 1 | G | G | G | G |
| 1 | G | G | | |
| 1 | C | C | | |
| 3.7096E-10 | **G** | **A** | | |
| 1 | A | G | | |
| 1 | G | G | | |
| 1 | G | G | | |
| 0.00746646 | **C** | **G** | | |
| 1 | T | T | | |
| 1 | A | A | | |
| 1 | T | T | T | T |
| 1 | A | A | A | A |
| 0.40719394 | **G** | **A** | **G** | **A** |
| 0.40719394 | **A** | **G** | **A** | **G** |
| 1 | ? | ? | | |
| 1 | **A** | **G** | | |
| 1 | **T** | **C** | | |
| 1 | **C** | **A** | **C** | **A** |
| 1 | G | G | G | G |
| 1 | T | C | T | C |
| 1 | A-stretch | A-stretch | A-stretch | A-stretch |
| 4.7488E-09 | **T** | **A** | **T** | **A** |
| 1 | G | G | G | G |
| 1 | G | G | G | G |
| 1 | C | C | C | C |
| 1 | **G** | **A** | **G** | **A** |
| 1 | **G** | **T** | **G** | **T** |
| 4.9728E-08 | **G** | **A** | **G** | **A** |

| | | | | |
|---|---|---|---|---|
| 1 | C | C | C | C |
| 1 | **A** | **G** | | |
| 1 | T | T | | |
| 1 | C | C | | |

**Supplementary Table 3. Primer sequences of PCR-based MPRA design**

| Purpose | Primer sequence |
|---|---|
| All APOE fragments | Gibson assembly overhang F; 5-CCGGTACCTGAGCTCG-3 |
| All APOE fragments | Gibson assembly overhang R; 5-CCGAGGCCAGATCTTGAT-3 |
| 1 | F; 5-GGTCATCTTTGTCCGAGGTG-3<br>R; 5-GATTGCGCCACTACACTTCA-3 |
| **2** | F; 5-GAGGTGTAGAGGTGGGGTCA-3<br>R; 5-GGAGACACAAGATCACGAGGA-3 |
| **3** | F; 5-TGGAGCCTCAGACCTCAACT-3<br>R; 5-TGAGGTTCGGCACACAGTAG-3 |
| **4** | F; 5-CACCGAGGCTGGAATACAGT-3<br>R; 5-CGAACCACTAGCAACAAGAGC-3 |
| **5** | F; 5-CCTCTCGAGTAGCTGGGATT-3<br>R; 5-GCATAGCTCGCATACCTCAG-3 |
| **6** | F; 5-TGGCGAAACCTGTCTACAAA-3<br>R; 5-CTCAGATGTCCCCTCTTCCA-3 |
| **7** | F; 5-GCCTGGAATCTGTATTTACCACA-3<br>R; 5-TGAGGTCAGGAGTGTGAGAC-3 |
| **8** | F; 5-ATGACCCCTAACCCATGAGC-3<br>R; 5-CTCTCTTGACCCCAGTTCACT-3 |
| **9** | F; 5-TTAGAATTCTCCCGAGGCAC-3<br>R; 5-TGCTAGGGGAGAGGTCAGAG-3 |
| **10** | F; 5-CTTGTGATCGGGAAGGCAAC-3<br>R; 5-CAGGGGATGAAGACTGGGAG-3 |
| **11** | F; 5-GGTGGCAGTGAGATTTGAGC-3<br>R; 5-ATGAGGCCGTCTTAGCTTGA-3 |
| **12** | F; 5-ACTCGGCCCAATTACTCCTC-3<br>R; 5-CCTCCAGTTTTGCCCTTCCT-3 |
| **13** | F; 5-CTACCGAGAGCCTGGAGATG-3<br>R; 5-ATGACCTGAGCGTTGAGACT-3 |
| **14** | F; 5-TGGAGAGTCATGGCATGGTT-3<br>R; 5-AGTTTTGTGACCACCCTCCA-3 |
| **15** | F; 5-AGTCTCAACGCTCAGGTCAT-3<br>R; 5-AGGTCAAGGTGGGAGGATTG-3 |
| **16** | F; 5-AGGCTGGTCTTGGAACTCC-3<br>R; 5-CCTACAGATCAGCCTCACCTAG-3 |
| **17** | F; 5-TGTGTCTTTGTGCAGCCATC-3<br>R; 5-TCAGGAACCGTCAGAAAAGC-3 |
| **18** | F; 5-TGAAGTTGGGGCATGAGACT-3<br>R; 5-CAAGCGACTGTGTGACTCTC-3 |
| **19** | F; 5-GCACCACCACACTCAGCT-3<br>R; 5-GTTGGGTGAGGTGGTGCA-3 |
| **20** | F; 5-TAAGCAATCCTCCCACGTCC-3<br>R; 5-GCTGGCATCATCTCTCTGTG-3 |
| **21** | F; 5-AATACACATGTGAGCCTGGC-3<br>R; 5-CATACCATTTTCCCGCCTCAG-3 |
| **22** | F; 5-GGCCTCCGTTTTCTCAAGTG-3 |

| | |
|---|---|
| | R; 5-TAGGGCTAGCAATGGACAGG-3 |
| **23** | F; 5-TGGCAATTTGACTCCAGAATCC-3<br>R; 5-GAAGGAGGTGGGGCATAGAG-3 |
| **24** | F; 5-ACCTGGGGATGGGGAGATAA-3<br>R; 5-TCACTATCTGCCTGCAATGC-3 |
| **25** | F; 5-GCTGAGACTACTGGCTAGCA-3<br>R; 5-CCGAAGCCTTGTCTCTACCA-3 |
| **26** | F; 5-GCCCCACCAACCTATGAAGA-3<br>R; 5-GATTGCAACATCCCTCTGGC-3 |
| **27** | F; 5-TAGTGATGGTGGGAAGCAGG-3<br>R; 5-TTTGGAGGCAGAAATTGGGC-3 |
| **28** | F; 5-GTGATCTAGGGACACGGTGT-3<br>R; 5-TTTCTTGCAGAGTGTAGGCG-3 |
| **29** | F; 5-TTCGACAGCCTTGAAAGTGG-3<br>R; 5-TCCCGGGTTCAAGCAGTT-3 |
| **30** | F; 5-GTGGCGTGCACCTGTAATC-3<br>R; 5-GCACTGCAGCCTAGATCTCC-3 |
| **31** | F; 5-GGAGATCTAGGCTGCAGTGC-3<br>R; 5-ACCCTGGCCTGGCTTTCAA-3 |
| **32** | F; 5-CTCGATCTCCTGACCTGGTG-3<br>R; 5-GCCCCTTCCCCAGAGTAAAG-3 |
| **33** | F; 5-GAAATTTCCCACACCCCAGC-3<br>R; 5-ACATAAGAGCCAGGCACAGT-3 |
| **34** | F; 5-AACCTCCACCACCTGAGTTC-3<br>R; 5-CCACATTTCCTTGACCCATTCA-3 |
| **35** | F; 5-TCCTGTCTTCTCACTCAGCA-3<br>R; 5-TAGAACGGATGGGGTCTTGC-3 |
| **36** | F; 5-GGCCTACAAACGGGACACTA-3<br>R; 5-GCTGGAGTACGATGGTGTGA-3 |
| **37** | F; 5-GATGATCGTGCAACCTTCCC-3<br>R; 5-TTCAGAGGCAGCACACAAAC-3 |
| **38** | F; 5-CCTCAGTTCCCATCCTCCAG-3<br>R; 5-CTCAGTGAGGGGCAAAGAGG-3 |
| **39** | F; 5-TGGCGTGGTTTAGGTAGTGT-3<br>R; 5-CAACCTCATCCTCCCCTCTG-3 |
| Barcode fragment | ACGTCTAAGGCCGCGACTCTAGAGTGTATCTTATCATGAGATCTGTGAAGCTTGTG**NNNNNNNNNNNNNNNNNNNN**TATAAGAGATCGGAAGAGCACACGTCTGAACTCCAGTCAATTCAACGTCTGCCATCTGCCGGCAAGTTGGACGCCCCGAGATTCTCATTAAGGCCAAGAACGGCCGCTTCGAGCAGACATGATA<br>(with RE cut sites underlined, in order XbaI, BglII, HindIII and EagI) |
| Tag-seq | PCR1-F; 5-CACGACGCTCTTCCGATCTTCATGAGATCTGTGAAGCTTGTG-3<br>PCR1-R (universal primer); 5-CAAGCAGAAGACGGCATACGAGATAACTTGCCGTGACTGGAGTTCAGACGTGTGCTCTTCCGATCT-3<br>PCR2; use Illumina 6bp index primers and universal primer |
